# Evolutionary diversity of cell-type-specific expression and stress response in Brassicaceae roots

**DOI:** 10.1101/2024.06.21.599952

**Authors:** Guannan Wang, Kook Hui Ryu, Andrea Dinneny, Jiyoung Lee, Dong-Ha Oh, Prashanth Ramachandran, Marina Oliva, Ryan Lister, José R. Dinneny, John Schiefelbein, Maheshi Dassanayake

**Author notes:** These authors contributed equally to this work (Guannan Wang, Kook Hui Ryu, Andrea Dinneny). Correspondence (J.R.D.); (J.S.); (M.D.).

## Abstract

Plants are composed of diverse cell types that facilitate adaptations to the environment, yet cross-species comparisons of such response programs at single-cell resolution remain scarce. To explore this diversity, here we profiled >200,000 root cells from five Brassicaceae species, including stress-sensitive species (*Arabidopsis thaliana, Sisymbrium irio*), extremophytes (*Eutrema salsugineum, Schrenkiella parvula*), and the polyploid crop *Camelina sativa* under control, NaCl, and abscisic acid (ABA) treatments. We developed a computational pipeline to characterize the conservation and divergence of cell-type gene expression across the Brassicaceae, revealing that approximately half of previously defined Arabidopsis cell-type markers fail to maintain conserved expression in one or more non-Arabidopsis species. Therefore, we curated a refined set of pan-Brassicaceae markers and identified orthologs whose expression profiles have diverged across lineages. Using *in situ* hybridization, we mapped distinct cortex subpopulations to specific cortical layers across species, and found flavonoid biosynthesis programs preferentially localized to the inner cortex layer, linking cortex-layer specification to metabolic specialization. Cell-type contributions to stress responses differed among species and across treatments, with lineage-specific losses of responsiveness occurring less frequently but evolutionarily more favored than lineage-specific gains. In *C. sativa*, sub-genomes contributed equally to stress responses, and homeologs with divergent responses typically lacked other signatures of functional divergence. Together, this work establishes a foundational root single-cell atlas and an analytical framework for multi-species comparative transcriptomics, providing insights into how stress responses diversify across cell types, stress-sensitive to stress-adapted species, and in crop lineages.

## INTRODUCTION

Environmental stress responses are complex traits that require spatiotemporal coordination of gene networks functioning in diverse assemblages of cell types. Plant roots, the primary interface with soil, are a mixture of functionally and transcriptionally diverse cell types, each perceiving and responding to environmental stresses in a development-dependent way^1–3^. Investigations of plant environmental responses at the cell-type level have been limited to model plants like *Arabidopsis thaliana*^4,5^, and thus provide only partial insights into the diverse physiological strategies employed by plants. This limitation becomes more evident when considering the rewired and novel stress tolerance mechanisms employed by naturally stress-adapted species (extremophytes) that exhibit a remarkable ability to optimize growth and survival under challenging environments^6^. However, such comparisons are complicated in distantly related species, where limited one-to-one orthology constrains evolutionary inference at the level of individual genes. Therefore, establishment of a comparative approach within a model clade of related species not only enhances our understanding of the diversity of stress adaptations but also uncovers evolutionary innovations that confer stress resilience.

In this study, we have selected ecologically diverse species within the Brassicaceae family that include the extremophyte models, *Eutrema salsugineum* and *Schrenkiella parvula,* the more stress-sensitive models, *Arabidopsis thaliana* (hereinafter referred to as Arabidopsis) and *Sisymbrium irio* (a close relative of the extremophyte models that exhibits better salt stress tolerance compared to Arabidopsis^7^), and the polyploid oil crop, *Camelina sativa*. Previous studies reported divergent physiological responses among these species not only to high salinity, but also to the stress hormone, abscisic acid (ABA)^8–10^. Moreover, it has been demonstrated that stress conditions can elicit distinct responses across different root cell types, and these cell-type specific responses can vary between species^1,2,11,12^. However, how individual cell types contribute to the diversity of stress responses remains largely unknown. We generated primary root transcriptome atlases for the selected species at single-cell resolution under stress-neutral as well as ABA- and salt-treated conditions, totaling >200,000 cells across five species (>174,000 cells across four diploid species plus >27,000 cells in *C. sativa*). This experimental approach allowed us to compare recently diverged lineages and to examine responses within a defined phylogenetic group with a shared common ancestor ∼33 Myo^13^. We developed a comparative framework and methodology for cross-species cell-type annotation and single-cell evolutionary transcriptomics that can be readily transferred to other multi-species comparisons. Using this framework, we (i) quantify the limited conservation of canonical Arabidopsis cell-type markers across the Brassicaeae and define a refined set of conserved markers; (ii) identify significant changes in ortholog expression profile on different phylogenetic branches; (iii) spatially map cortex subpopulations to defined cortex layers across species and provide evidence for cortex functional specialization; (iv) resolve how distinct cell types contribute to salt versus ABA responses across lineages and identify patterns of evolutionary turnover in responsiveness; and (v) evaluate the extent to which subgenomes and homeologs diversify stress responses in the polyploid crop *C. sativa*. Together, this work establishes a foundational root cell atlas to explore cell-specific, species-specific, or stress-dependent mechanisms across ecologically diverse but phylogenetically close Brassicaceae model plants.

## RESULTS

### The expression of nearly half of the detected Arabidopsis cell type marker genes is not conserved across species

We generated 36,000 to 50,000 root cell transcriptomes for four diploid Brassicaceae species under control (stress-neutral), 5 µM ABA, and 100 mM NaCl treatments (n = 2 for each condition) using the 10X Genomics scRNA-seq platform to provide over 174,000 root cell transcriptomes from the four diploid species (Fig. 1 and Supplementary Fig. 1; Supplementary Movie 1). To investigate the impact of genome duplication on stress adaptation, we additionally generated a comparable dataset of over 27,000 root cell transcriptomes for an allohexaploid Brassicaceae species using an identical experimental design. The percentage of orthologs between species generally decreases with increasing species divergence (Supplementary Fig. 2), which limits comparative studies investigating large ortholog gene networks. We therefore selected more closely-related species (diverged max. ∼33Mya) of Arabidopsis^13^ from the Brassciceae family for our comparison, including the diploid *E. salsugineum*, *S. irio*, *S. parvula*, and hexaploid *C. sativa*. This approach allowed us to identify 15,198 unambiguous 1-to-1 orthologous genes across the diploid species and the Arabidopsis orthologs for over 90% of genes in *C. sativa* (details in Methods). Given the complex 1-to-3 orthology relationship inherent to hexaploidy, we intentionally excluded *C. sativa* from 1-to-1 ortholog based cross-species comparisons (Figs. 1-6) and analyzed this species separately (Fig. 7) to preserve sub-genome resolution. Regardless of the species and conditions, we consistently generated an average of 35,077 reads for each cell, resulting in the detection of a median of 1,959 unique transcripts and 1,077 genes expressed per cell (Supplementary Data 1 and Supplementary Fig. 3). We used ABA as a more generic abiotic stress stimulus and NaCl as a more specific stimulus. We selected treatment strength and duration based on our prior work showing that ABA (μM, 3h) and NaCl (100-150 mM, 24h) trigger strong root transcriptomic responses and physiological changes across species, including extremophytes, without causing immediate tissue necrosis in sensitive plants like Arabidopsis^9,10,12,14^. To facilitate the utility of this multi-species dataset as a community resource, we generated an accompanying interactive web browser to enable exploration of this atlas at https://plantbiology.shinyapps.io/brassicaceaeatlas. Preprocessed data matrices partitioned by cell types are provided in the searchable browser for convenient reanalysis, as well.

**Fig. 1.**
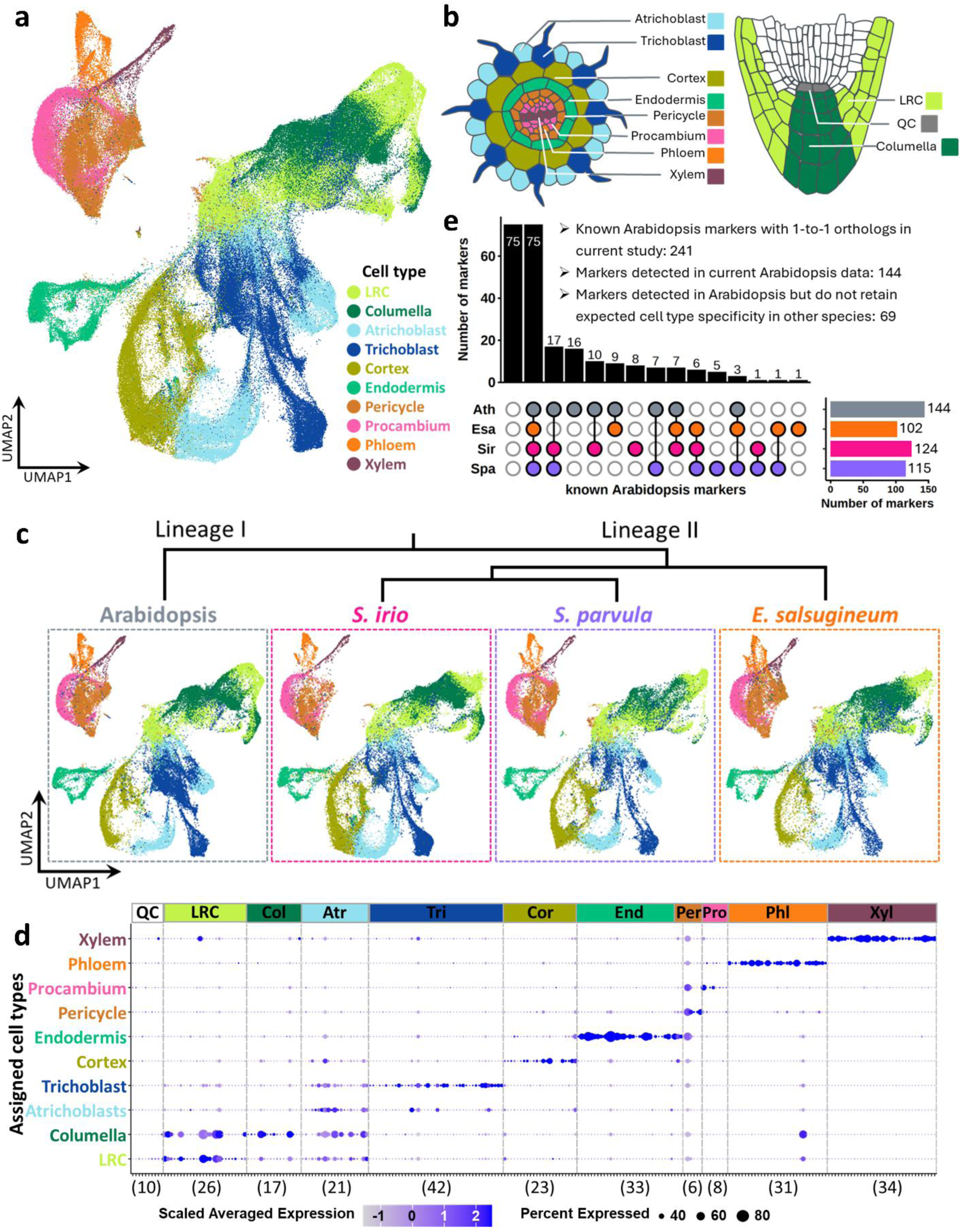
Single cell RNA profiling of root cells across 4 diploid Brassicaceae species: Arabidopsis, *Sisymbrium irio*, *Schrenkiella parvula*, and *Eutrema salsugineum*. a Integrated UMAP view of root atlas (174,820 cells) from 4 diploid Brassicaceae species representing 10 major root cell types. The integrated atlas included control, 5 μM ABA- and 100 mM NaCl-treated samples. b Schematic illustration of major Arabidopsis root cell types annotated in current study. Illustration was adapted from the Plant Illustrations repository^88^. c Root cell atlas for individual species on the integrated UMAP space. The atlas for each species included control, 5 μM ABA- and 100 mM NaCl-treated samples. d Preferential expression of known cell-type markers based on cells from all species under control condition. See Supplementary Fig. 8 for detailed preferential marker gene expression in each of the four species. See Supplementary Data 2 for the list of markers used. Dot size represents the percentage of cells in which each gene is expressed. Dot color intensity indicates the normalized expression of each gene in each cell type. Numbers in parenthesis indicate the number of marker genes per cell type. QC, Quiescent center; LRC, Lateral Root Cap; Col, Columella; Atr, Atrichoblast; Tri, Trichoblast; Cor, Cortex; End, Endodermis; Per, Pericycle; Pro, Procambium; Phl, Phloem; Xyl, Xylem. e Number of known Arabidopsis markers that were detected in each species under control condition in the current study. Ath, Arabidopsis; Esa, *Eutrema salsugineum*; Spa, *Schrenkiella parvula*; Sir, *Sisymbrium irio*.

Root cell-type marker genes have been established for Arabidopsis^15–21^, though their applicability as markers for root cell types in related species has been untested. We developed a two-step strategy to annotate cell types for Arabidopsis and non-Arabidopsis cells in our dataset (Supplementary Fig. 4; Methods). We used Arabidopsis cells from the control condition, and annotated these individual cells based on the collective outcome of three independent methods: correlation-based annotation (where individual cells were annotated based on their transcriptomic similarity to reference cell type transcriptomes), marker-based annotation (where individual cells were annotated based on the preferential expression of known cell type markers), and integration-based annotation (where individual cells were annotated based on their mapping to reference single-cell datasets) (Supplementary Fig. 4 - Step 1). This provided us with a reference atlas to identify cell types for the remaining species. We employed three prediction algorithms (TransferData in Seurat^22^, Symphony^23^, and scArches^24^) to transfer Arabidopsis annotations to individual cells in the other species based on 15,198 unambiguous 1-to-1 orthologous genes identified (Supplementary Fig. 4 - Step 2, details in Methods). We additionally annotated our dataset with differential developmental zones using a similar approach. This resulted in the identification of ten major root cell types and three developmental zones for all four species from all conditions in our dataset (visualized on the integrated UMAP space, Figs. 1a-1c, Supplementary Fig. 5 and 6). We did not detect a recognizable cell population for quiescent center (QC) cells in our dataset (Supplementary Fig. 7), likely due to under-sampling of this rare cell type. Our overall cell-type identification indicates that these diverse Brassicaceae species do not differ in the presence of major root cell types.

The expression conservation of known Arabidopsis cell-type marker genes was examined in the other target species (Fig. 1d and Supplementary Fig. 8). Out of 461 previously established Arabidopsis cell-type markers, we found 1-to-1 orthologs for 254 of them, 241 of which were representative marker genes for the ten major root cell types described above (Supplementary Data 2). Only 144 of these Arabidopsis markers had expression levels above the threshold of detection in our Arabidopsis data (Fig. 1e). Among these Arabidopsis markers, while many retained robust expression and cell-type specificity across all species, 48% (69/144) of markers lost expression specificity in one or more species (Figs. 1d and 1e, Supplementary Fig. 8 and 9). Among the non-conserved Arabidopsis markers, lost conservation can be attributed in most cases to low detectable expression in the expected target cell types in one or more non-Arabidopsis species (Supplementary Fig. 9b). The lowest rate of conservation for Arabidopsis cell-type markers was observed in *E. salsugineum* (Fig. 1e and Supplementary Fig. 8) despite the data quality for this species being comparable to that of other species (Supplementary Data 1). The cell-type markers for vascular tissues were generally highly conserved across species, while markers from outer cell layers of the root, especially the root cap and epidermal cells, were least conserved (Supplementary Fig. 8). Similar patterns of conservation were also observed when comparing tissue-specific gene expression programs between environmental conditions in Arabidopsis^1^.

To obtain a panel of pan-Brassicaceae cell type markers, we identified orthologous genes exhibiting highly conserved expression for each of the ten cell types (Supplementary Fig. 10; Supplementary Data 3). To test the accuracy of this panel, we optimized a whole-mount hybridization chain reaction (HCR)-based *in situ* hybridization method^25^ for *in vivo* mapping of the selected markers. *In situ* expression patterns matched the expected cell-type specificity based on the scRNA-seq data (Fig. 2 and Supplementary Fig. 11). This provided independent support for the cell-type assignments in the target species and shows that these orthologous genes represent broadly conserved cell-type markers.

**Fig. 2.**
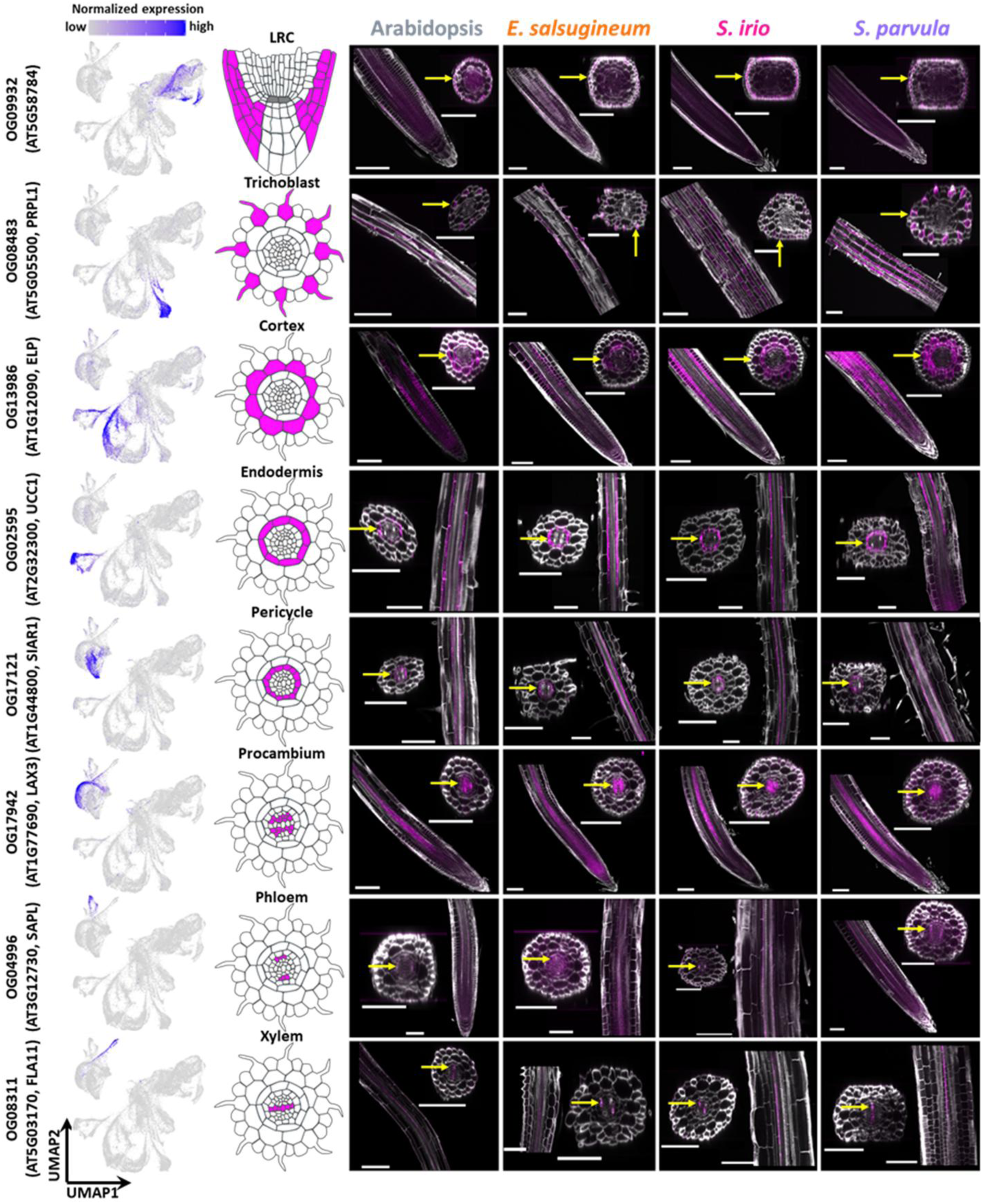
Whole-mount *in situ* HCR of cell type markers that are conserved between the 4 Brassicaceae species under control condition. Left panel, the expression of selected cell type markers in our integrated atlas from the control samples (for visualization purpose only, the markers were identified separately for each species); middle panel, schematic representation of Arabidopsis root cell types with the target cell type highlighted in magenta; right panel, whole-mount *in situ* HCR images of selected cell type markers across species. Note that *S. irio* and *S. parvula* have two layers of cortex. Arrows in yellow mark the cell type where the marker expression is expected. PRPL1, proline-rich protein-like 1; ELP, extensin-like protein; UCC1, uclacyanin 1; SIAR1, siliques are red 1; LAX3, like aux1 3; SAPL, Sister of APL; FLA11, FASCICLIN-like arabinogalactan-protein 11. Scale bars, 100 μm.

### Altered cell-type expression specificity drives lineage-specific molecular phenotypes

We observed that orthologs that retained similar cell type specificity across all four species accounted for only a small proportion (∼ 3% of the 1-to-1 orthologs with all pairwise Pearson correlations greater than 0.9) of the entire ortholog sets (Fig. 3a and Supplementary Fig. 12). The overall correlations of cell-type expression among orthologs were modest even between each pair of species (Fig. 3a). To determine the patterns of divergence in cell-type expression between species under stress-neutral conditions, we compared 1-to-1 ortholog expression across the four species using a phylogenetic framework, and detected 350 orthologs showing diverged expression profiles in one phylogenetic branch compared to others or between extremophytes and non-extremophytes (Fig. 3b; Supplementary Data 4). For example, there were 78 orthologs whose expression pattern in Arabidopsis (lineage I) differed from that of orthologs in the other three (lineage-II) species. We also found 43 orthologs whose cell-type expression differentiate extremophyte and non-extremophyte species (Fig. 3b), independent of their phylogenetic relationships.

**Fig. 3.**
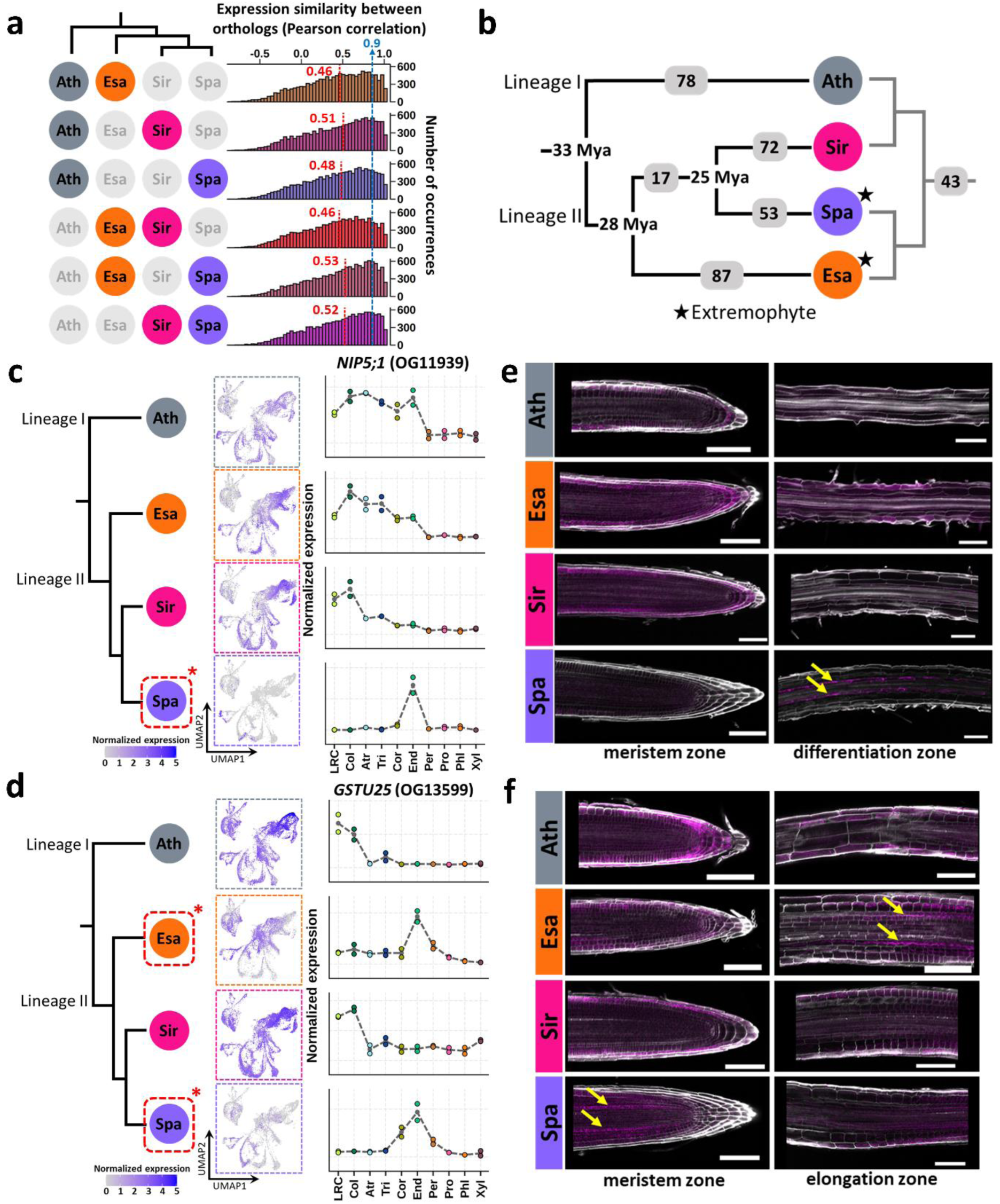
Evolution of gene expression profiles across cell types under stress-neutral conditions. a Distribution of Pearson correlation coefficients for the cell type expression of 15,027 1-to-1 orthologs between each pair of species under control condition. The cell type expression was calculated by averaging the normalized expression level for each gene across all the single cells within each cell type, see Methods for details. The line and text in red indicate the median. b The numbers of orthologs showing diverged expression profiles between species groups under control condition. Orthologs whose cell type expression in one species group is not conserved in another species group were considered as orthologs with diverged expression profiles, see details in Methods. c-d Cell type expression of selected examples with diverged expression across species from control samples, *NIP5;1* (c) and *GSTU25* (d). Asterisks indicate where the diverged expression was detected. Left, expected expression of the selected examples based on scRNA-seq data visualized on UMAP space. Right, pseudo-bulk cell type expression of the selected examples. LRC, Lateral Root Cap; Col, Columella; Atr, Atrichoblast; Tri, Trichoblast; Cor, Cortex; End, Endodermis; Per, Pericycle; Pro, Procambium; Phl, Phloem; Xyl, Xylem. e-f Whole-mount *in situ* HCR of *NIP5;1* (e) and *GSTU25* (f) across species under control condition. Arrows in yellow mark the cell type where the diverged expression is expected. Scale bar, 100 μm. Ath, Arabidopsis; Esa, *Eutrema salsugineum*; Spa, *Schrenkiella parvula*; Sir, *Sisymbrium irio*.

We used HCR to examine two of top-ranked diverged cell-type expression profile changes associated with the extremophytes: restricted expression of *NOD26-like intrinsic protein 5;1* (*NIP5;1*) in *S. parvula*, and extremophyte-shared expression of *glutathione S-transferase U25* (*GSTU25*) (Figs. 3c and 3d). As both *NIP5;1* and *GSTU25* are members of larger gene families, to ensure the detected expression changes across species is not due to the misassignment of orthologs from the same gene family, we identified all *NIP*s and *GSTU*s among these species and extracted their protein sequences for phylogeny construction. As expected, *NIP5;1* and *GSTU25* orthologs used in our analyses formed their own branches separated from all other *NIP*s and *GSTU*s, respectively, confirming the assignment of orthologs (Supplementary Fig. 13). In the *GSTU25* branch, we noticed the presence of an additional gene from *S. parvula*, *Sp1g15325*, which is a non-reciprocal-best-match for *GSTU25*s from other species (Supplementary Fig. 13). It exhibited a *SpGSTU25*-like cell type expression profile that was shared with *E. salsugineum* but not with the non-extremophytes (Supplementary Fig. 14).

Both scRNA-seq data and HCR results showed that the gene *NIP5;1*, which encodes a plasma membrane-localized aquaporin, is predominantly expressed in the outer cell layers, from epidermal to endodermal cells, in Arabidopsis, *S. irio*, and *E. salsugineum,* while its expression is restricted to differentiation zone endodermal cells in *S. parvula* (Figs. 3c and 3e). Arabidopsis AtNIP5;1 serves as a channel responsible for transporting boron into roots^26^. The restricted expression of *NIP5;1* in *S. parvula* may reduce boron uptake into the vascular stream. *S. parvula* is native to boron-rich soils in Anatolia, Turkey and can tolerate high boron levels^14^ while the other three species are not associated with exceptionally high boron habitats. Previous studies have demonstrated that *S. parvula* achieves boron toxicity tolerance mainly by maintaining low intracellular boron levels^27^, which is consistent with the more restricted expression of *NIP5;1* in this species.

Our scRNA-seq data indicate that *GSTU25*, encoding a plant-specific Tau class glutathione S-transferase, is preferentially expressed in the root cap of non-extremophytes (Fig. 3d). In the extremophytes, its expression becomes enriched in the endodermal cells of the meristematic and early elongation zones (Fig. 3d). HCR experiments confirmed the absence of endodermal expression of *GSTU25* in the non-extremophytes (Fig. 3f). However, our HCR observations did not fully align with the scRNA-seq data, particularly for *E. salsugineum* where *GSTU25* expression was also detected in epidermal and cortex cells (Figs. 3d and 3f). This discrepancy may be due to the technical differences between the two complementary RNA detection methods. Specifically, because of its signal amplification mechanism and better cell resolution, HCR was able to detect the expression of *GSTU25* in the cell population that might have been lost or under-sampled in scRNA-seq. Glutathione S-transferases protect cells from oxidative damage by utilizing glutathione to scavenge reactive oxygen species (ROS)^28^. *GSTU25* has been reported to be induced under oxidative, salt, and heat stresses^29,30^. The convergent endodermal expression of *GSTU25* in the two extremophytes might suggest a role of *GSTU25* in ROS scavenging in endodermal cells, which may contribute to salt tolerance in the two extremophytes.

### Transcriptomic specialization of cortex cell subpopulations supports lineage-specific root anatomy

Anatomical analysis of *S. irio* and *S. parvula* reveals that both species possess two cortical cell layers, whereas Arabidopsis and *E. salsugineum* possess a single cortical cell layer at the early developmental stage of seedlings (Supplementary Fig. 15a). Cortex cells from the single cortical layer in Arabidopsis have been reported to contain two transcriptionally distinct subpopulations^31^. Agreeing with this observation, UMAP-based analysis of cortex cells in our dataset identified two major cortex subpopulations for Arabidopsis and *E. salsugineum*.

Contrastingly, three cortex subpopulations were found in *S. parvula* and *S. irio* (Supplementary Fig. 15b). We hypothesized that the extra cortex cell population identified in *S. irio* and *S, parvula* is derived from the additional cortex layer. If supported, we would expect that the extra cortex cell population should be absent or underrepresented in Arabidopsis and *E. salsugineum*. To test this, we selected the cortex subpopulations from *S. irio*, which had the highest separation of clusters, as the reference and categorized the three subpopulations from this species into “left”, “middle”, and “right” subgroups (Fig. 4a). Cortex cells from the other three species were mapped to the reference *S. irio* cortex subpopulations based on 1-to-1 orthologs (Fig. 4b). While some representation of the three cortex subpopulations could be detected in all species (Figs. 4b and 4c), the middle population was highly depleted in Arabidopsis and *E. salsugineum*, but not in *S. parvula* (Fig. 4c). To test whether the “middle” cortex subpopulation represented the extra cortex layer found in *S. irio* and *S. parvula*, we identified marker genes that were preferentially expressed in the “middle” population (Supplementary Fig. 16a; Supplementary Data 5) and examined their spatial expression in roots using HCR. The *in situ* hybridization results showed that *Si-s1384-01650*, which was specifically expressed in the middle population (Fig. 4d, upper panel), distinctly marked the inner cortex layer in *S. irio* (Fig. 4d, lower panel), in support of our hypothesis. Similarly, the ortholog (in the ortholog group OG14187) of *Si-s1384-01650* showed expression predominantly in the inner cortex layer of *S. parvul*a, while in *E. salsugineum* and Arabidopsis, expression was detected in all cells of the single cortex layer (Fig. 4e). These results suggest that the middle subpopulation is derived from the inner cortex layer, while the other two populations originate from the outer layer. To characterize the transcriptomic signatures of the inner and outer cortex layers, we identified genes that were preferentially expressed in either of the layers (Supplementary Fig. 17). Both layers highly expressed genes involved in water transport and environmental sensing (Supplementary Fig. 17). Genes preferentially expressed in the inner cortex are mostly associated with secondary metabolism (particularly flavonoid biosynthesis), cell-to-cell communication via plasmodesmata, and jasmonic acid and ABA signaling. In contrast, the outer cortex prioritized hypoxia sensing, thermal stress response and gibberellin signaling pathway gene functions (Supplementary Fig. 17).

**Fig. 4.**
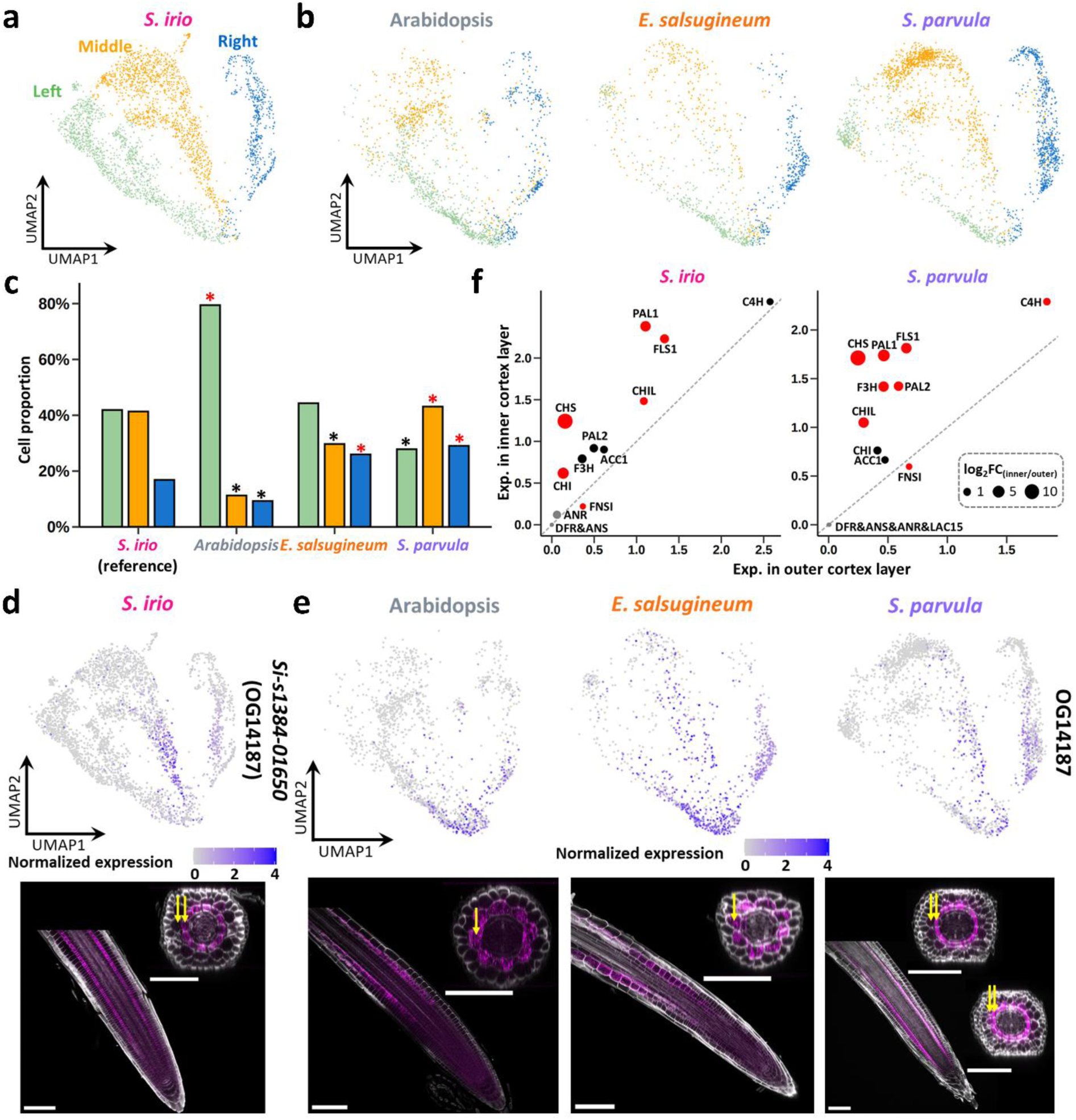
Comparison of cortex subpopulations among species from control samples. a UMAP representation of three cortex subpopulations in *S. irio*. b Projection of cortex cells from Arabidopsis, *E. salsugineum*, and *S. parvula* onto the three cortex subpopulations in *S. irio*. c Proportion of the three cortex subpopulations in the four Brassicaceae species. Asterisks represent significance determined by comparing each species to *S. irio* using hypergeometric test at *p*-value ≤ 0.01 with red asterisks indicating over-representation and black asterisks indicating under-representation. d Normalized expression of *Si-s1384-01650* (OG14187) on UMAP space (top) and its whole-mount *in situ* HCR (bottom) in *S. irio*. Arrows in yellow mark the cortex layers. Scale bars, 100 μm. e Normalized expression of *Si-s1384-01650* orthologs (OG14187) on UMAP space (top) and their whole-mount *in situ* HCR (bottom) in Arabidopsis, *E. salsugineum*, and *S. parvula*. Arrows in yellow mark the cortex layers. Scale bars, 100 μm. f Normalized expression of genes involved in flavonoid biosynthesis in inner and outer cortex layers in *S. irio* (left) and *S. parvula* (right). Red dots indicate genes that were significantly differentially expressed (adjusted *p*-value ≤ 0.01 in FindMarkers) between the two cortex layers and size of the dots corresponds to the log_2_FC. PAL1/2, phenylalanine ammonia-lyase1/2; C4H, cinnamic acid 4-hydroxylase; 4CL3, 4-coumaric acid:CoA ligase; ACC1, acetyl-CoA carboxylase; CHS, chalcone synthase; CHI, chalcone isomerase; CHIL, chalcone isomerase-like; F3H, flavanone 3-hydroxylase; F3′H, flavonoid 3′-hydroxylase; FLS1/3, flavonol synthase1/3; DFR, dihydroflavonol reductase; ANS, anthocyanin synthase; ANR, anthocyanin reductase; LAC15, laccase 15; FNSI, flavone synthase I.

To further investigate the transcriptomic differences between the inner and outer cortex layers in *S. irio* under control conditions, we compared the transcriptomes between the inferred inner (middle subpopulation) and outer cortex layers (left and right subpopulations, Supplementary Fig. 16). This identified several cellular processes (among 155 genes that were differentially expressed between the two layers) that were differently partitioned between the two layers, including flavonoid biosynthesis (Supplementary Fig. 16) which has been previously shown to be preferentially localized to cortex cells^32^. We detected a strong expression bias towards the inner layer in *S. irio* for about 69% of the genes involved in flavonoid biosynthesis, including CHALCONE SYNTHASE (CHS) (Fig. 4f, left panel), which catalyzes the first committed step in this biosynthetic pathway. Intriguingly, this inner cortex layer-bias is also conserved in *S. parvula* cortex cells (Fig. 4f, right panel), suggesting that this transcriptional bias preceded the divergence of this subclade of species. Similar biases in flavonoid biosynthetic gene expression were not observed in other species with more than one layer of cortex (Supplementary Fig. 18), including rice^33^ and Medicago^34^, suggesting that the partitioning of flavonoid biosynthesis is unlikely to be a conserved signature of cortex functional diversification. Flavonoids are known to function as antioxidants, modulators of reactive oxygen species and hormone transport and signaling, and have been implicated in plant tolerance to environmental stresses^35,36^. The spatial restriction of flavonoid biosynthesis to the inner cortex therefore suggests a lineage-specific partitioning of a growth and stress-relevant metabolic pathway within the root. Together these data demonstrate transcriptional divergence in the cortex cell populations, corresponding with the increased anatomical complexity observed in *S. irio* and *S. parvula* roots. Our results suggest that the presence of two cortex layers may have enabled the inner layer to specialize in flavonoid metabolism.

### ABA and NaCl induce distinct cell-type-specific responses

To obtain an overview of cell-type transcriptional diversity across species and in response to stress inducers, we performed a principal component analysis (PCA) using the 15,198 1-to-1 orthologous genes from pseudo-bulk cell-type transcriptomes (Fig. 5a). Although the transcriptomes of all cell types clustered according to their respective species, those originating from the two extremophytes formed a distinct subgroup that was separate from the transcriptomes of other non-extremophytes. Since *S. parvula* and *S. irio* are more closely related phylogenetically than are the two extremophytes, the observed similarities between extremophytes suggests that these shared features arose independently of phylogenetic relatedness. Within each species, outer root cell layers (root cap, epidermal, cortex, and endodermal cells) formed a distinct cluster from the inner cell layers (pericycle, procambium, phloem, and xylem cells), regardless of treatments. We next tested if these observed differences could be a consequence of protoplasting using published single-nucleus RNA-seq for Arabidopsis^37^ where cells were not subjected to a protoplasting step. The two groups, inner and outer cell types, remained as two distinct groups (Supplementary Fig. 19). Therefore, the transcriptomic divergence between the inner and outer cell layers suggests that these two zones of the root operate within distinct signaling contexts and exhibit a more evolutionary conserved and robust developmental trajectory across species.

**Fig. 5.**
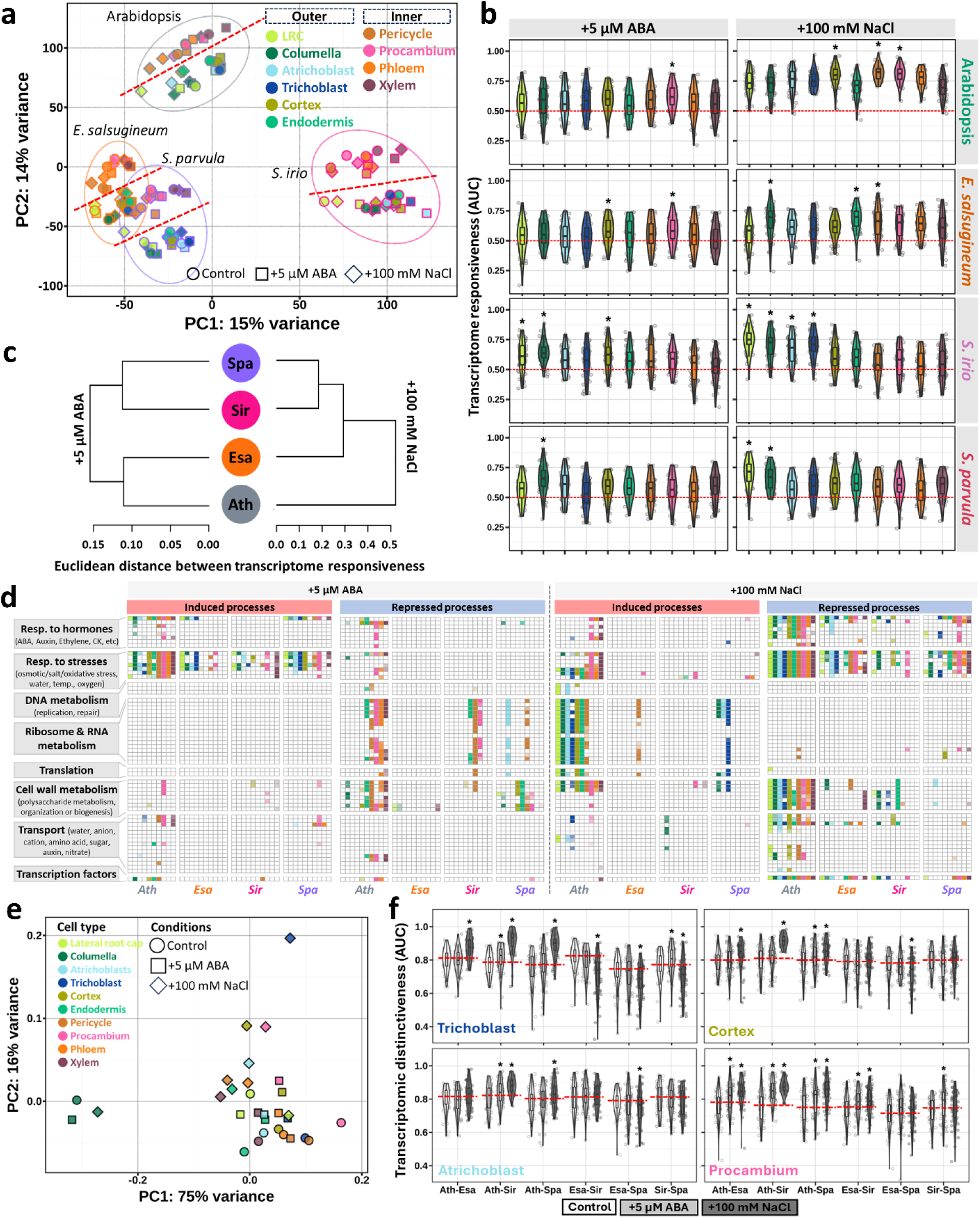
Divergent cell type transcriptomic response to ABA and NaCl. a Principal component (PC) analysis of pseudo-bulk cell-type transcriptomes from control, 5 μM ABA- and 100 mM NaCl-treated samples. Each symbol represents a pseudo-bulk cell-type transcriptome. The color and the shape of each symbol correspond to the cell type and the condition, respectively. Samples from each species are encircled by a line (based on 95% confidence level) and labeled. The red dashed lines separate cell type transcriptome subgroups within each species. b Comparison of transcriptome responsiveness as averaged Augur area under the curve (AUC) scores across cell types in response to ABA and NaCl. AUC here quantifies the separability between control and treatment samples for each cell type. AUC = 0.5 indicates random separability; AUC = 1.0 indicates perfect separability. Asterisks indicate significance after Wilcoxon testing (n = 50) at adjusted *p*-value cutoff of 0.05. c Hierarchical clustering of species based on Euclidean distance between their cell type transcriptome responses in (b). d Summary of GO functional enrichment among DEGs in response to ABA (left) and NaCl (right) for each cell type in each species. The detailed list of GO terms is present in Supplementary Data 7. The colors of the tiles represent different cell types. The shade represents the *p*-value assigned by the enrichment test at adjusted *p*-value < 0.05 with darker shades indicating smaller *p*-values. e Principal component analysis of transcriptomic divergence (as AUC scores) between species for each cell type. AUC here quantifies the separability between species for each cell type and was calculated for each pair of species under control, ABA and NaCl treatments separately. Each symbol represents the transcriptomic divergence between six possible species pairs under one condition. The color and the shape of each symbol correspond to the cell type and the condition, respectively. f Transcriptomic distinctiveness (as AUC scores) between species pairs under different conditions. AUC here quantifies the separability between species for each cell type. AUC = 0.5 indicates random separability; AUC = 1.0 indicates perfect separability. Asterisks indicate significance after comparing to control (n = 50) with Student’s t-test at *p*-value cutoff of 0.05. The red dash lines mark the medians the transcriptomic distinctiveness under control condition. Ath, Arabidopsis; Esa, *Eutrema salsugineum*; Sir, *Sisymbrium irio;* Spa, *Schrenkiella parvula*.

To explore which genes, processes, or cell states drive the overall distinct stress responses between inner and outer cell layers, we next identified differentially expressed genes (DEGs) on a per cell type, species, and treatment basis (details in Methods; Supplementary Data 6). To determine the stress response in each developmental zone, we applied the same approach to identify DEGs in all zones (Supplementary Data 6). We selected three ortholog groups exhibiting species-specific transcriptional responses to explore using HCR. For example, in response to ABA, members of orthologous group OG04736 (AT3G15670), which encode a late embryogenesis abundant protein (LEA), showed a ubiquitous increase in expression across tissues, which was largely captured by HCR (Supplementary Fig. 20). HCR also confirmed expression changes in another ortholog group, OG03053 (AT2G37870), which exhibited the highest expression in stele cells compared to all other cell types and was further induced by ABA (Supplementary Fig. 21). In another example, the expression of the *tonoplast intrinsic protein 2;3* (*TIP2;3*; AT5G47450; OG10873) was consistently reduced in multiple cell types across species with exposure to NaCl. The HCR signal for *TIP2;3*s displayed a similar trend in expression across all species (Supplementary Fig. 22). These data support the overall trends observed with scRNA-seq and further attest to the fidelity of the cell-type transcriptional profiles generated through single cell protoplasting and scRNA-seq analysis.

Previous studies have identified Arabidopsis as the most sensitive of the four species, exhibiting the strongest transcriptional response to ABA- and NaCl-induced stress^9,10^. Our results support this finding, showing Arabidopsis has the highest number of differentially expressed genes (DEGs), ranging from 2 to 10 times more than the other species (Supplementary Fig. 23). The number of DEGs also differed among different developmental zones across species with a substantial portion of them being specific to individual zones (Supplementary Fig. 24). To compare the responses to ABA and NaCl treatments, we performed a direct comparison of stress responses by identifying genes that showed shared or distinct responses to ABA and NaCl (Supplementary Fig. 25). This quantitative assessment revealed that genes responsive to both stimuli typically accounted for less than 50% of the total responsive genes, with a notable exception for Arabidopsis where up to 78% of the ABA-responsive genes were shared with NaCl stress response (Supplementary Fig. 25). While these results agree with previously reported overlap between these two treatments^9^, it also suggests that the majority of the transcriptomic remodeling is indeed stress-specific, and salt stress activates not only ABA-mediated responses but also a large additional set of ABA-independent pathways.

Quantification of cell type responsiveness using the number of DEGs can be confounded by factors such as the number of cells per cell type and different effect sizes of genes^38^. To assess cell-type response signals isolated from internal noise or other confounding factors, we implemented Augur^38^, a method to rank cell types based on the magnitude of their response in single-cell data. Procambium cells showed the strongest response to ABA in Arabidopsis and *E. salsugineum,* whereas columella cells were the most responsive in *S. irio* and *S. parvula* (Fig. 5b). The species-specific cell-type responses further intensified when plants were treated with NaCl. In Arabidopsis and *E. salsugineum*, inner cell layers were the most responsive cell types, whereas in *S. irio* and *S. parvula* the root cap was most responsive (Fig. 5b). While the two extremophytes’ transcriptomes cluster closer to each other (Fig. 5a), the overall normalized transcriptional responses to stress in each species indicated that *S. parvula* and *S. irio* were more similar, aligning with their closer phylogenetic relationship (Fig. 5c).

We examined the functional processes enriched in DEGs to identify the most predominant cellular processes associated with stress responses (Fig. 5d; Supplementary Data 7). Response to stress was among the most commonly shared processes enriched in DEGs induced by ABA and those repressed by NaCl across most cell types in all four species. RNA metabolism and ribosome biogenesis were found exclusively enriched among repressed genes in response to ABA. Additionally, these processes exhibited the greatest variation among cell-type specific responses across species, being primarily enriched in endodermal and stele cells in Arabidopsis and *S. irio*, but in atrichoblast cells in *S. parvula* (Fig. 5d). On the contrary, these same processes were enriched only among induced DEGs in response to NaCl across species. Moreover, these over-represented functions were primarily found in outer cell types in Arabidopsis, but restricted to only pericycle cells in *E. salsugineum* and epidermal cells in *S. parvula* (Fig. 5d). These results indicate that although stress signaling involves similar biological functions in each species, the key differences lie in the cell-specific patterns of these responses.

The observed divergence in stress responses among species seemed to primarily occur between cell types. Therefore, we computed alterations in transcriptomic distinctiveness between pairs of species for each cell type in response to ABA and NaCl treatments. This allowed us to pinpoint the cell types that exhibited the most pronounced divergence in their responses. We observed overall larger transcriptomic distinctiveness between species in response to 100 mM NaCl compared to that of 5 µM ABA (Fig. 5e, Supplementary Fig. 26). The columella cells exhibited the highest transcriptomic divergence compared to other cell types under stress-neutral conditions (Fig. 5e and Supplementary Fig. 27), consistent with a previous report for grass species^39^. ABA and NaCl treatments did not dramatically alter the high level of columella transcriptomic divergence between species (Supplementary Fig. 27). In contrast, trichoblast cells showed the most pronounced shifts for between-species transcriptomic distinctiveness upon NaCl treatment where transcriptomic divergence increased between Arabidopsis and lineage II species but decreased among lineage II species. Cortex and atrichoblast transcriptomes significantly increased their similarity following NaCl treatment only between the two extremophytes (Fig. 5f, quantified as NaCl-induced reduction in transcriptomic distinctiveness between the extremophytes in Supplementary Fig. 28), indicating convergent salt-induced transcriptomic adjustments in these two cell types for the extremophytes. Procambium showed the most prevalent increases in transcriptomic divergence between species, particularly in response to 5 µM ABA treatment, whereas most other cell types remained transcriptomically similar between species (Fig. 5f, and Supplementary Figs. 27 and 28). Since procambium orchestrates the development of vascular tissues and is sensitive to stress and hormonal signaling, the pronounced divergence observed in procambium suggests that stress-responsive transcriptional programs may be preferentially rewired in tissues coordinating growth and transport rather than being restricted to terminally differentiated cell types.

### Phylogenetic modeling reveals asymmetric evolutionary rewiring of stress responses across species

We aimed to investigate the occurrences of ortholog response shifts between stress-sensitive differential expression (DEGs) and stress-insensitive expression (non-DEGs), examining whether such changes correlated to phylogenetic relationships. For this analysis, we considered all ortholog groups that had at least one DEG responsive to ABA or salt in at least one cell type. There were more DEGs responsive to salt than to ABA in all species (Supplementary Fig. 23). Consequently, the responsive ortholog groups were more frequent for salt than for ABA (a total of 6,848 and 30,133 ortholog groups from all cell types for ABA and NaCl response, respectively) (Supplementary Fig. 29).

To determine whether the observed expression shifts were greater than expected by chance, we developed a bootstrapping approach that preserved the original data structure.This allowed us to generate an expected distribution and calculate the *p*-values for each pattern (see Methods). A gain of response was recorded when an ortholog switched its expression from a non-DEG to a DEG, whereas a loss of response was recorded when an ortholog switched from a DEG to a non-DEG within one lineage, relative to all other orthologs in the group. Lineage-specific gains in stress response were consistently observed less than expected and exhibited a wide distribution across cell types and species (Fig. 6a and Supplementary Fig. 29). Contrastingly, frequencies for lineage-specific loss of stress responses exceeded expected frequencies. These occurrences were confined to pericycle and procambium cells in response to ABA but were pervasive across most cell types under salt stress (Fig. 6a and Supplementary Fig. 29). This distinctive pattern suggests that transcriptional responses to stress in xylem and phloem cells are particularly stable across species. Synapomorphic (shared derived traits from a common ancestor) and convergent (independently evolved) responses were observed at frequencies greater than expected, especially in comparison to divergent responses (Fig. 6a and Supplementary Fig. 29). We additionally compared the response shifts among ortholog groups with gene duplication events and observed nearly identical trends as the 1-to-1 orthologs (Supplementary Fig. 30), indicating the transitions between DEGs and non-DEGs are likely to be insensitive to local duplications.

**Fig. 6.**
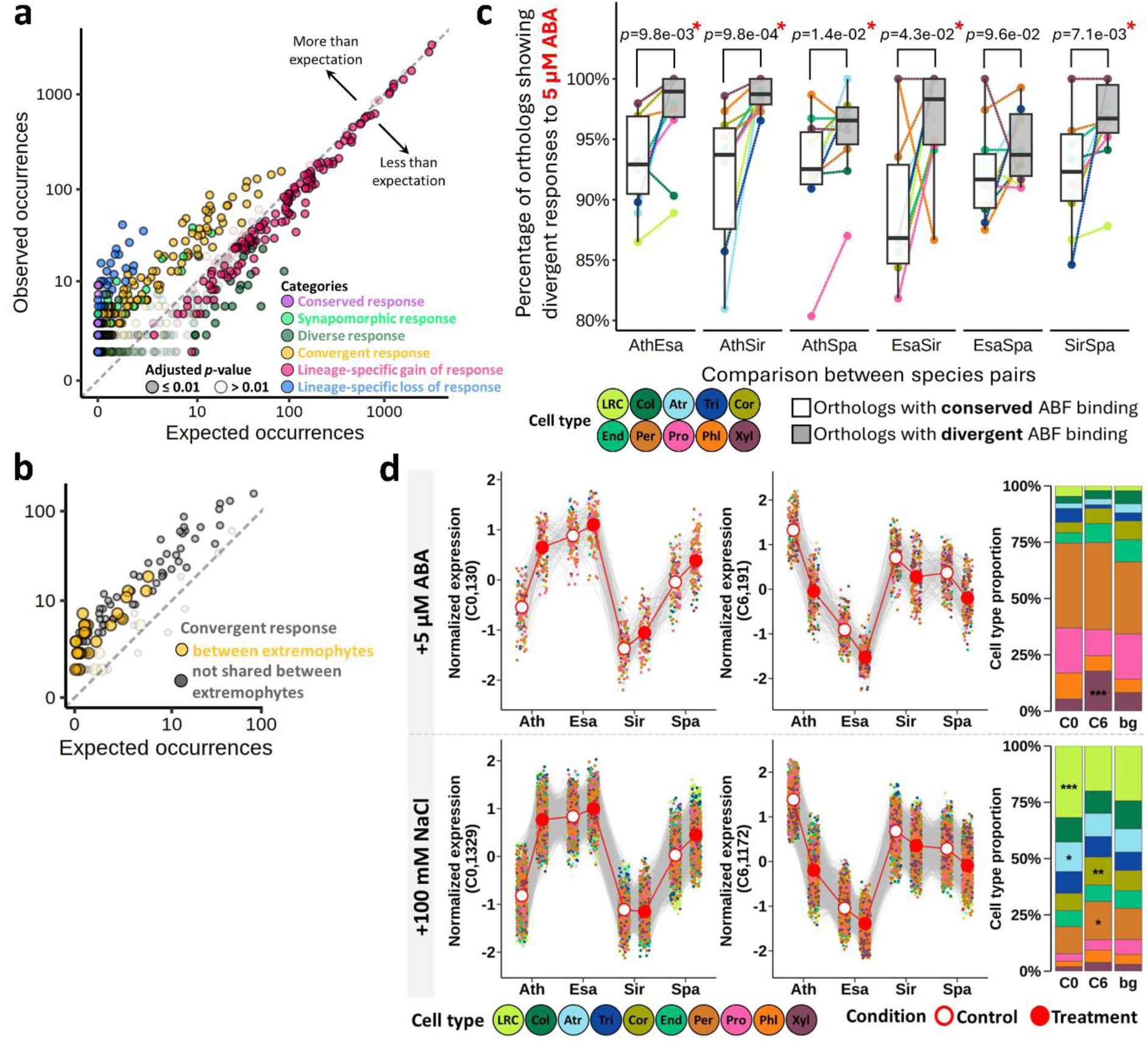
Stress responses between orthologs. a-b Differences between observed and bootstrapped (10,000 times) occurrences of response patterns among all 1-to-1 orthologs (a) and orthologs showing convergent response (b). The darker and lighter color intensities represent significant and insignificant deviation of the observed occurrences from the expected distribution, respectively, at Benjamini-Hochberg-adjusted *p*-value ≤ 0.01. c Percentage of divergently regulated orthologs (in response to 5 µM ABA) with conserved or divergent ABF binding. ABF binding was defined as the presence of ABF binding events in the 2-kilobase upstream of the transcription start site using the DAP-seq data from Sun *et al.*, 2022^9^. See Supplementary Fig. 33a for more explanations. Numbers on the top represent *p*-values determined by the Wilcoxon test. Asterisks represent significance at the *p*-value cutoff of 0.05. d Contributions of each cell type to co-expression networks. Cluster ID and the number of orthologs in the cluster are shown in parentheses in the Y-axis labels. Each gray line represents the expression of an ortholog across conditions and species. The red line indicates the median of all orthologs in the given cluster. Cell type proportions of the cluster were compared to background (bg) where orthologs from all clusters were averaged (see Methods for details). Asterisks represent significance determined by hypergeometric test, * *p*-value < 0.01, ** *p*-value < 0.001, *** *p*-value < 0.0001. LRC, Lateral Root Cap; Col, Columella; Atr, Atrichoblast; Tri, Trichoblast; Cor, Cortex; End, Endodermis; Per, Pericycle; Pro, Procambium; Phl, Phloem; Xyl, Xylem. Ath, Arabidopsis; Esa, *Eutrema salsugineum*; Sir, *Sisymbrium irio;* Spa, *Schrenkiella parvula*.

As shown in Fig. 6b and Supplementary Fig. 29, convergent responses were found between species from different lineages regardless of their phylogenetic relationship. We identified orthologs exhibiting convergent stress responses shared between the two extremophytes, and found 50 and 91 orthologs responding to ABA and NaCl treatments, respectively (Fig. 6b and Supplementary Fig. 29). We also detected 123 and 683 orthologs that were only responsive to ABA and NaCl in non-extremophytes (Fig. 6b and Supplementary Fig. 29). While the ABA-responsive orthologs shared between the two non-extremophytes were mostly enriched for nucleic acid metabolism, those shared by the extremophytes were mostly associated with response to abiotic stress (Supplementary Data 8). When exposed to NaCl, defense response pathways were strongly enriched among orthologs that were induced only in non-extremphytes or those orthologs repressed only in the extremophytes (Supplementary Data 8).

Given that ABA-responsive element binding factors (ABFs) are key regulators of ABA responsive gene expression, we next asked whether the divergent ABA and salt responses observed across species could, in part, result from differences in ABF binding. To answer this, we compared the differential regulation of orthologs under ABA treatment with ABF binding in their promoters, as previously identified using DAP-seq^9^. We observed consistently higher percentages of genes with ABF promoter binding among ABA induced genes compared to repressed genes or NaCl responsive genes (Supplementary Fig. 31). Additionally, regardless of species, genes with more than one ABF binding event were generally differentially regulated in more cell types than those with only one ABF binding event upon ABA treatment, and to a much lesser extent, NaCl treatment (Supplementary Fig. 32). Interestingly, we found that orthologs with divergent ABF binding were more likely to be divergently regulated by ABA, but not by NaCl (Fig. 6c and Supplementary Fig. 33). There was a substantial portion of orthologs (roughly 86% to 94%) showing different ABA responses even when all orthologs in the group had conserved ABF binding sites in their promoters (Fig. 6c). Together, these results indicate that divergence in ABF binding among orthologs can partially explain their divergent responses to ABA but that the divergent salt responses likely involve more complex ABA-independent regulation.

### Cell types contribute differentially to the stress-prepared transcriptomes in the extremophytes

Previous studies examining salt stress responses at the organ level in Arabidopsis and its extremophyte relatives have suggested that extremophytes exhibit stress-prepared transcriptomes, characterized by minimal overall changes to stress treatment, which may serve as a key determinant of their resilience to stress^10,40,41^. We aimed to investigate whether the stress-prepared state at the organ level in extremophytes resulted from a collective response of individual cells exhibiting a uniform low response or if contrasting responses in different cell types culminated in the absence of a net transcriptomic response captured at the organ level. We examined co-expression networks of all stress-responsive orthologs across species and conditions from all cell types. This resulted in 18 clusters with distinct co-expression patterns (Supplementary Fig. 34) with uneven representation of each cell type.

We calculated the contribution of each cell type to the overall co-expression pattern observed. We focused on the co-expression clusters, clusters C0 and C6, where the orthologs from the extremophytes showed a constitutively-prepared expression comparable to the stress-induced or - repressed expression of their counterparts in Arabidopsis (Fig. 6d). In ABA co-expression cluster 0 (4%-11% of the total clustered ABA-responsive orthologs in different cell types), where extremophyte orthologs exhibited consistently higher expression both before and after ABA exposure that those in the non-extremophytes, all cell types contributed evenly to this constitutive expression pattern (Fig. 6d, upper left). Conversely, in cluster 6 (4%-16% of the total clustered ABA-responsive orthologs in different cell types), characterized by the constitutively low expression in the extremophytes, the predominant contributors were xylem cells (Fig. 6d, upper right). In response to salt, the high constitutive expression of the extremophytes in cluster 0 (6%-14% of the total clustered NaCl-responsive orthologs in different cell types) was primarily attributed to lateral root cap and atrichoblast cells, whereas the consistently low expression of the extremophytes in cluster 6 (8%-16% of of the total clustered NaCl-responsive orthologs in different cell types) was predominantly contributed by cortex and pericycle cells (Fig. 6d, bottom panel). The co-expressed orthologs from lateral root cap and atrichoblast in cluster 0 under salt were both predominantly enriched for RNA metabolism- and ribosome biogenesis-related processes (Supplementary Fig. 35), which were also among the most responsive processes in Arabidopsis outer cell layers (Fig. 5d). This underscores the tendency for certain cell types to play a more substantial role in shaping the collective stress-prepared signal, as opposed to an equal contribution from all cell types.

Moreover, we noticed that only a subset of, but not all, surface cell types, including LRC and atrichoblasts, exhibited disproportionately higher contribution to the organ-level stress-prepared state upon salt treatment. However, in response to ABA, more internal cell types, including cortex and pericycle cells, showed stronger contributions to the stress-prepared signal than the surface cell types (Fig. 6d). These results suggest that the contribution of individual cell types to organ-level stress preparedness is likely shaped by a combination of their spatial position within the tissue, the nature of the stress, and the cell type-specific capacities for stress perception and signal transduction.

### Cell type assignments and sub-genome dominance at cell-type specificity in a polyploid

To assess the generality and transferability of our comparative single-cell framework developed for diploid species, we next applied our approach to the polyploid *Camelina sativa*. Unlike the four diploid species analyzed above, *C. sativa* contains three closely related sub-genomes, providing an opportunity to examine how cell-type identity and stress-responsive transcriptional programs are distributed among sub-genomes and homeologs in a polyploid genomic context. *C. sativa* is an allohexaploid in the Brassicaceae lineage I that has emerged as a promising oil crop for sustainable biofuel production^42^. *C. sativa* can grow in diverse environments, but its overall growth, seed oil yield, and oil quality are severely impacted by salt and other abiotic stressors^43–45^. We generated a single cell atlas of 27,441 root cells from *C. sativa* under control, 5 µM ABA, and 100 mM NaCl, as described for the other four species (Fig. 7). Using the cross-species cell-type annotation strategy developed for this study, we were able to identify all ten major root cell types in *C. sativa* (Fig. 7a), which exhibited a clear separation between inner and outer cell layers as observed for other Brassicaceae species (Fig. 7b). The individual sub-genomes from *C. sativa* did not significantly affect the integration between *C. sativa* and Arabidopsis or the cell-type assignments (Supplementary Fig. 36), suggesting a relatively even representation of the three sub-genomes. Our result is consistent with the undifferentiated sub-genome structure previously reported for this species^46,47^. Moreover, we found that sub-genome G3 consistently exhibited a higher global expression compared to the other sub-genomes across cell types and conditions, except for columella cells where sub-genome G1 showed the highest mean expression under control and ABA-treated conditions (Supplementary Fig. 37). This agrees with prior work that has established that the sub-genome G3 in *C. sativa* exhibited partial expression dominance over the other two sub-genomes^48^.

**Fig. 7.**
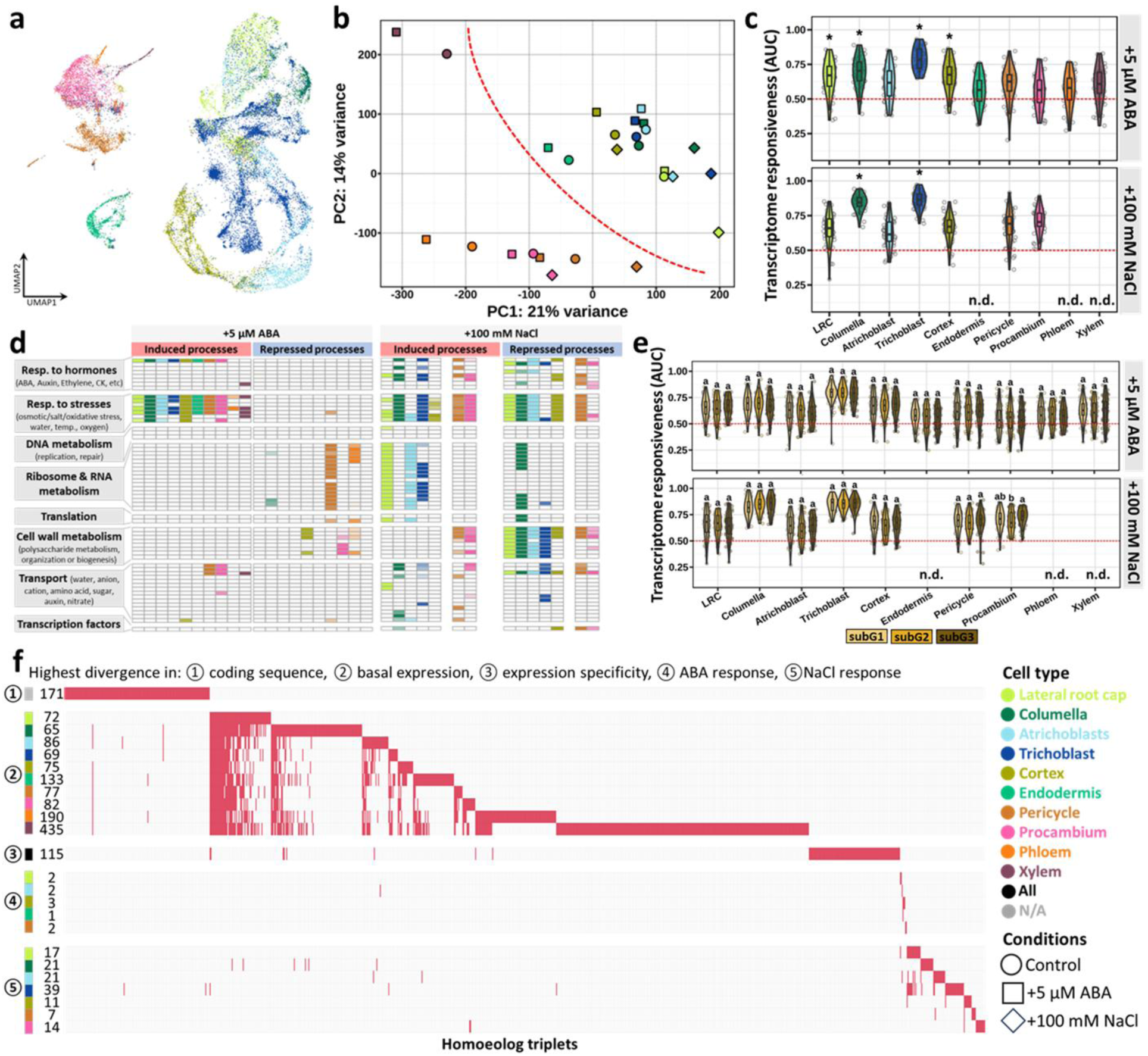
Cell type stress responses in *C. sativa*. a Root single cell atlas (27,442 cells) for *C. sativa*. The atlas included control, 5 μM ABA- and 100 mM NaCl-treated samples. b Principal component analysis of pseudo-bulk cell-type transcriptomes. Each symbol represents a pseudo-bulk cell-type transcriptome from control, 5 μM ABA- and 100 mM NaCl-treated samples. The color and the shape of each symbol correspond to the cell type and the condition, respectively. c Comparison of transcriptome responsiveness as averaged Augur area under the curve (AUC) scores across cell types in response to ABA and NaCl. AUC here quantifies the separability between control and treatment samples for each cell type. AUC = 0.5 indicates random separability; AUC = 1.0 indicates perfect separability. Asterisks indicate significance after Wilcoxon testing (n = 50) at adjusted *p*-value cutoff of 0.05. d Summary of GO functional enrichment among DEGs combined from all sub-genomes in response to ABA (top) and NaCl (bottom) for each cell type in *C. sativa*. The detailed list of GO terms is present in Supplementary Data 7. The shade represents the *p*-value assigned by the enrichment test at adjusted *p*-value < 0.05 with darker shades indicating smaller *p*-values. e Comparison of transcriptome responsiveness (as AUC scores) for each sub-genome in *C. sativa*. AUC here quantifies the separability between control and treatment samples using each sub-genome for each cell type. AUC = 0.5 indicates random separability; AUC = 1.0 indicates perfect separability. Lowercase letters on top of violin plots indicate significantly different groups (n = 50) as determined by one-way analysis of variance (ANOVA) with post-hoc Tukey honest significant difference (HSD) testing at *p*-value < 0.01. f Ranked homoeologous triplets based on their divergence in coding sequence (①), expression (②), cell type specificity (③) as well as expression changes in response to stresses (④ and ⑤). Numbers on the left indicate the number of candidates considered for each category. Endodermis, phloem and xylem cells were removed from the comparison for NaCl treatment (n.d.) due to insufficient number of cells (< 50 cells) in c-f. N/A, not applicable.

Although sub-genome expression dominance has been extensively examined in many polyploid species, it still remains unknown whether sub-genomes differentially contribute to stress response. To determine this, we identified DEGs from each sub-genome in each cell type (Supplementary Data 6). Despite the presence of a species-specific response, the impact of stress on cell type transcriptomes and the functional processes represented by the DEGs in *C. sativa* largely resembled the other four species (Figs. 7c-7d and Supplementary Fig. 38). The proportion of DEGs did not significantly differ between sub-genomes (0.2%-2.4% and 1%-25% of expressed genes across cell types for ABA and NaCl treatments, respectively), irrespective of whether we considered all detected genes within individual sub-genomes or only complete homeologs (defined as triplets)^48^ (Supplementary Fig. 39). Consistently, transcriptomic responsiveness measured as AUC scores did not differ significantly among the sub-genomes for most cell types under ABA or NaCl treatments (Fig. 7e). These findings show that, in contrast to the expression bias observed among sub-genomes across cell types under stress-neutral conditions, stress responses did not further differentiate the sub-genomes in *C. sativa*, suggesting a lack of large-scale sub-genome dominance in stress responses.

The recent polyploidization of the *C. sativa* genome^46^ allowed us to ask what genomic changes corresponded to divergence in gene expression across the three copies of each homeolog (18,565 complete triplets). First, to quantify coding-sequence divergence, we calculated the nonsynonymous/synonymous rate ratio (ω) for each complete triplet. Second, to assess the regulatory sequence divergences among homeologs, we considered divergence of expression at three different parameters: basal level expression, cell-type preferential expression, and expression changes in response to treatment. This analysis identified triplets exhibiting high divergence (top 1% for each metric) among homeologs in 1) coding sequences (171 triplets), 2) basal expression level (1,384 triplets), 3) expression specificity (115 triplets), and 4) stress responses (140 triplets) (Fig. 7f; Supplementary Data 9). We found that these highly divergent triplets were often shared across different cell types within each defined category, particularly for these showing divergent basal expressions. Few triplets were divergent across multiple categories with only 6 out of the 171 fast evolving triplets also showing high expression divergence (overlap between group ① and groups ② and ⑤ in Fig. 7f). These results indicate that coding-sequence and expression level divergences among triplets are largely uncoupled from one another in *C. sativa*.

## DISCUSSION

Comparing stress responses at the cellular level between extremophytes and non-extremophytes holds the potential to identify innovations in stress resilience that may expedite the development of climate-change ready crops. Here, we selected five closely related Brassicaceae species with distinct responses to ABA and NaCl and generated their root transcriptomes at single cell resolution. We developed a two-step approach for cross-species and cross-condition cell-type annotations and confirmed the resulting annotation by *in situ* hybridization of selected markers preferentially expressed in specific cell types, conditions, or species. In addition to serving as a cell atlas resource for Brassicaceae roots under control and stress conditions, our method of cell-type identity and response assessments under induced stressed conditions serves as a model for other multi-species and treatment comparisons. Our approach was further applied on an allohexaploid oilseed crop, *C. sativa* to demonstrate the versatile detection and transfer of cell-type response features between diploid and polyploid genomes. We anticipate these data and the methods presented in our study will provide new opportunities and perspectives for comparative single cell transcriptomics to enrich our understanding of cell-type-specific evolution across species. The searchable database we established to disseminate these data will facilitate their broader use by the community. We envision that this resource will serve as a foundation for downstream functional and experimental studies, especially in a phylogenetic context. It not only facilitates targeted visualization and manipulation of cell types of interest, especially in species where such information is currently limited, but also guides the selection of candidate genes for testing cell type-specific stress responses across species.

To date, experimentally validated root cell-type markers in plants have been largely derived from model species like Arabidopsis^49,50^. However, the transferability of these markers to other species where root cell-type markers are unavailable has not been determined. We evaluated the conservation of previously established Arabidopsis cell-type markers in the selected Brassicaceae species and found that only 75 of the 241 Arabidopsis markers could serve as cell-type markers across all species tested. The main basis for this lack of conservation was the low expression levels among orthologs in related species. This suggested that moderate to high expression of orthologs, in addition to preferential expression in the target cell type, should be an important practical feature of widely used cell-type markers. We have newly identified over 300 cell-type preferentially expressed genes that are conserved across these Brassicaceae species (Supplementary Data 3), similar to the resource available for grass species^39^. A previous study comparing rice and Arabidopsis demonstrated that marker genes for the majority of root cell types were poorly conserved^20^. This raises interesting questions about the relationship between conservation of cell-type markers and phylogenetic distances among species. For example, at which evolutionary distances do cell-type marker conservation erode and how do genes lose or gain cell-type specificity? Advances in our understanding of cell type evolution will inform innovations in synthetic biology, particularly cell-type specific engineering of traits or metabolite production across species.

Identifying new cell populations across species enabled by scRNA-seq provides an unprecedented opportunity to trace distinct cell states or origins of cell types and infer cell-type evolution^51^. The presence of an extra cortex cell layer specific to *S. irio* and *S. parvula* root anatomy was recapitulated with corresponding scRNA-seq data, which suggested the additional cortex layer is transcriptionally specialized for flavonoid biosynthesis. Such evolutionary innovations in cell layer compartmentalization may enable the spatial partitioning of growth and stress-relevant metabolic functions. However, the exact link between this biased metabolic partitioning and plant growth and stress responses remains unresolved. A second layer of cortex, called middle cortex, can be also detected in Arabidopsis^52,53^, but at a later developmental stage of roots. Whether the inner cortex layer in *S. irio* and *S. parvula* is developmentally and functionally similar to the Arabidopsis middle cortex, whether the flavonoid pathway is also preferentially partitioned to middle cortex in Arabidopsis, or whether the early development of a second cortex layer presents an evolutionary derived trait remains to be elucidated.

Beyond the comparisons under stress-neutral conditions, our dataset enables interrogation of stress responses in each cell type across species. Our results demonstrate that environmental stresses trigger distinct, cell-type- and stress-specific changes in transcriptome divergence across species, with cell types exhibiting differential contributions to stress-responsive co-expression networks. These findings highlight the distinct roles of cell types in driving stress response divergence among species and the approaches developed here can therefore facilitate the identification of cell types that play more significant roles in determining the divergence.

The cell-type responses of *C. sativa* highlighted transcriptional processes that are shared as well as unique to root cell types in the polyploid oil crop. The emerging diversity of cell-type responses shows the necessity of examining stress responses of crop species to assess the transferability of response mechanisms from model plants. Although the sub-genome G3 demonstrated a greater contribution of highly expressed genes across the majority of cell types, the three sub-genomes within this allohexaploid have not yet differentiated to the extent of contributing to distinct stress responses in any single cell type. This lack of sub-genome stress response dominance suggested that all three sub-genomes in *C. sativa* contribute comparably to stress responses. Such distributed regulatory contributions may increase robustness by providing functional redundancy among homeologs and buffering against divergence or loss of individual gene copies. However, we do not infer that balanced sub-genome contributions directly enhance stress tolerance, rather, these results indicate that *C. sativa* does not restrict stress-responsive regulation to a single genomic origin. Our findings on response divergence rarely co-occurring with sequence and expression divergence within individual homeolog groups suggest the evolution of these divergences occurs largely independently. Together, these findings add insights into the coordination among sub-genomes in polyploid species and the fate of homeologs, and how we may generate agriculturally preferable neopolyploid crops^54^.

## METHODS

### Plant materials and growth conditions

*Arabidopsis thaliana* (Arabidopsis, ecotype Col-0), *Eutrema salsugenium* (ecotype Shandong) and *Schrenkiella parvula* (ecotype Lake Tuz) seeds were surface sterilized using a 30% (v/v) bleach, 0.1% (v/v) Triton X-100 solution for 10 min and rinsed with sterile water 5 times.

*Sisymbrium irio* (original accession from Chris Pires, University of Missouri) seeds were sterilized using 95% EtOH for 5 min, then 20% (v/v) bleach, 0.1% (v/v) Tween 20 solution for 5 min, and rinsed with sterilize water 4 times. *Camelina sativa* (ecotype DH55) seeds were surface sterilized using 20% (v/v) bleach, 0.1% (v/v) Triton X-100 solution for 15 min and rinsed with sterile water 4 times. After seed sterilization, seeds were incubated at 4°C for 2∼3 days prior to plating. Seeds were placed densely in 4∼5 rows and incubated vertically on standard media (¼× Murashige and Skoog (MS) nutrients, 1% sucrose, 0.05% MES, 1% agar, adjusted to pH 5.7 with 1 M KOH) covered with 100/47 μm sterilized mesh and incubated under continuous light at 22°C for 3 days (*Camelina sativa*) or 4∼6 days (Arabidopsis, *Eutrema salsugenium*, *Schrenkiella parvula* and *Sisymbrium irio*) until roots were ∼2-3 cm in length. For ABA treatment, the mesh with the seedlings was transferred to plates containing either 5 μM ABA or standard media (as control) and incubated for 3 hrs. For NaCl treatment, the mesh with the seedlings was transferred to plates containing either 100 mM NaCl or standard media (as control) and incubated for 24 hrs.

### Preparation of root samples for scRNA-seq

After seedling growth on plates containing ABA, NaCl, or standard media, primary root tips, which approximately corresponded to the top one third (0.7-1 cm) from the root tips, were harvested from each species. The harvested root tips included root cap, meristematic, elongation, and differentiation zones. Harvested root tips were placed into a 35 mm petri dish containing a 70 um strainer and 4 mL enzyme solution (1.25% [w/v] Cellulase [“ONOZUKA” R-10, Yakult], 0.1%[w/v] Pectolyase [P-3026, Sigma-Aldrich], 0.4 M Mannitol, 20 mM MES [pH 5.7], 20 mM KCl, 10 mM CaCl_2_, 0.1% [w/v] bovine serum albumin). The sample dish was rotated at 85 rpm for 50 min at 25°C. The root tips were then transferred to a glass slide and gently pressed with another glass cover using a pencil rubber to release protoplasts. The tissues were transferred back to the enzyme solution and incubated for another 10 minutes. The protoplast solution was filtered through a 70 μm filter, and then through a 40 μm filter twice using 50 mL conical tubes. The filtered solution was centrifuged at 500g for 10 min at 22°C using a swinging bucket rotor and the supernatant was discarded. Protoplasts were resuspended using 500 μL Solution A (0.4 M Mannitol, 20 mM MES [pH 5.7], 20 mM KCl, 10 mM CaCl_2_, 0.1% [w/v] bovine serum albumin) and transferred to a 2 mL round bottom tube. After centrifugation at 200g for 6 min, the pelleted protoplasts were resuspended with 30–50 μL Solution A to achieve the desired cell concentration (∼1,000 protoplasts/μL).

### scRNA-seq library construction and sequencing

The protoplast suspension was loaded into Chromium microfluidic chips (Next GEM Single Cell 3’ kit) and barcoded with a 10X Chromium Controller (10X Genomics). mRNA from the barcoded cells were subsequently reverse-transcribed and sequencing libraries constructed with reagents from a Chromium Single Cell 3’ reagent kit v3.1 chemistry dual index (10X Genomics) according to the manufacturer’s instructions. The quality of cDNA and final library was assessed with a Bioanalyzer (Agilent). Paired-end sequencing was performed with Illumina NovaSeq 6000 according to the manufacturer’s instructions (Illumina).

### Bulk RNA-seq library preparation and sequencing

Seedlings were grown under the same stress-neutral conditions as the scRNA-seq samples. RNA was collected from the lower 1/3 of primary roots (which includes the entire developing root tip) from each plant species using the RNeasy Plant Mini Kit (Qiagen). Library construction and sequencing were performed by the University of Michigan Sequencing Core using the Illumina TruSeq Kit followed by paired-end sequencing on Illumina NovaSeq 6000. Three replicates were obtained per species.

### Orthologous gene sets

We used two different sets of orthologous genes in our study: (1) comparative analyses among Arabidopsis, *E. salsugenium*, *S. irio*, and *S. parvula* were performed using 15,198 unambiguous 1-to-1 ortholog groups between the four species. These 1-to-1 orthologues were identified by comparing the primary protein-coding sequences and their syntenic positions using CLfinder^55^ with MMSeq2^56^ as the aligner with default settings. (2) Species-specific analyses were performed using all orthologs expressed in the given species. The orthologs between each non-Arabidopsis species (including *E. salsugenium*, *S. irio*, *S. parvula*, and *C. sativa*) and Arabidopsis were identified using the following steps: orthologous gene pairs of the protein-coding genes between each non-Arabidopsis species and Arabidopsis were determined reciprocally using MMseqs2^56^ and were subsequently merged to generate a non-redundant list. Only orthologous gene pairs that were considered as orthologs by both OrthoFinder (v2.5.4)^57^ with granularity -I set to 1.6 and CLfinder were kept. This filtered list was supplemented with reciprocal best hits identified by blastp^58^ to generate the final non-redundant orthologues between each non-Arabidopsis species and Arabidopsis.

### Raw reads processing

Raw single-cell RNA-seq FASTQ files were analyzed by Cell Ranger (v5.0.1, 10X Genomics) to generate gene-by-cell count matrices. For each species, we retrieved the reference genome and corresponding annotation as follows: Arabidopsis (TAIR10) from The Arabidopsis Information Resource (TAIR; https://www.arabidopsis.org/), *C. sativa* (v2) from Brassicaceae Database (BRAD; http://brassicadb.cn), *E. salsugineum* (v1.0) and *S. parvula* (v2.2) from Phytozome 12 (https://phytozome-next.jgi.doe.gov/) (Phytozome ID 574 and 173, respectively), reference genome for *S. irio* (v1.0) from Brassicaceae Database (BRAD; http://brassicadb.cn) with the updated gene model annotation^9^ from the CoGe database (https://genomevolution.org/) with the Genome ID 57216. To account for the low accuracy of 3’ UTR annotation in non-Arabidopsis species, we extended the 3’ UTR of all genes in these species by 500bp, as reported before^59^. Reads from each species were aligned to the corresponding reference genome and counts were quantified with the Cell Ranger ‘count’ command with default parameters.

### Comparison of gene expression from bulk and pseudo-bulk samples

Raw 150-bp paired-end reads were quality checked with FastQC (v0.11.5; https://www.bioinformatics.babraham.ac.uk/projects/fastqc) and aligned to the respective genomes with HISAT2 (v2.2.1)^60^. Gene counts were calculated using featureCounts (v2.0.3)^61^ with the modified gene annotations. Pseudo-bulk gene counts from each scRNA-seq sample were generated by aggregating Cell Ranger-derived counts for all filtered cells. We next implemented edgeR (v3.36.0)^62^ to identify genes that exhibited differential expression (FDR ≤ 0.001, absolute log_2_FC ≥ 2) between the bulk and pseudo-bulk samples to be aware of any genes that may deviate between how the data was generated. The list is provided in Supplementary Data 10.

### Identification of doublets

Doublets in each scRNA-seq dataset were detected with DoubletFinder (v3)^63^ and subsequently removed prior to any downstream analyses. Three input parameters are required for doublets prediction: the number of expected real doublets (nExp), the number of artificial doublets (pN) and the neighborhood size (pK). We selected a stringent nExp of 7.5% based on the estimated doublet rate in 10X Genomics Chromium Single Cell 3’ Reagent Kit User Guide, which was subsequently adjusted according to the proportion of homotypic doublets and doublet ratio. For pN, the default value 25% was used as DoubletFinder performance is largely invariant of pN^63^. The optimal pK value was determined by sequentially loading pre-processed Seurat ^22^ data into the “paramSweep_v3 (PCs = 1:15)”, “summarizeSweep”, and “find.pK” functions. For each sample from each species, the single easily discernible maximum pK value was selected as the optimal pK parameter. Only cells which were flagged as singlets were used for downstream analyses.

### Data integration, clustering

Data integration and clustering were performed using R package Seurat (v4.0.5)^22^. Prior to data integration, we further removed cells in which the numbers of expressed genes were less than 400 and more than 10,000 or < 5% mitochondrial reads, and genes expressed in < 3 cells.

For integration of datasets from each species, we first log-normalized each dataset using the *NormalizeData* function and identified the top 2,000 highly variable genes (HVGs) using a variance-stabilizing transformation in *FindVariableFeatures*. We then used *SelectIntegrationFeatures* to determine 2,000 integration features, which were subsequently used in *FindIntegrationAnchors* to decide integration anchors between datasets. Datasets from each species were eventually integrated using *IntegrateData* with 30 principal components (PCs). The integrated dataset for each species was scaled and reduced dimensionally with Principal Component Analysis (PCA) followed by Uniform Manifold Approximation and Projection (UMAP) using 50 PCs. *FindNeighbors* and *FindClusters* were used to determine the nearest neighbors and clusters, respectively.

For global integration of datasets from all species, feature names in each of the individual stringently filtered datasets were replaced by their corresponding ortholog group ID. Each dataset was normalized using the *SCTransform* function, and then *SelectIntegrationFeatures* was used to identify 3,000 integration features. Integration features were combined using *PrepSCTIntegration* and integration anchors were selected using *FindIntegrationAnchors*.

Datasets from all species were then integrated at once using the Seurat “SCT” approach in *IntegrateData* function with setting Arabidopsis control datasets as the reference. Finally, dimensional reduction of the integrated dataset was performed using the UMAP algorithm with the top 100 PCs identified by PCA.

Dimensionality-reduction methods were used for distinct analytical purposes in our study. UMAP was employed for visualization, integration, and cell-type annotation of single-cell data, where preservation of local neighborhood structure is advantageous. In contrast, PCA was applied to pseudo-bulk cell-type transcriptomes for cross-species and cross-condition comparisons, as PCA preserves global variance structure and enables direct interpretation of transcriptomic similarity and divergence. Because UMAP embeddings are nonlinear, stochastic, and sensitive to parameter choices, and they are not used for quantitative distance- or variance-based comparisons.

### Cell type assignment

#### Step 1. Establishment of a reference atlas using Arabidopsis control samples

*Correlation-based annotation* (Step1 - M1.1 in Supplementary Fig. 4): We obtained bulk RNA-seq data^64^ and ATH1 microarray data^65^ for 14 Arabidopsis root cell types isolated with fluorescence activated cell sorting (FACS) as fragments per kilobase of transcript per million fragments mapped (FPKM) and normalized probe values, respectively. We then computed the expression variance for all genes within each dataset to identify highly variable genes (HVGs) and categorize them into percentile bins. We hypothesized that the transcriptome signature of an individual cell should remain most similar to that of the cell type it originates from regardless of the HVGs used. Therefore, we calculated the Pearson correlation coefficient between each cell from Arabidopsis control samples and the whole-transcriptome reference expression profiles for cell types using HVGs from top 10% to 100% percentile bins. At each percentile bin, each cell was annotated as the cell type with the highest correlation value.

*Marker-based annotation* (Step1 - M1.2 in Supplementary Fig. 4): We retrieved the normalized counts and uniform manifold approximation and projection (UMAP) embeddings in Seurat for all cells from Arabidopsis control samples, which were used to calculate the enrichment scores of all genes for each cell using a cluster-independent approach in SEMITONES^66^. Similarity between cells was estimated by a radial basis function kernel over the multidimensional UMAP space, and the enrichment scores for all genes were computed using a simple linear regression framework. We subsequently extracted the enrichment scores for 454 (after removing stele markers) known Arabidopsis root cell type marker genes (Supplementary Data 2). This set was compiled through an extensive literature review^15–21^. To identify statistically significantly enriched markers for individual cells, an empirical distribution of enrichment scores for each cell was obtained by permutating expression data 100 times. A gene is considered significantly enriched in the given cell if its enrichment score is more than n standard deviations (nsds) away from the mean of the empirical distribution. We determined the significantly enriched markers at nsds of 5, 8, 10, 15, 20, 25, 30, 35, 40, 45, 50 for each cell, and annotated the cell according to the marker with the highest enrichment score at each nsds.

*Integration-based annotation* (Step1 - M1.3 in Supplementary Fig. 4): We used a previous Arabidopsis root single-cell study^21^ with cell type annotations for individual cells.

*FindTransferAnchors* from Seurat was used to find transfer anchor cells using the published data as the reference and cells from Arabidopsis control samples in our dataset as the query.

*TransferData* was subsequently employed to predict the identity of each cell in the query dataset. These steps were run with different thresholds for the key parameters: ndim at 30, 50 to 500 with an increment interval of 50; kscore at 10, 30, 50; kweight at 10, 20, 30, 50, 80, 100. Cell type prediction for each cell was extracted at every parameter combination.

*Combination of annotation methods* (Step1 in Supplementary Fig. 4): We first determined the consensus annotation in each of the three annotation approaches for each cell by identifying the most recurring (typically over 50%) annotation from all the runs. The final annotation for each cell in Arabidopsis control samples was assigned by combining the information from the three annotation approaches. If a cell was annotated to the same cell type from at least two of the three approaches, then it was assigned that cell type. In rare cases where a cell cannot be annotated using a consensus, the cell type was assigned according to the marker-based annotation.

#### Step 2. Annotation transfer from Arabidopsis to other species

The integration of cells from different treatments and different species in our dataset suggested there were no treatment- or species-specific cell populations. We next employed three different reference-based cell type annotation algorithms, including integration-based prediction (Step2 - M2.1 in Supplementary Fig. 4) in Seurat^22^, clustering-based prediction (Step2 - M2.2 in Supplementary Fig. 4) in Symphony^23^, and the machine learning-based prediction (Step2 - M2.3 in Supplementary Fig. 4) in scArches^24^, to transfer cell type annotation from Arabidopsis control samples to treatment samples and samples from other four species. To avoid probabilistic annotations in these methods, we varied the key parameters in each method: Seurat was run with ndim at 30, 50 to 500 with an increment interval of 50, kscore at 10, 30, 50, kweight at 10, 20, 30, 50, 80, 100; Symphony was run with hvg_num from 2000 to 14000 with an interval of 2000, pc_num from 20 to 100 with an interval of 20, and k_num at 5, 10, 20, 30, 50, 80, 100; and scArches was run with hvg_num from 2000 to 14000 with an interval of 2000. Similar to the approach we used to annotate cells in the Arabidopsis reference, we next identified the most recurring cell type annotation for a cell as the consensus annotation for that cell in each of the three methods. The annotation for a cell with the highest votes from all three methods was assigned as the final annotation for that cell. In rare cases where a cell was annotated differently from the three methods, we compared the average probability given by each method and assigned the cell to the prediction with the highest probability.

### Cell-type marker gene identification

To identify cell type marker genes for each species, we used the Seurat *FindMarkers* function on scaled expression using the default Wilcoxon test. Significant markers were selected based on the following criteria: adjust *p*-value ≤ 0.01, average log fold change ≥ 1, cell type specificity ≥ 0.9 and pct.diff > 0.75, where pct.dff is the ratio of the difference in the percentage of cells expressing the gene between the target cell type and the rest of the cells to the percentage of cells expressing the gene in the target cell type. To examine the expression of the cell-type markers, the cell-type pseudo-bulk expression was computed by normalizing the raw gene counts in each cell by the total counts of that cell, and these normalized counts were then averaged across all cells belonging to the same cell type. This cell-level normalization ensures that pseudo-bulk profiles reflect the average transcriptional state of individual cells rather than being dominated by differences in cell numbers or library size.

### Cell type specificity indexes

Cell type specificity index (τ) is based on the Tau metric of tissue specificity^67^ as following: 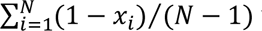 where N is the number of cell types and x_i_ is the expression level in cell type normalized by the maximal expression of the given gene across all cell types. For this analysis, the normalized expression as described for generating cell-type marker expression was used.

### Identification of orthologues with divergent expression

To identify divergently expressed orthologues (Fig. 3b), we compared gene expression of the 15,198 1-to-1 orthologues along the defined cell types across Arabidopsis, *E. salsugenium*, *S. irio*, and *S. parvula*. Within each species, we determined the genes that were differentially expressed between any pairs of cell types with Seurat *FindMarkers* function. We then compiled these genes from all four species and used the normalized cell-type pseudo-bulk expression (the cell-type pseudo-bulk expression was computed using the same approach as that used for cell type specificity and then z-score normalized across cell types) of these genes as the input to cluster gene expression along the ten cell types using an unsupervised soft clustering method implemented in the R package Mfuzz (v.2.54.0)^68^, which assigns probabilistic membership of each ortholog to expression-pattern clusters rather than enforcing hard classifications. The number of clusters and the fuzzification parameter required by Mfuzz were estimated by the Dmin and mestimate functions. Subsequently, we inferred within a phylogenetic framework, as described by Cardoso-Moreira *et al*.^69^, the probability that there were changes in the expressions of the 1-to-1 orthologues along the defined cell types, that is, that genes changed their cluster assignment in specific branches, using 0.001 as the probability cut-off. This probability-based threshold followed the original framework^69^ and was selected to prioritize robust expression shifts over marginal or noisy changes. In this framework, the ancestral cluster probabilities along the phylogenetic tree were calculated as the weighted averages from the child-nodes. The weights are given by the inverse branch lengths, which were retrieved from a previous study ^13^ using WebPlotDigitizer (https://automeris.io/WebPlotDigitizer), so that closer child-nodes have more weight. Since this framework relies only on quantitative expression measurements across homologous biological contexts and a resolved species phylogeny, it is agnostic to organismal lineage and can be applied to any systems that meet these criteria.

To construct the phylogeny for aquaporin subfamily NIPs and Tau class glutathione S-transferase GSTUs across species, we first identified members from these families in Arabidopsis from previous studies^70,71^, and extracted their orthologs from *E. salsugenium*, *S. irio*, and *S. parvula* using the ortholog network generated from CLfinder^55^. The protein sequences of genes in each subfamily were aligned using MAFFT^72^, the results of which were subsequently used for phylogenetic tree construction with FastTree^73^ with default settings. The inferred phylogenies were annotated with ggtree^74^ in R.

### Prioritization of cell types responsive to ABA and NaCl

To estimate the degree of transcriptomic change for each cell type under treatments (Fig. 5b), we used Augur^38^ to rank cell types based on their degree of response to biological perturbations (i.e. ABA or NaCl treatments in our current study). Augur uses a machine-learning frame to quantify the separability of perturbed and unperturbed cells as AUC scores, based on the assumption that cell types more responsive to a perturbation should be more separable than less affected cell types within high-dimensional single-cell measurements. Augur analysis was performed for ABA- and NaCl-treated samples separately compared to the control within each of the five species. The significance of cell type prioritization for each cell type was determined by comparing the AUC scores from all subsamples of the specific cell type of interest to all rest of cell types for each treatment using one-tailed Wilcoxon rank-sum tests^75^.

To compare cell-type transcriptomic responses across Arabidopsis, *E. salsugenium*, *S. irio*, and *S. parvula*, we modified the Augur method to estimate transcriptome divergence of each cell type between different species under each condition. Specifically, instead of using cells from different conditions, we fed cells from different species under the same condition into Augur. This process was repeated for all six pairwise combinations from the four species under each condition. For each pair of species, the AUC scores of each cell type from ABA- and NaCl-treated samples were compared to these from the control using Student’s t-Test.

### Pseudo-bulk differential expression analysis

Recent benchmark studies to assess methods to determine differentially expressed genes (DEGs) for scRNA-seq data have shown that pseudo-bulk methods, which aggregate cell-level counts into sample-level “pseudo-bulk” counts, outperform single-cell DE methods and achieve highest fidelity to the experimental ground truth^76,77^. Therefore, we employed a pseudo-bulk approach to detect genes that responded to the treatments for each cell type in each species. Pseudo-bulk transcriptomes were aggregated for each of the cell types by taking the sum of raw cell-level counts. Generalized linear model likelihood ratio test from edgeR (v3.36.0)^78^ was performed between the treatment and the control. For each comparison, cell types with less than 30 annotated cells under either treatment or control conditions were excluded from the differential expression analysis. A gene was considered differentially expressed in a given cell population if the false discovery-rate adjusted *p*-value was <= 0.01. Additionally, to calculate the transcriptomic distance between samples, the raw counts in each cell type for the 1-to-1 orthologs were extracted from each species and each condition, and then were normalized in DESeq2^79^ as variance stabilized expression. The variance stabilized expression was then used for PCA-based transcriptomic distance measurements.

### Identification of differential regulation of orthologs between species

We extracted DEGs from each species under each treatment that were included in the 1-to-1 orthologs and compared their response to the treatment across species to determine differentially regulated orthologs. We considered the stress response of the 1-to-1 orthologs as being upregulated and downregulated if log_2_FC > 0 and log_2_FC < 0, respectively, regardless of their response magnitude. To further distinguish observed differential regulation patterns between the orthologs which occurred above chance due to random sampling (Fig. 6a), we started with a fixed order of all 1-to-1 orthologs and then permuted the order of DEGs for 10,000 times independently in each species for each cell type and generated an expected occurrence distribution for each pattern when assuming independence between the species. The deviation of the observed occurrence for each pattern was computed as the number of standard deviations away from the mean of the corresponding expected distribution. The *p*-value of the observed occurrence for each pattern was calculated by fitting the observed occurrences into the corresponding bootstrapped distributions using R function *pnorm* and subsequently corrected for multiple testing using the Benjamini-Hochberg procedure in R function *p.adjust*. To additionally test the impact of gene duplication on the differential regulation of orthologs, we identified ortholog groups with more than one copy in only one species where identification of correct corresponding orthologs across species is feasible. The same approach was applied to these duplicated ortholog groups to determine the deviation between the occurrences and the expectations.

To compare the differential regulation among orthologs to their ABF binding events, we retrieved DAP-seq data of four ABFs in Arabidopsis, *E. salsugenium*, *S. irio*, and *S. parvula* from a previous study^9^. For each of the six possible species pairs, we extracted ortholog pairs where at least one gene responded to the treatments and at least one gene in the pair contained a minimum of one ABF binding event in its promoter region. Two kilobases upstream of the transcription start sites were used as the searched promoter sequence space for ABF binding. We then categorized these ortholog pairs into the following two groups: 1) orthologs with conserved ABF binding where both genes contained at least one ABF binding event in their promoters; and 2) orthologs with divergent ABF binding where only one gene contained at least one ABF binding event in its promoter. From these groups, we identified orthologs with only one gene responding to the treatments in each cell type, which we categorized as divergently regulated orthologs. The significance of differences in the percentage of divergently regulated orthologs between the two defined groups were determined with *t.test* function in R.

### Determination of cell type contribution to co-expression patterns

For each cell type under each treatment, orthologs that were considered differentially expressed in at least one of the four species were extracted, and subsequently combined for aggregated clustering. The normalized pseudo-bulk cell type expression levels of these orthologs were subjected to fuzzy k-means clustering implemented in aerie^80^. Ortholog pairs in each of the resulting clusters were further filtered based on: (1) the membership of a given ortholog pair was no less than 0.5; (2) the Pearson correlation between the expression of a given ortholog pair and the mean expression of the cluster it belongs to was no less than 0.8 with a *p*-value smaller than 0.01; and (3) the cluster needed to have a minimum of 50 members. To test the representation of each co-expression pattern among cell types under each treatment, we compared the proportion of orthologous DEGs derived from different cell types in each co-expression cluster to that combined from all clusters using R function *phyper* with lower.tail = FALSE. The resulting *p*-values were corrected for multiple testing using the Benjamini-Hochberg procedure in R function *p.adjust*.

### Identification of extra cortex population

To determine potential topological differences in cortex between species, we performed non-linear dimensional reduction for cortex cells from each species under control condition using Seurat^22^ to generate species-specific UMAP representations. The UMAP representation of cortex cells in *S. irio* was selected as the reference because of the presence of three distinct subpopulations, which were subsequently annotated as left, middle, and right subpopulations based on Monocle^81,82^ trajectories. Genes that are differentially expressed in the three subpopulations were identified using Seurat *FindMarkers* function with the default Wilcoxon test.

Cortex cells from Arabidopsis, *E. salsugenium*, and *S. parvula* were mapped to the selected reference, and assigned to reference left, middle, and right subpopulation using *MapQuery* in Seurat^22^. Under- and over-representation of each subpopulation in these three species compared to *S. irio* were determined by hypergeometric test using R function *phyper* with lower.tail = TRUE and lower.tail = FALSE, respectively. The resulting *p*-values were corrected for multiple testing using the Benjamini-Hochberg procedure in R function *p.adjust*.

To examine the expression of flavonoid biosynthesis genes in other species with more than one layer of cortex, we’ve retrieved published scRNA-seq data for rice^33^ and Medicago^34^. The annotated cortex cells were extracted from these dataset and re-analyzed using Seurat^22^. The orthologs of flavonoid biosynthesis genes from these two species were identified by comparing their proteomes to the Arabidopsis proteome using OrthoFinder (v2.5.4)^57^. The expression of the identified flavonoid biosynthesis orthologs were then examined in the extracted cortex population for each species.

### Comparison of basal expression and stress responses among sub-genomes in *C. sativa*

To compare the contribution of sub-genomes to the data integration between Arabidopsis and *C. sativa*, we used the latest refined sub-genome structure for *C. sativa* as well as the refined syntenic orthologs between these two species^48^. The integration was performed separately with all sub-genomes together as well as with each sub-genome separately (Supplementary Fig. 36) using the default settings in IntegrateData from Seurat. For comparison of basal expression and stress responses, we only considered triplets with three complete homeologs. The distribution of DEGs among the three sub-genomes was tested using R function phyper with lower.tail = FALSE. The resulting *p*-values were corrected for multiple testing using the Benjamini-Hochberg procedure in R function p.adjust.

### Identification of fast diverging homeologous groups in *C. sativa*

We identified fast diverging homeologs at the following levels: 1) coding sequences, 2) basal expression level, 3) expression specificity, and 4) stress responses. Divergence at the coding level among homeologs was evaluated using the ω value (dN/dS), which was calculated for each complete triplet using CODEML^83^. Any homeologous groups where either dN or dS was estimated to be zero were removed from downstream analyses. Divergence in expression and expression specificity were computed as the coefficient of variation of variance stabilized expression derived from DESeq2^79^ and Tau indexes of the homeologs within each homeologous group. Similarly, we used the standard deviation of the fold changes of the homeologs as the estimation for the divergence in stress responses for each triplet. For each of these categories, only homeologous groups that fall in the top one percentile of the metrics, regardless of cell types, were considered as the fast diverging homeologous groups.

### Gene ontology analysis

Enrichment analysis of biological pathways on gene sets of interest was performed using BiNGO^84^. Gene ontology (GO) annotations for Arabidopsis were retrieved from the GO consortium (http://geneontology.org/) on December 12^th^, 2021. For non-Arabidopsis species, this analysis was conducted either by replacing gene IDs with their Arabidopsis ortholog IDs (for *E. salsugenium*, *S. irio*, and *S. parvula*) or annotating genes with GO annotations transferred from Arabidopsis (for *C. sativa*). Redundant GO terms with > 50% overlap with similar terms were further clustered using Markov clustering implemented via GOMCL^85^ (https://github.com/Guannan-Wang/GOMCL).

### Whole-mount *in situ* hybridization

Seeds were stratified and then germinated on sterile 1x MS standard media (1X Murashige and Skooge nutrients (Caisson MSP01), MES hydrate 0.5g/L, 1% sucrose and 0.7% agar (Daishin CAS# 9002-18-0) at pH 5.7). The plates were incubated vertically in growth chambers (Percival Scientific CU-36L4) at 22°C with a 16/8 hour light/dark cycles and grown until the roots were ∼2-3 cm in length (5 days for Arabidopsis, 4 days for *S. irio*, 4-5 days for *S. parvula*, and 6-7 days for *E. salsugineum*).

Probes for selected candidates were designed and synthesized by Molecular Instruments (https://www.molecularinstruments.com). Sample fixation, dehydration, permeabilization, protease digestion, probe hybridization and amplification were performed as described in Oliva *et al*.^31^ with the following modifications: after hybridization and washes, roots with were left in ClearSee^86^ overnight and then stained with 0.22% calcofluor solution (Flourescent Brightener 28, Santa Cruz Biotechnology sc-218504) to demarcate cell layers. Samples were mounted on glass slides and imaged with SP8 Leica Confocal Microscope, using a 20X oil immersion objective. Calcofluor white and Alexa 647-coupled amplifiers were excited at 405 nm and 640 nm, and collected at 425-475 nm and 650-675 nm, respectively. Leica LAS X Navigator was used to collect and merge tiled images wherever the differentiated zone was included. Images were processed using Fiji^87^.

## DATA AVAILABILITY

All data are available in the main text, the Supplemental Information, referenced databases and public databases. All raw sequence data generated in the current study have been deposited at NCBI under BioProject PRJNA1113801. All processed data can be found at NCBI GEO database under accession code GEO: GSE268881. A web browser to interactively explore the data from the current study is available at: https://plantbiology.shinyapps.io/brassicaceaeatlas.

## CODE AVAILABILITY

All software used in this study is publicly available as described in the Methods. Detailed parameters used for analyzing each type of sequencing data have been described in the Methods. All original code used throughout the data analysis is publicly available at the following GitHub repository: https://github.com/Guannan-Wang/BrassicaceaeAtlas.

## ACKNOWLEDGEMENTS

This work was supported by the Department of Energy (BER-DE-SC0020358 to J.R.D., J.S., and M.D.; BER DE-SC0022985 to J.R.D and M.D.), the US National Science Foundation (NSF-BSF-IOS-EDGE 1923589/2019610 to M.D.), and Howard Hughes Medical Institute Investigator award (JRD). Single-cell library prep and next-generation sequencing was carried out in the Advanced Genomics Core at the University of Michigan, supported by the National Cancer Institutes of Health Award P30CA046592 to J.S..The authors acknowledge Dr. Song Li and his lab members at Virginia Tech for sharing feedback during group discussions. The authors also acknowledge the LSU High Performance Computing services and Stanford Research Computing Center for providing computational resources needed for data analyses.

## AUTHOR CONTRIBUTIONS

Conceptualization, G.W., J.R.D., J.S., and M.D.; Methodology, G.W., K.H.R., J.R.D., J.S., and M.D.; Software, G.W.; Validation, A.D.; Formal Analysis and Investigation, G.W.; Resources, D-H.O., M.O. and R. L.; Data Curation, G.W.; Writing - original draft, G.W. and M.D.; Writing - Review & Editing, all authors; Visualization, G.W.; Supervision, J.R.D., J.S., and M.D.; Funding Acquisition, J.R.D., J.S., and M.D.

## Supplemental information

**Supplementary Figures 1-39, Supplementary Data 1-10, and Supplementary Movie 1**

**Supplementary Data 1. Statistics of scRNA-seq data generated in the current study.**

**Supplementary Data 2. Known Arabidopsis markers and their expression in the current study.**

**Supplementary Data 3. Markers identified from the current study.**

**Supplementary Data 4. Orthologs with divergent expression across species.**

**Supplementary Data 5. Genes preferentially expressed in different cortex subpopulations in *S. irio*.**

**Supplementary Data 6. Differentially expressed genes from each species in response to ABA and NaCl.**

**Supplementary Data 7. GOs enriched among stress-responsive genes in each cell type from each species.**

**Supplementary Data 8. Stress response patterns among orthologs.**

**Supplementary Data 9. Homeologs exhibiting high divergence in coding sequences and expression.**

**Supplementary Data 10. Comparison of gene expression between bulk and pseudo-bulk samples.**

**Supplementary Movie 1. 3D view of integrated atlas.**

---

**Supplementary Fig. 1.**
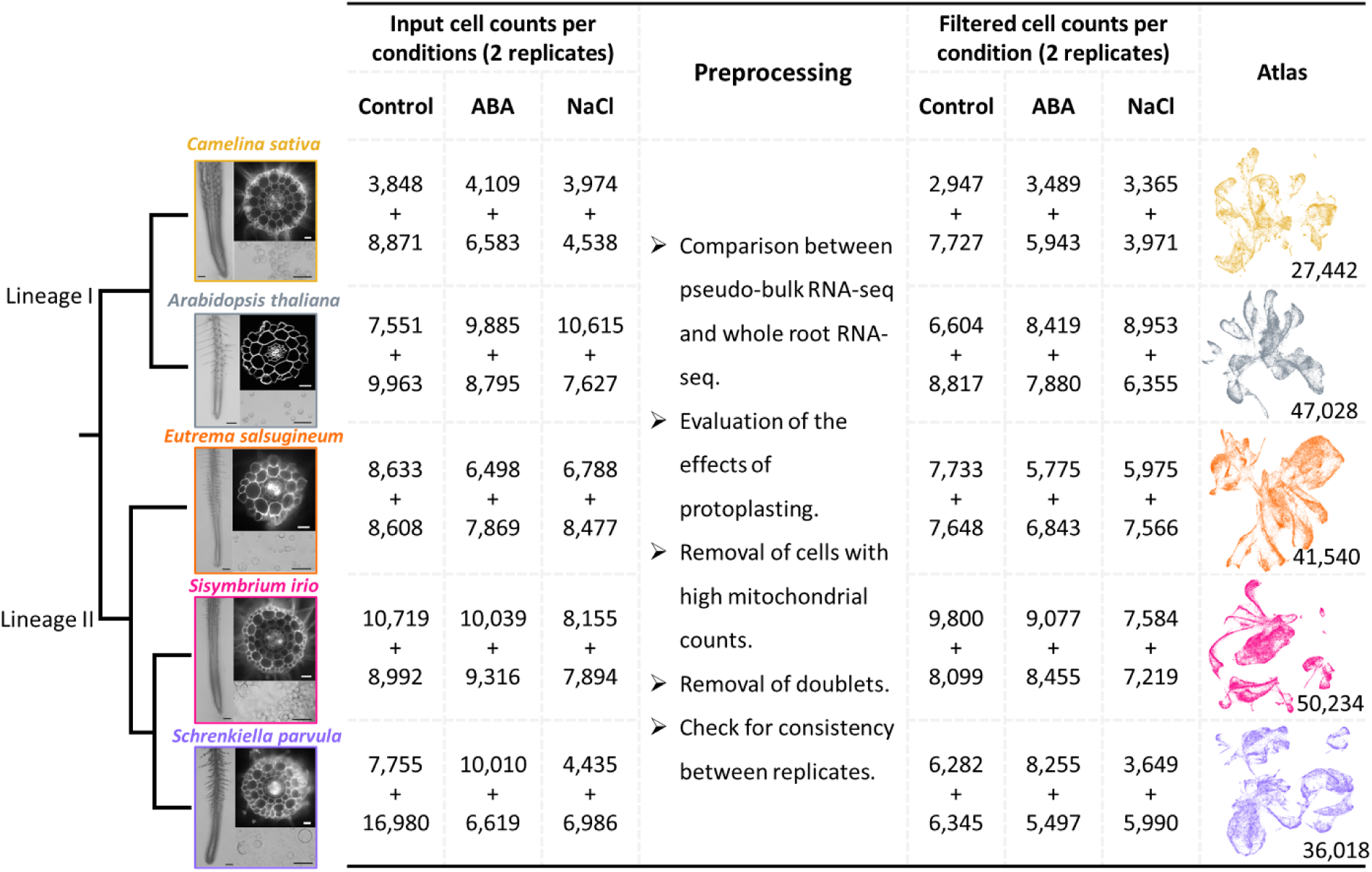
Overview of scRNA-seq dataset for four 5 Brassicaceae species and data preprocessing. The number in each UMAP represents the total number of filtered cells from control, 5 μM ABA- and 100 mM NaCl-treated samples.

**Supplementary Fig. 2.**
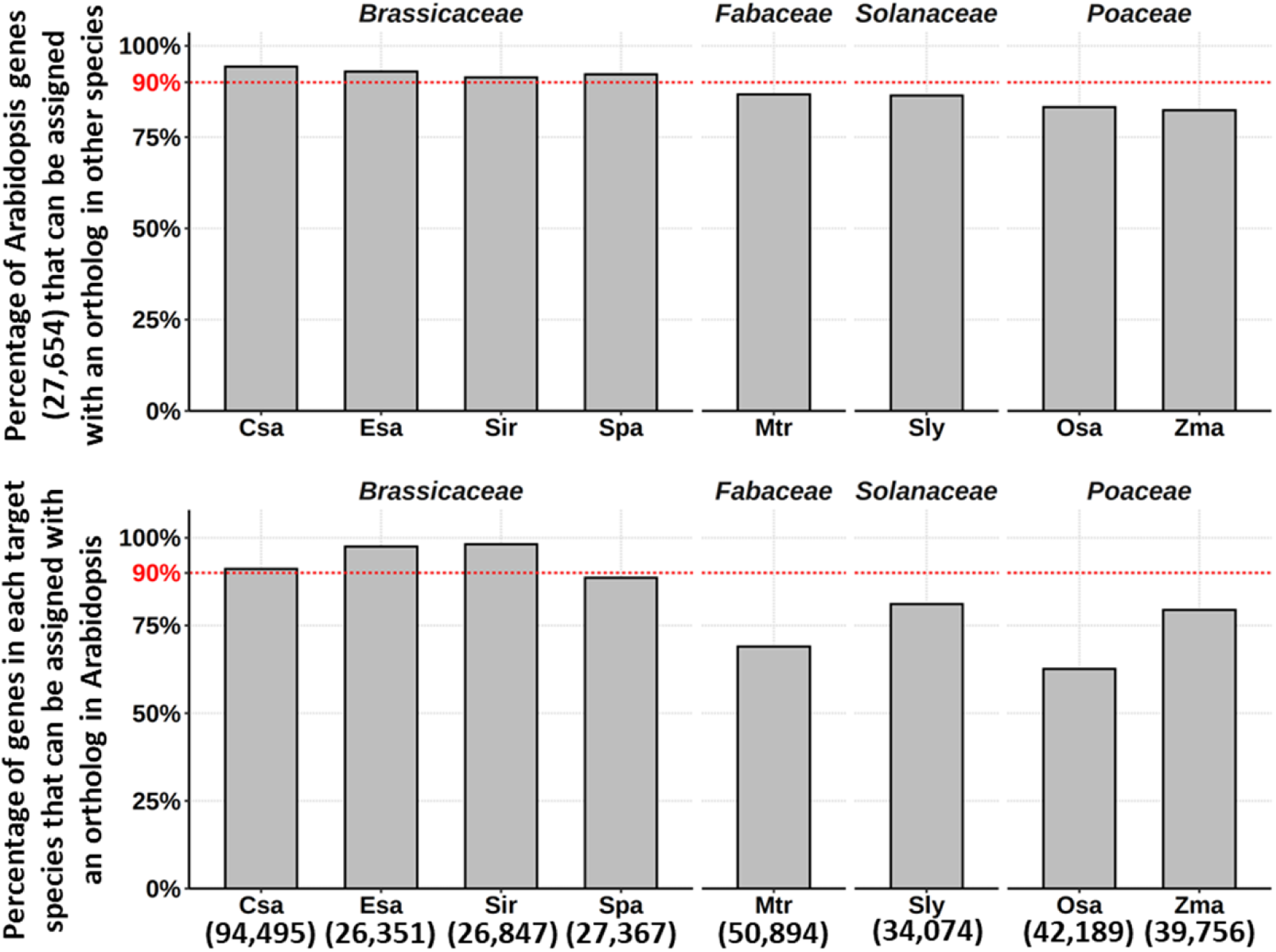
Percentage of genes with orthologs between Arabidopsis and other species. Csa, *Camelina sativa*; Esa, *Eutrema salsugineum*; Spa, *Schrenkiella parvula*; Sir, *Sisymbrium irio*; Mtr, *Medicago truncatula*; Sly, *Solanum lycopersicum*; Osa, *Oryza sativa*; Zma, *Zea mays*. Numbers in parenthesis indicate the numbers of protein-coding genes for each species.

**Supplementary Fig. 3.**
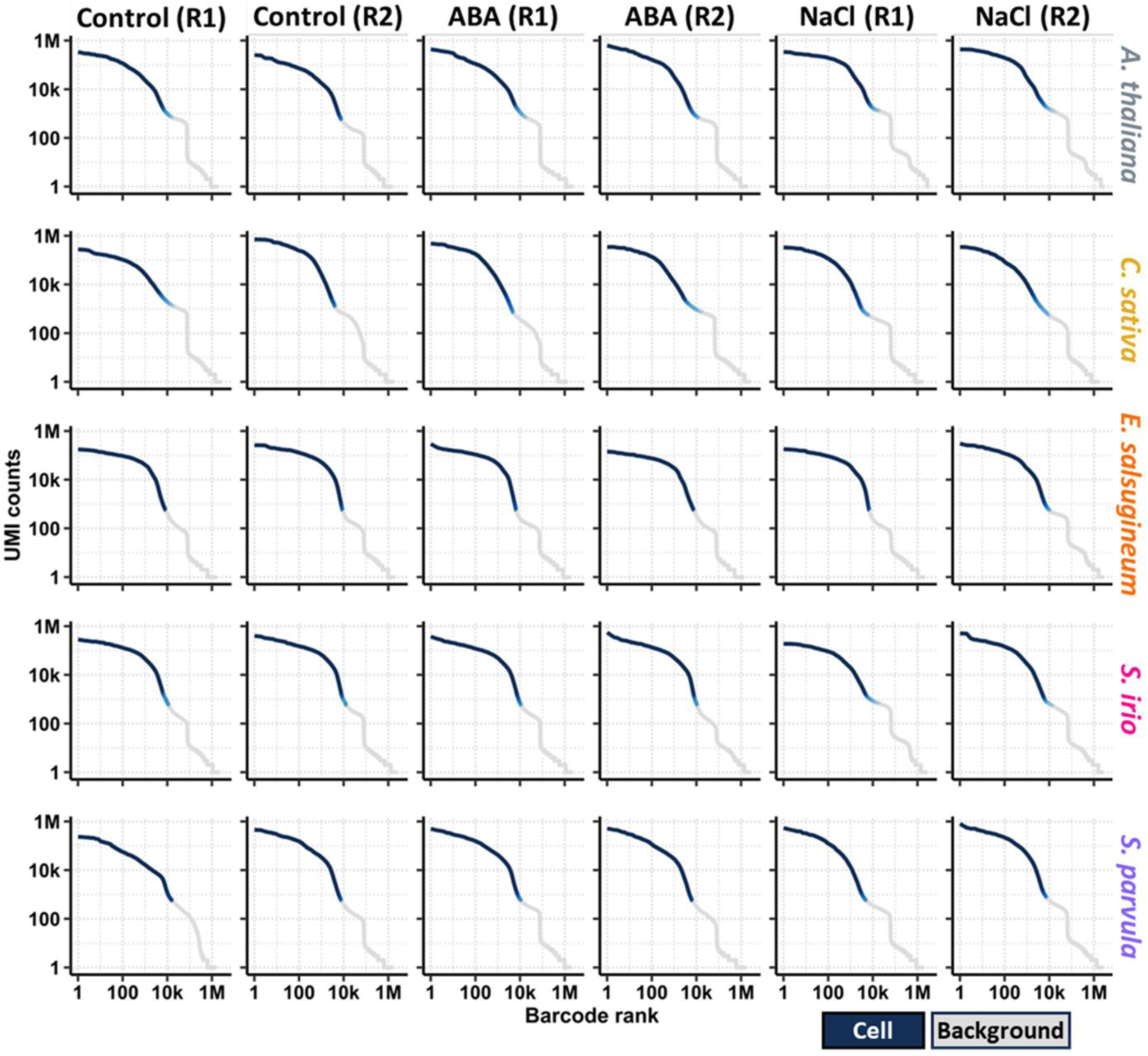
Knee plots for each sample in our study. For each species, samples were harvested from control, 5μM ABA-, and 100 mM NaCl-treated conditions. Two independent replicates (R1 and R2) were used for each condition. The high-confidence cell-associated barcodes were clearly separated from the background ones.

**Supplementary Fig. 4.**
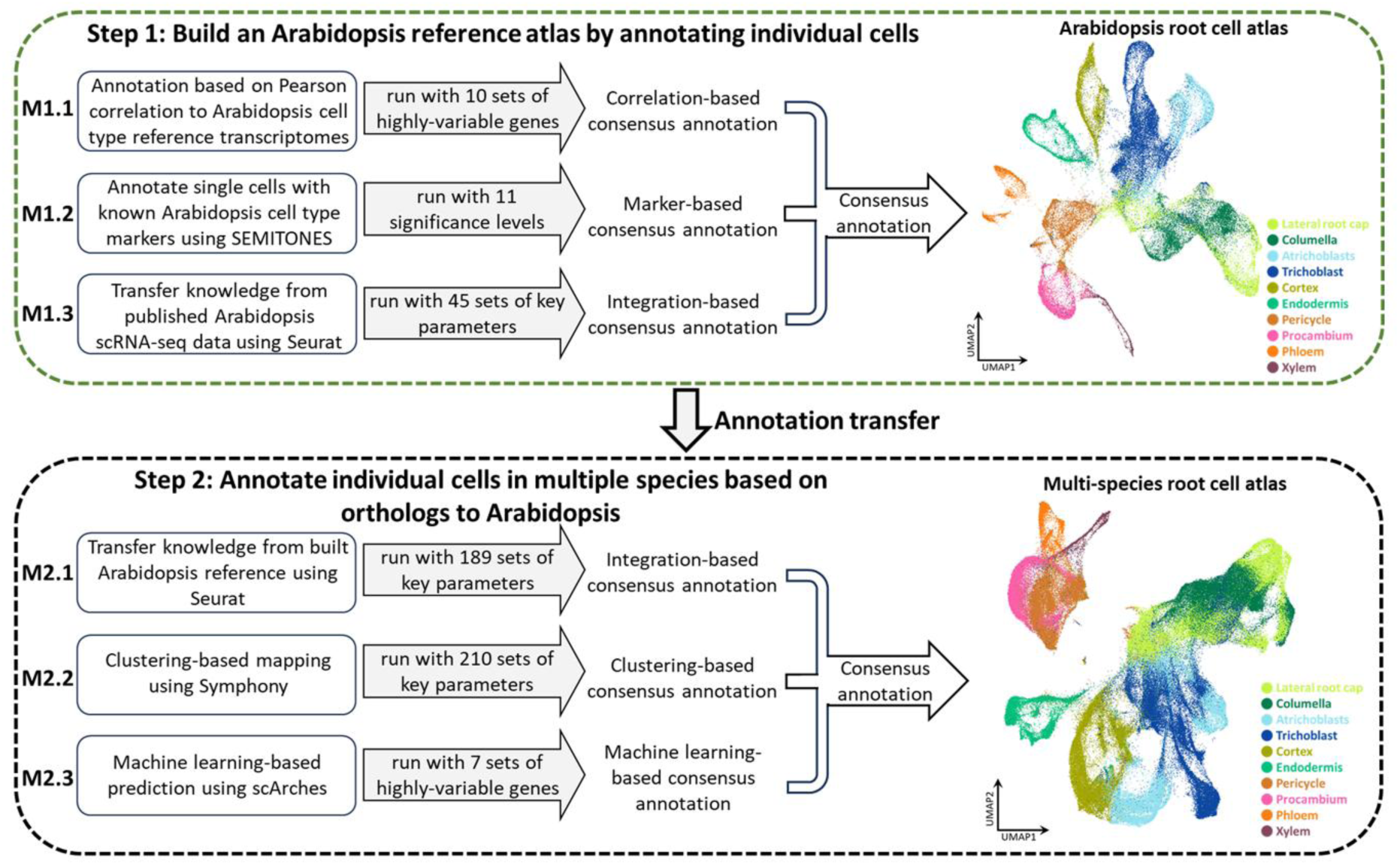
A two-step strategy for cross-species cell type annotations using three independent methods. The Arabidopsis cell type reference transcriptomes were from Li *et al.*, 2016^64^ and Brady *et al.*, 2007^65^. The different sets of highly-variable genes were obtained by categorizing them into different percentile bins. The key parameters for each method were selected based on the tools used, see details in Methods.

**Supplementary Fig. 5.**
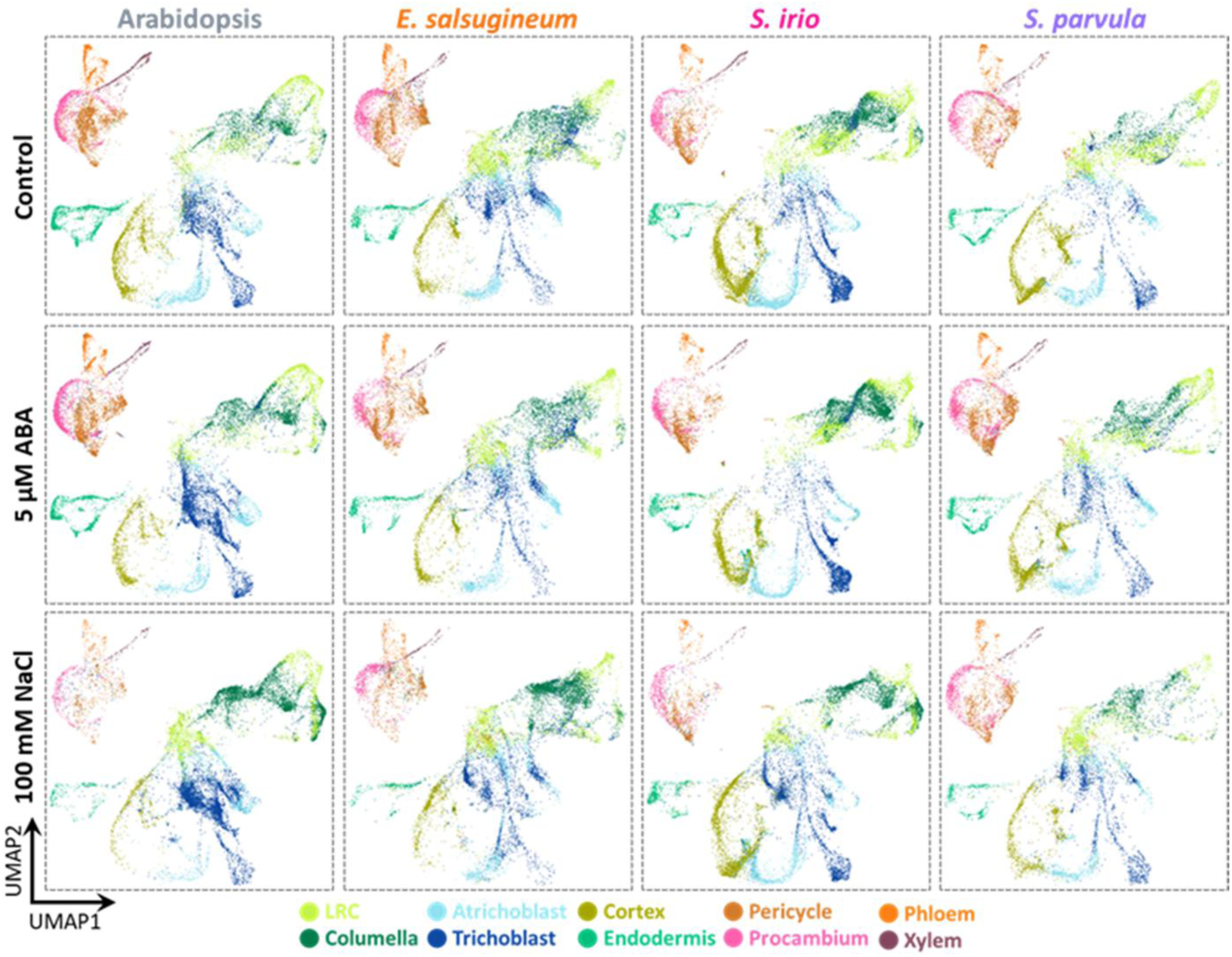
Root cell atlas for each species under different conditions on the integrated UMAP space. All ten root cell types were detected in each species under different conditions. LRC, Lateral Root Cap.

**Supplementary Fig. 6.**
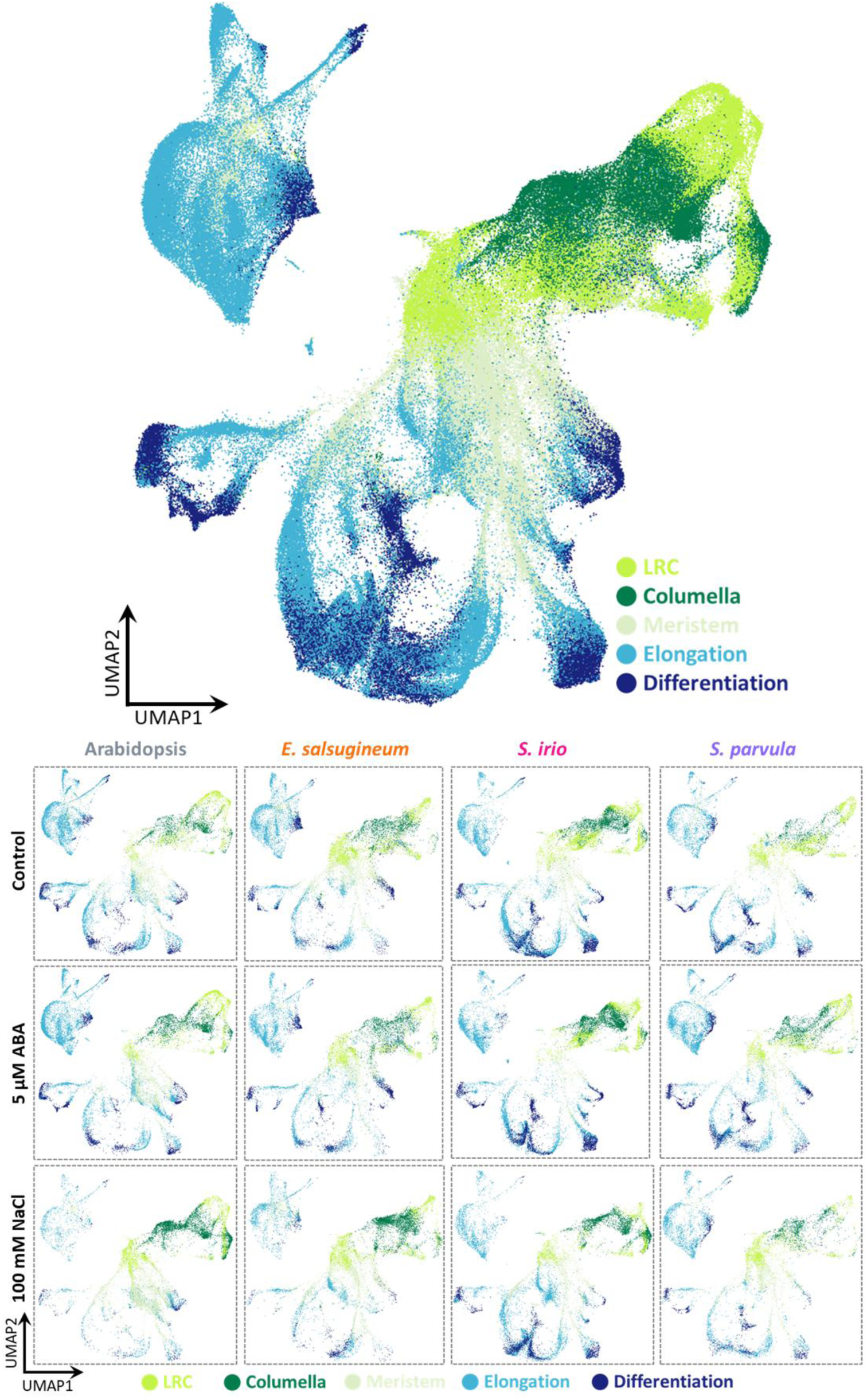
Developmental zone annotation for the integrated root cell atlas from four species (top) and in each species (bottom). All developmental zones were detected in each species under different conditions. LRC, Lateral Root Cap.

**Supplementary Fig. 7.**
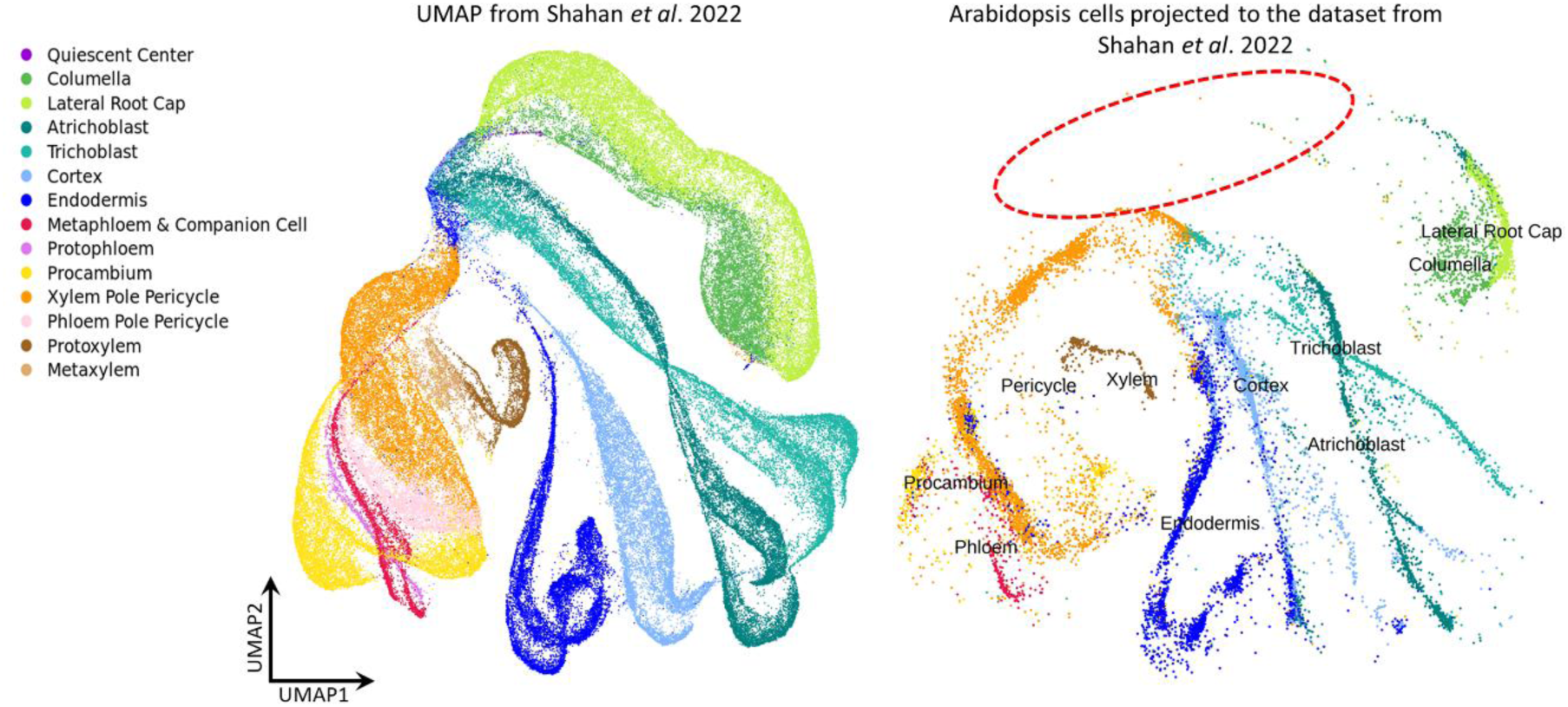
Projection of Arabidopsis control cells from current study to Shahan *et al*^21^. The red dashed oval marks the expected space for quiescent center cells, currently not detected in the current dataset for Arabidopsis.

**Supplementary Fig. 8.**
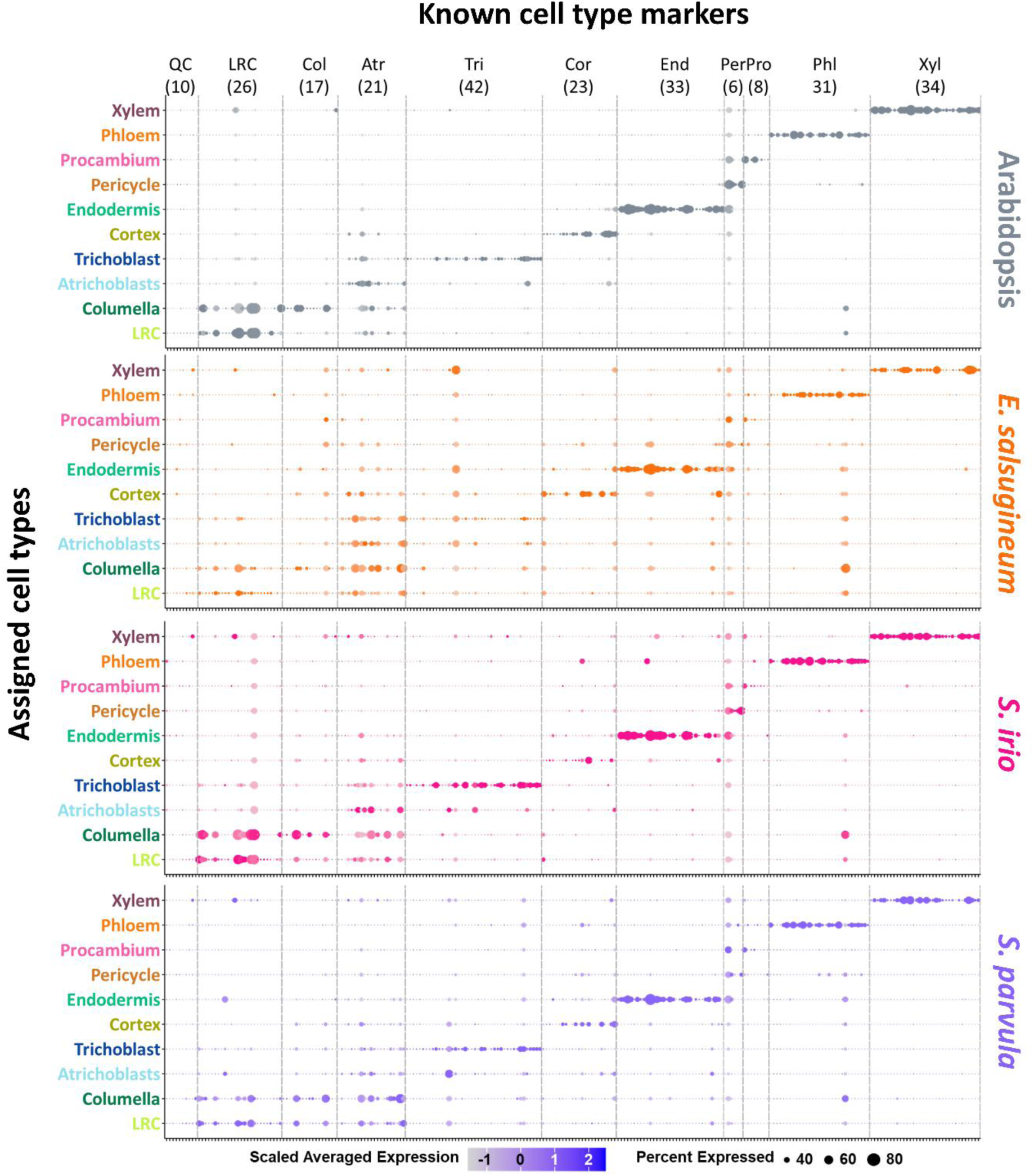
Cell-type expression of known Arabidopsis cell-type markers and their 1-to-1 ortholog in other species under control condition. See Supplementary Data 2 for the list of markers used. Dot size represents the percentage of cells in which each gene is expressed. Dot color intensity indicates the normalized expression of each gene in each cell type. QC, Quiescent center; LRC, Lateral Root Cap; Col, Columella; Atr, Atrichoblast; Tri, Trichoblast; Cor, Cortex; End, Endodermis; Per, Pericycle; Pro, Procambium; Phl, Phloem; Xyl, Xylem.

**Supplementary Fig. 9.**
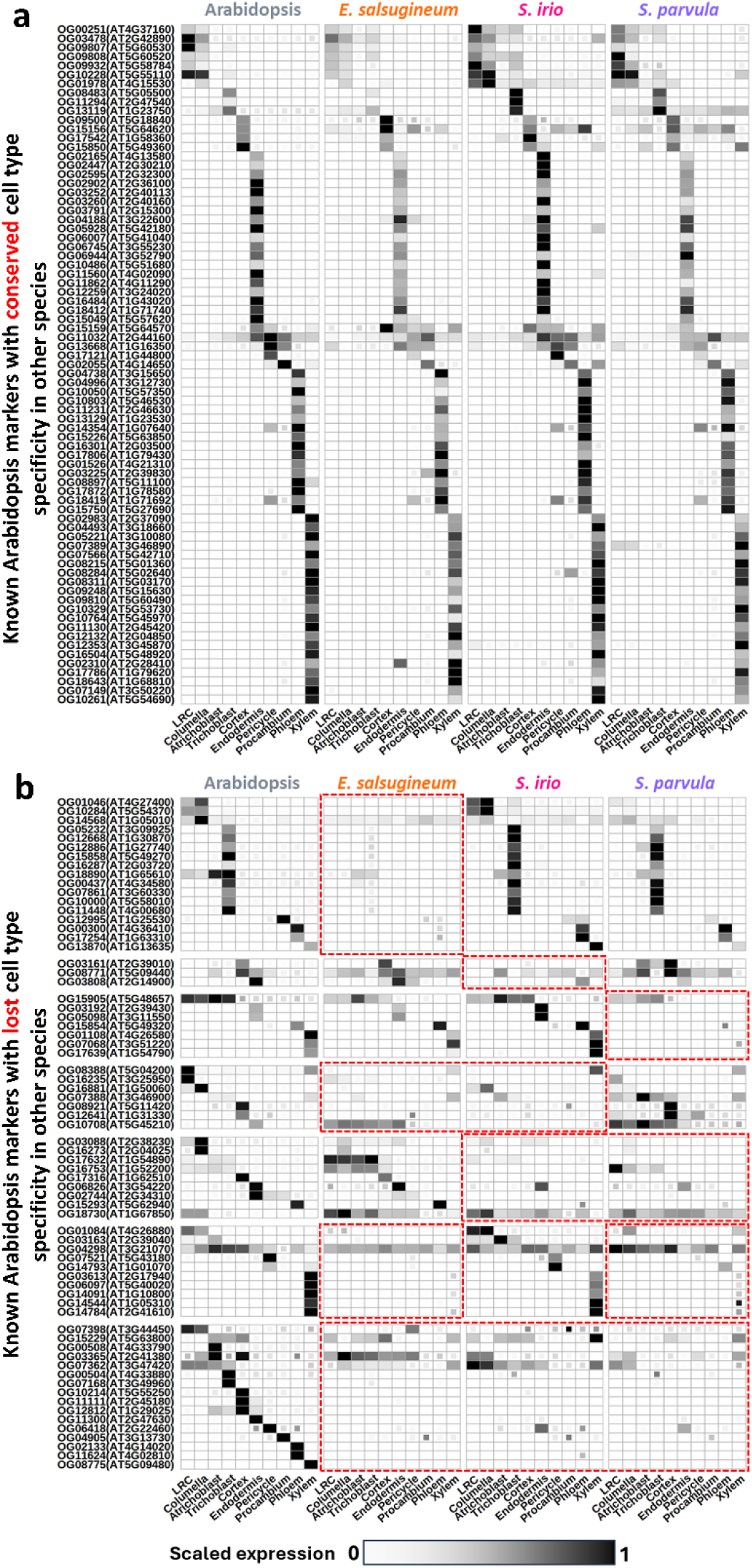
Expression of conserved (a) and diverged (b) Arabidopsis cell type markers across species under control condition. Red boxes mark the species where these cell type markers lost their expression specificity. Tiles with incomplete fill indicate these genes were expressed in less than 10% of cells from the target cell types. They were not considered for marker identification.

**Supplementary Fig. 10.**
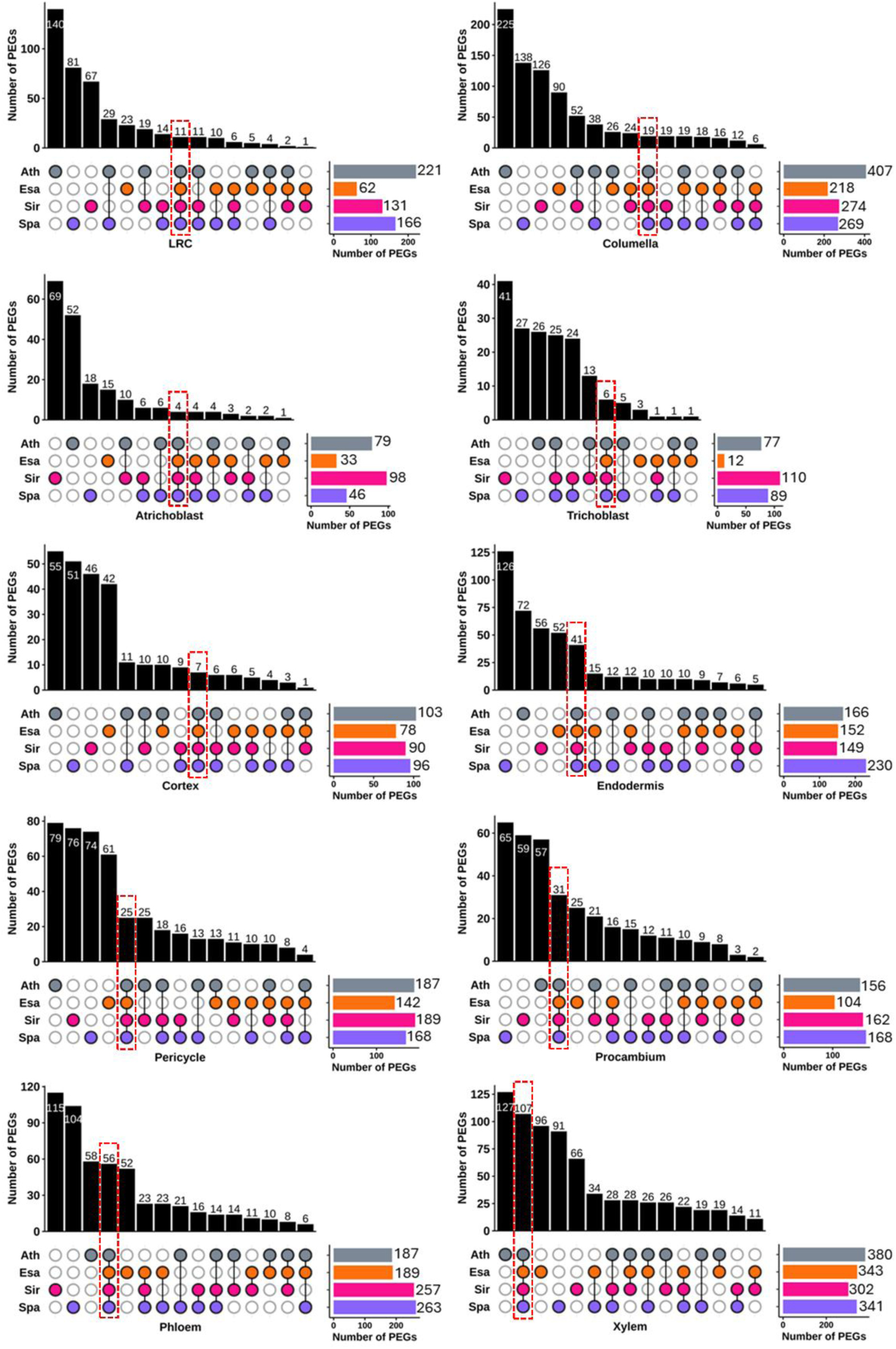
Identification of cell type preferentially enriched genes (PEGs) that were conserved across species for each cell type in the current study. Ath, Arabidopsis; Esa, *Eutrema salsugineum*; Spa, *Schrenkiella parvula*; Sir, *Sisymbrium irio*. The conserved PEGs for each cell type are highlighted with red boxes.

**Supplementary Fig. 11.**
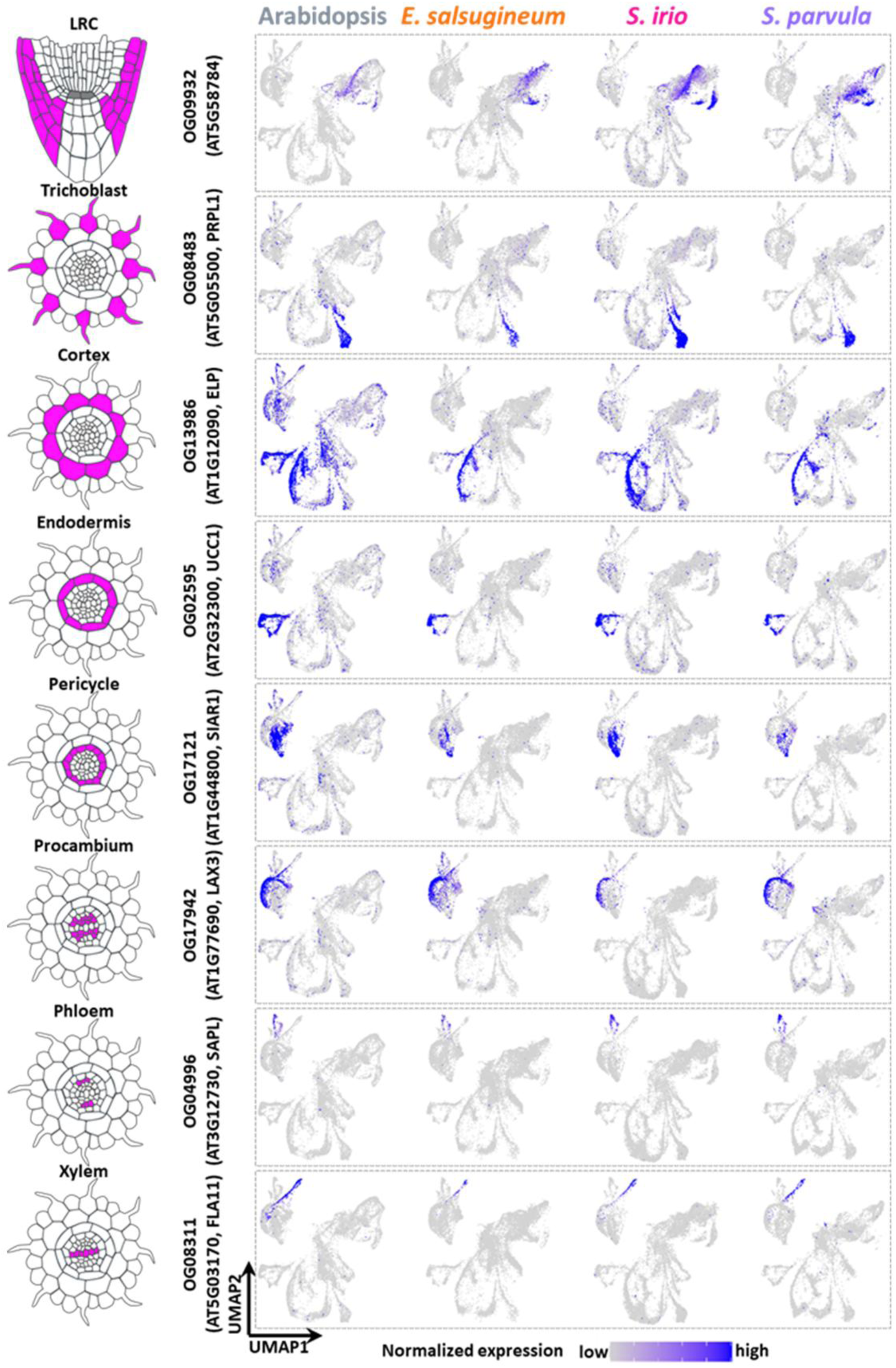
Expression of selected cell type markers in Fig. 2 across species. The expression shown is from the control samples. PRPL1, proline-rich protein-like 1; ELP, extensin-like protein; UCC1, uclacyanin 1; SIAR1, siliques are red 1; LAX3, like aux1 3; SAPL, Sister of APL; FLA11, FASCICLIN-like arabinogalactan-protein 11.

**Supplementary Fig. 12.**
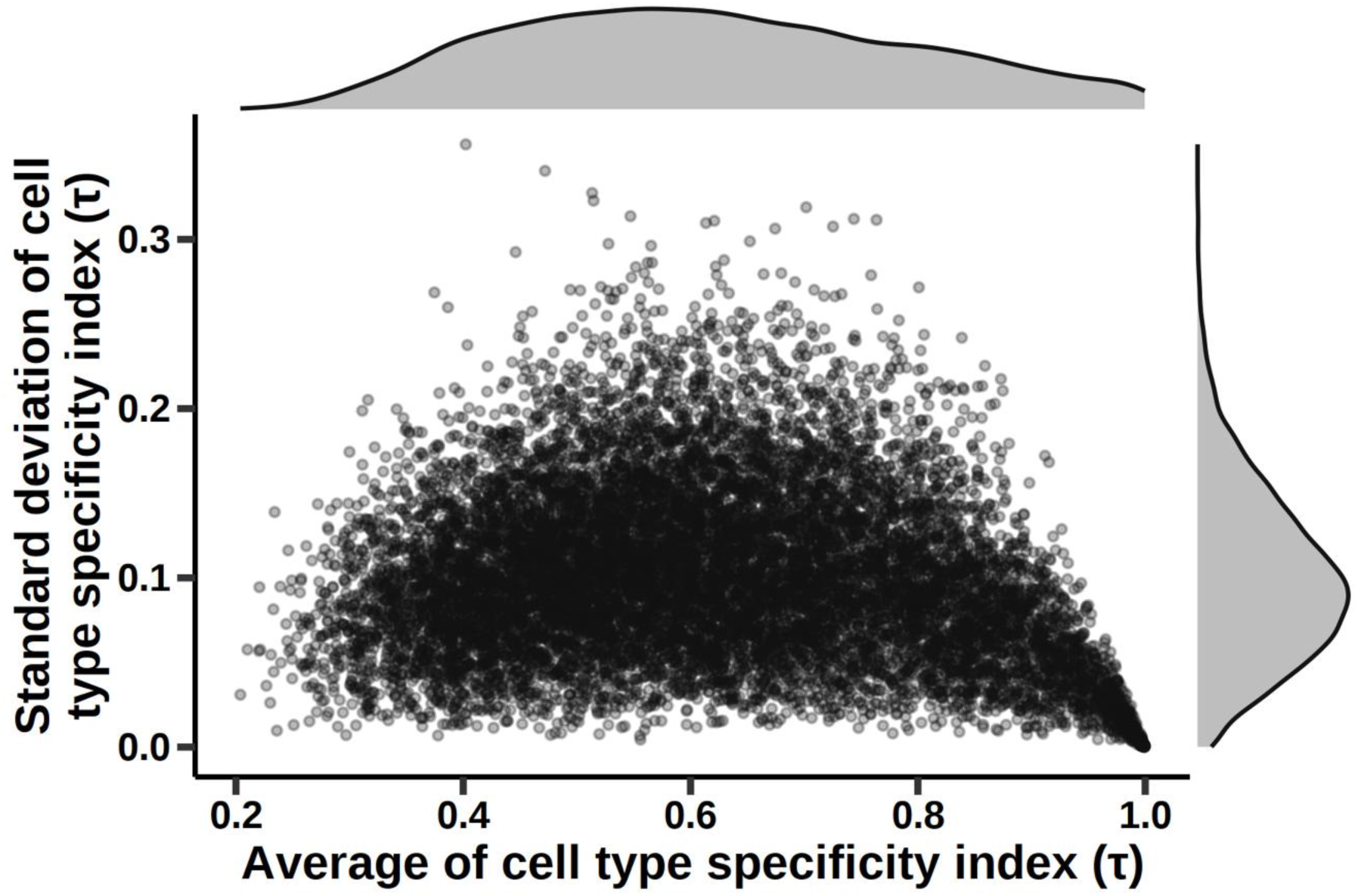
Comparison of the average and the standard deviation of cell type specificity index for the 1-to-1 orthologs. Cell type specificity index (τ) is based on tau metric of tissue specificity in the control samples of each species and calculated as 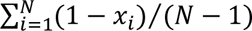, where N is the number of cell types and x_i_ is the expression level in cell type is normalized by the maximal expression of the given gene across all cell types. Cell type specificity index (τ) ranges from 0 to 1, where 0 means broad expression in all cell types and 1 indicates specific expression in one cell type. The cell type specificity indexes for each 1-to-1 orthologs were used to calculate the average and standard deviation for these orthologs.

**Supplementary Fig. 13.**
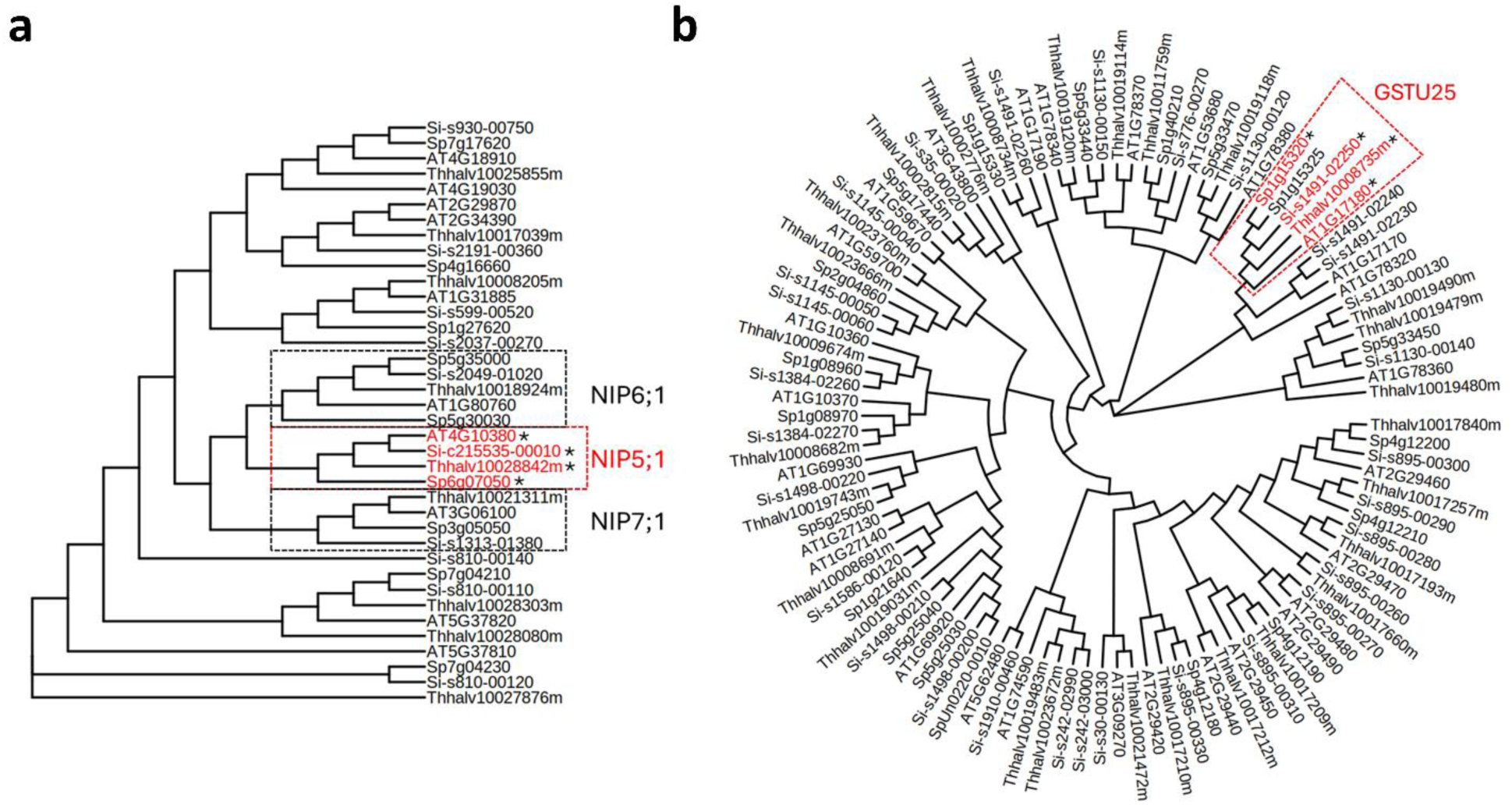
Phylogenetic tree of NIPs (a) and GSTUs (b) from Arabidopsis, *S. irio*, *S. parvula*, and *E. salsugineum*. Genes marked with asterisks are the reciprocal best matches of each other.

**Supplementary Fig. 14.**
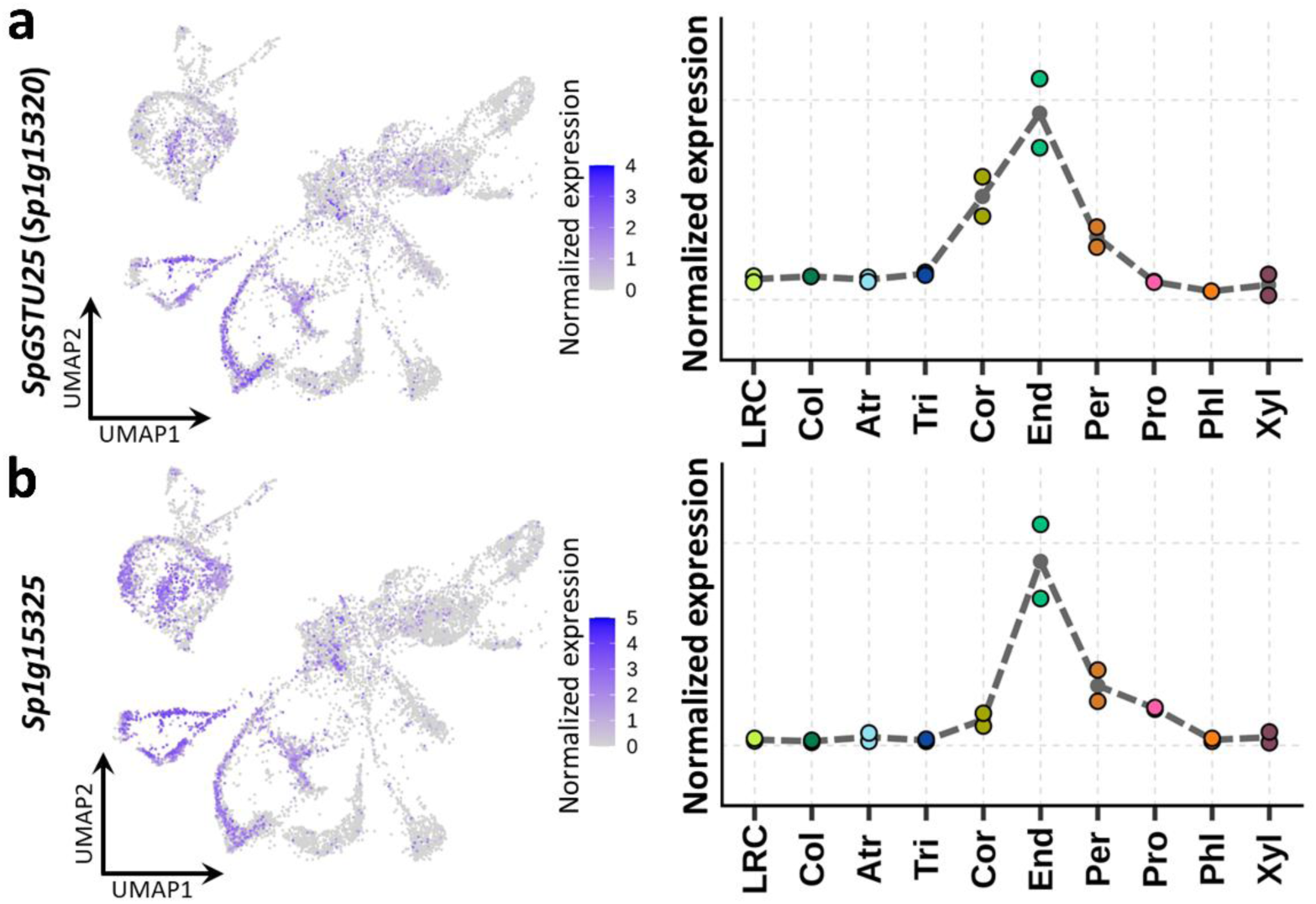
Expression of *SpGSTU25* (*Sp1g15320*) (a) and *Sp1g15325* (b) based on scRNA-seq data for *S. parvula* under control condition. Left panels display the expression of the selected genes visualized on UMAP space. Right panels show the pseudo-bulk cell type expression of the selected genes. LRC, Lateral Root Cap; Col, Columella; Atr, Atrichoblast; Tri, Trichoblast; Cor, Cortex; End, Endodermis; Per, Pericycle; Pro, Procambium; Phl, Phloem; Xyl, Xylem.

**Supplementary Fig. 15.**
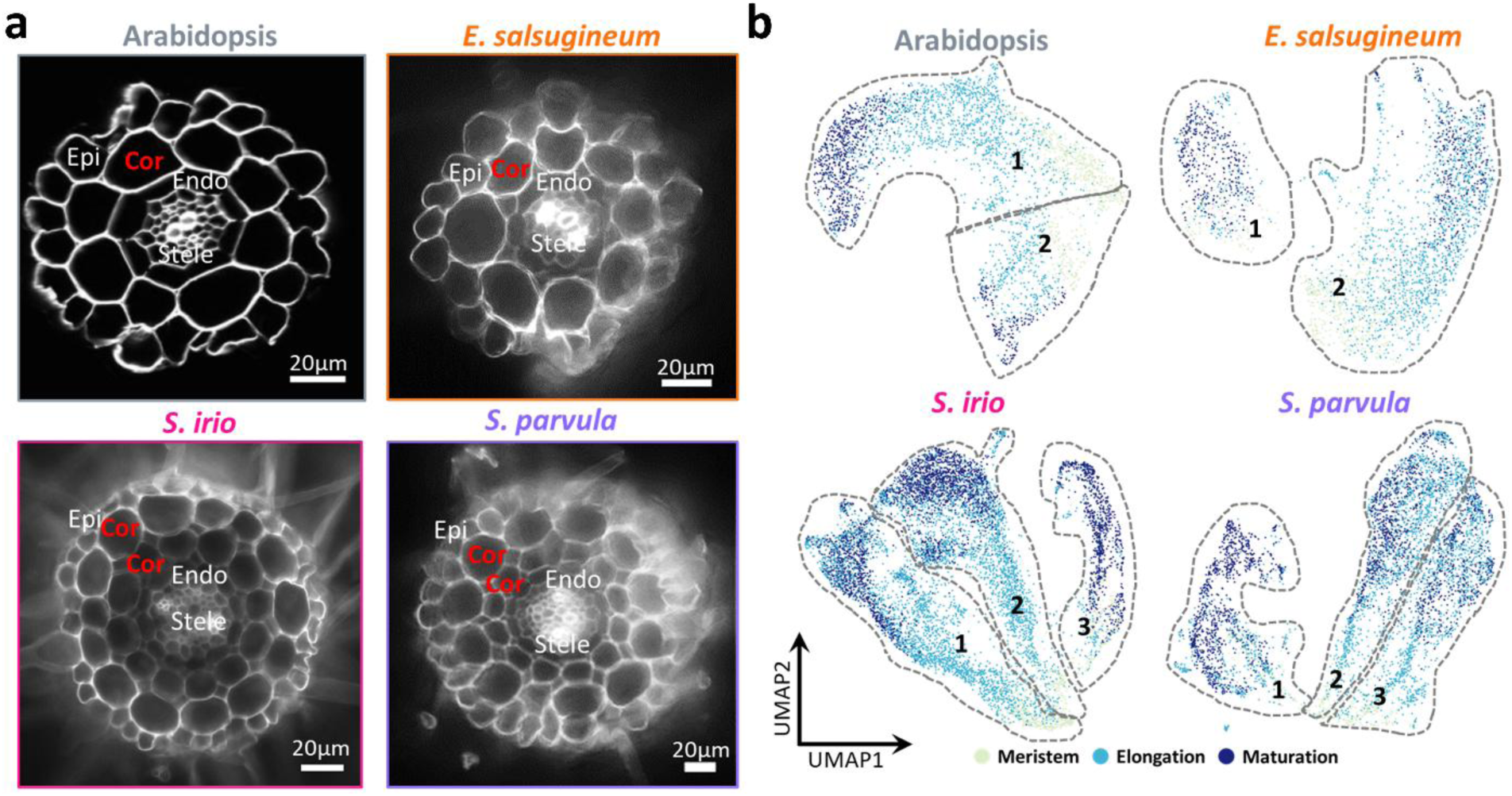
Comparison of cortex cells across species from control samples. a Cross-sections of roots at the maturation zone in each species. Roots were obtained from 5–6-day old seedlings for each species. Scale bars, 20 μm. b UMAP representation of cortex cells in each species. Sub-populations in each species are manually circled based on trajectories identified by Monocle.

**Supplementary Fig. 16.**
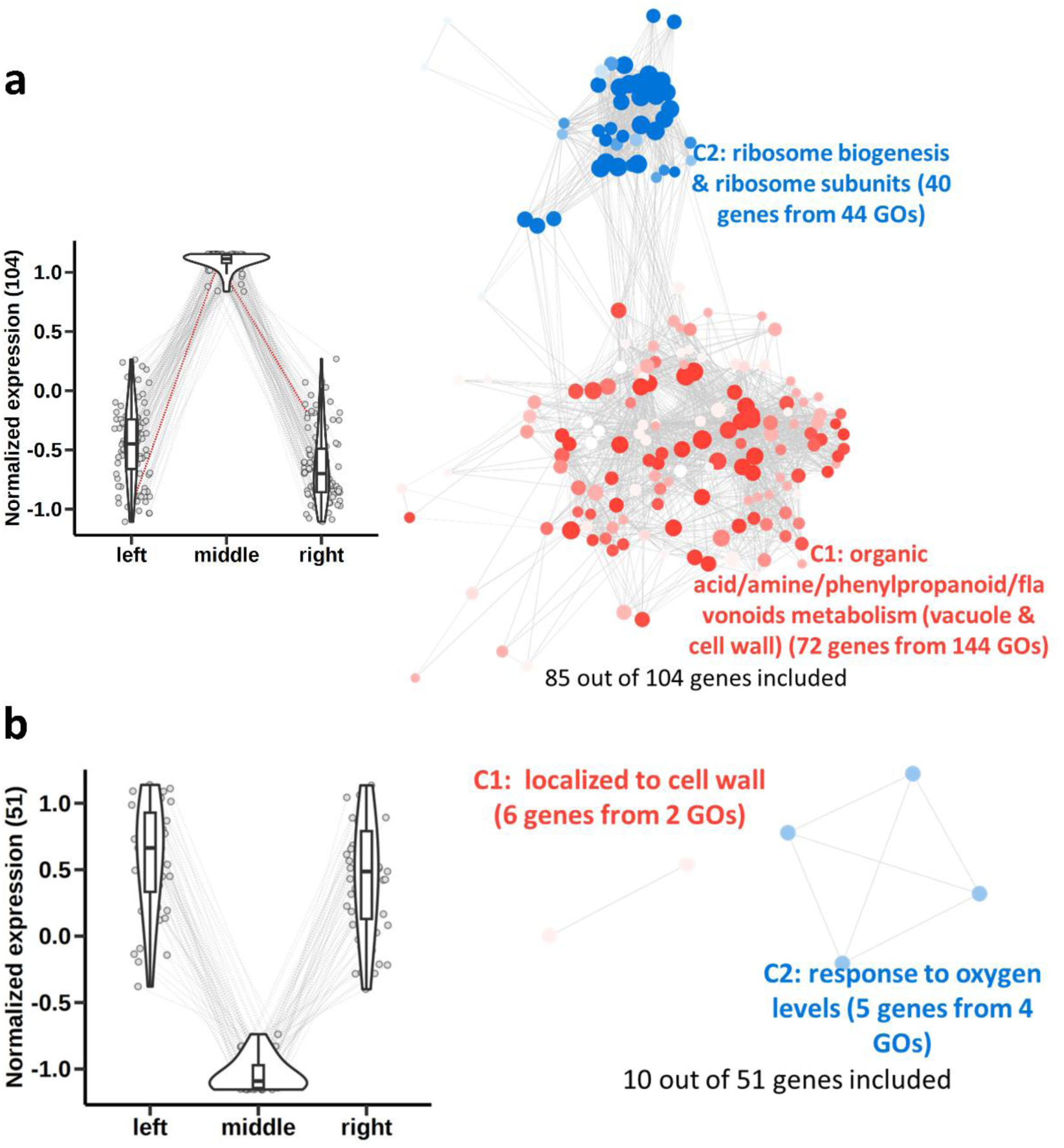
Functional representations of genes that were preferentially expressed (a) and depleted (b) in the middle cortex subpopulation of *S. irio*. Numbers in the parenthesis on the y-axis indicate the number of genes. Box plots represent the median (central value), upper and lower quartile (box limits) with whiskers at ± 1.5 × the interquartile range. The red dash line in the line graph in (a) marks the normalized expression of *Si-s1384-01650*, the HCR result of its expression in *S. irio* is shown in Fig. 4d. GO clusters are differently colored and labeled with the representative functional term. Each node represents a gene ontology (GO) term; node size represents genes in the test set assigned to that functional term; GO terms sharing > 50% of genes are connected with edges; and the shade of each node represents the *p*-value assigned by the enrichment test at adjusted *p* values < 0.05 with darker shades indicating smaller *p*-values.

**Supplementary Fig. 17.**
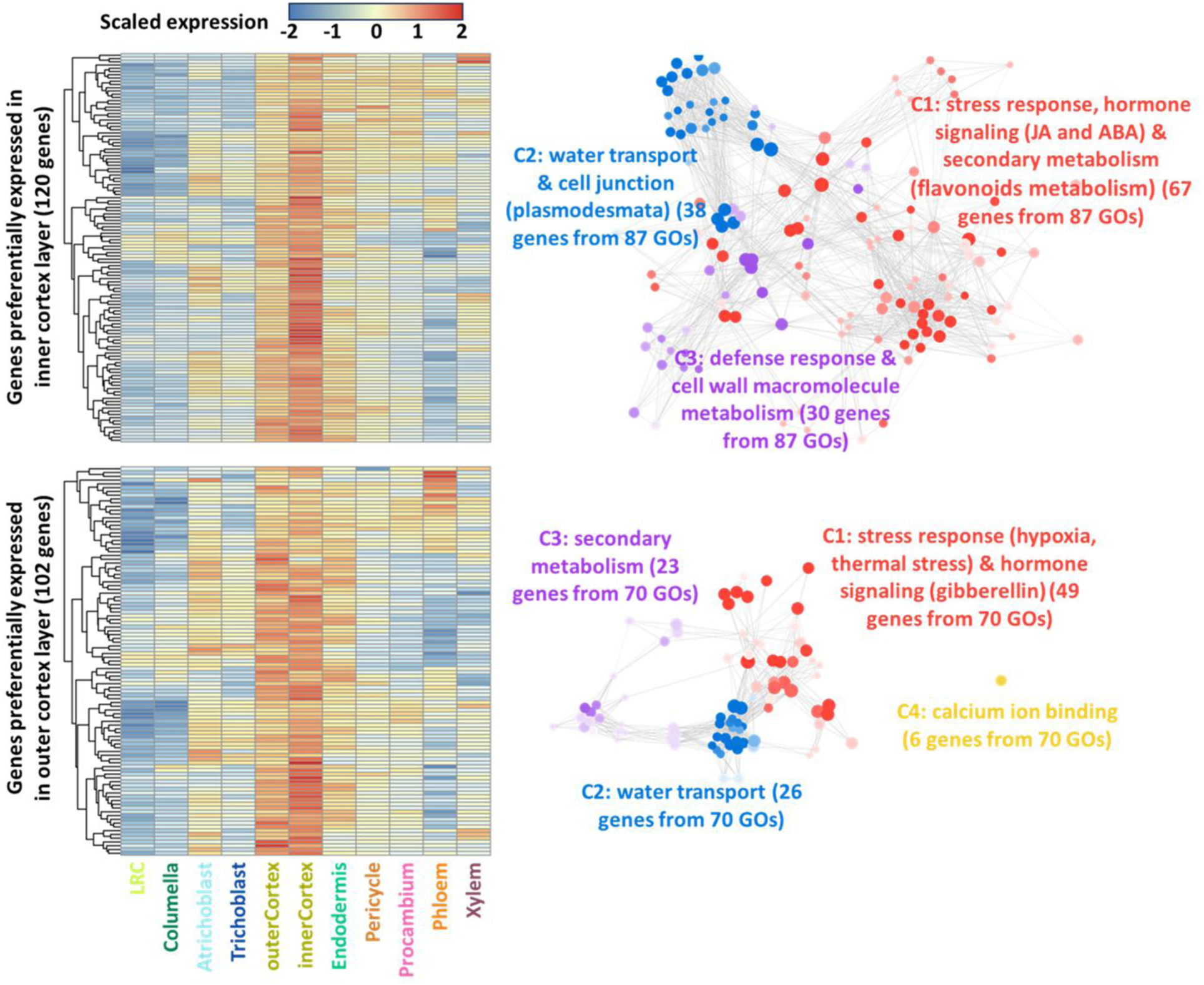
Expression of genes that are preferentially expressed in inner (top) and outer (bottom) cortex layers in *S. irio* and their functional representations. GO clusters are differently colored and labeled with the representative functional term. Each node represents a gene ontology (GO) term; node size represents genes in the test set assigned to that functional term; GO terms sharing > 50% of genes are connected with edges; and the shade of each node represents the p-value assigned by the enrichment test at adjusted p values < 0.05 with darker shades indicating smaller p-values.

**Supplementary Fig. 18.**
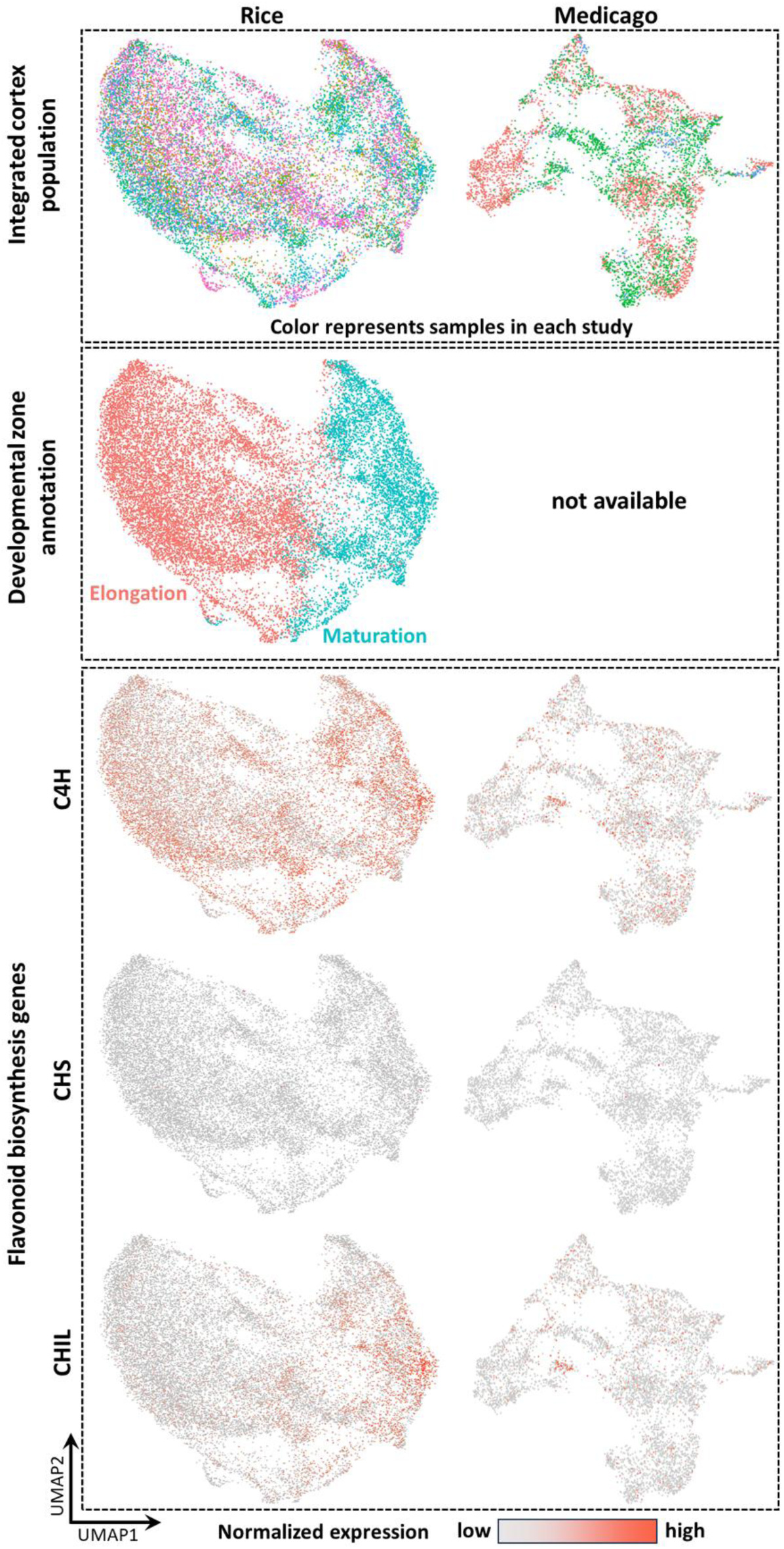
The expression of flavonoid biosynthetic genes in rice and Medicago. The spatial separation of the flavonoid pathway in different cortex layers is not conserved in rice and Medicago. C4H, cinnamic acid 4-hydroxylase; CHS, chalcone synthase; CHIL, chalcone isomerase-like.

**Supplementary Fig. 19.**
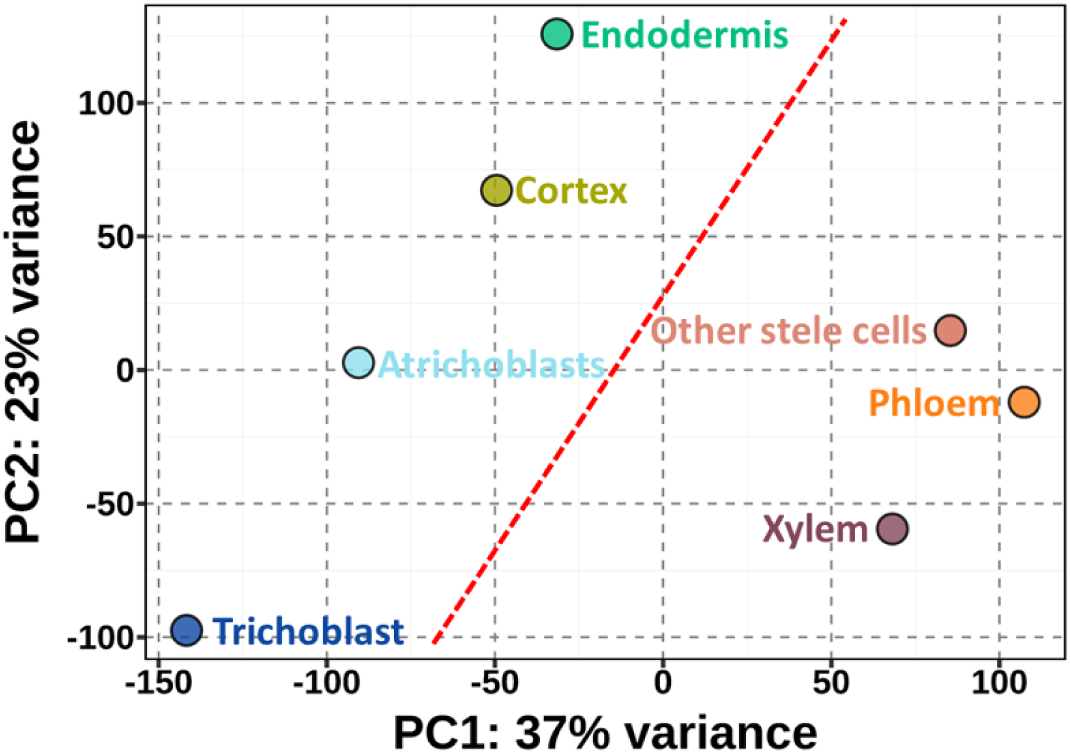
Principal component (PC) analysis of pseudo-bulk cell-type transcriptomes generated from single-nucleus RNA-seq of Arabidopsis (Farmer *et al*., 2021^37^). Each symbol represents a pseudo-bulk cell-type transcriptome. The red dashed lines separate cell type transcriptome subgroups within each species.

**Supplementary Fig. 20.**
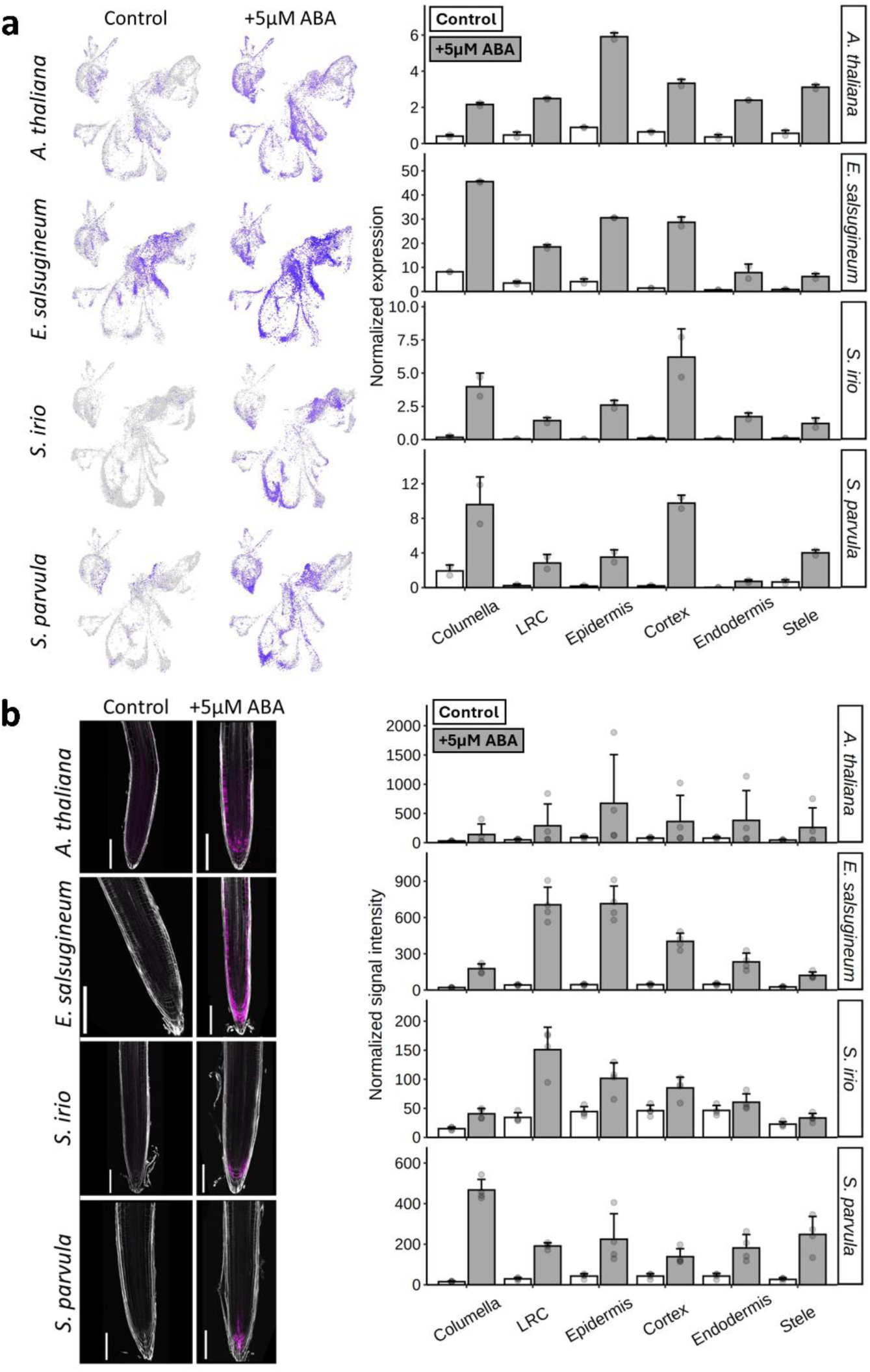
Expression changes of OG04736 orthologs (*AT3G15670*) across species in response to ABA. a Expression of OG04736 orthologs across species from scRNA-seq data under different conditions. Left, UMAP representation of the expression of OG04736 orthologs. Right, pseudo-bulk cell type expression of OG04736 orthologs. b Whole-mount *in situ* HCR detection of expression changes of OG04736 orthologs across species in response to ABA. Left, confocal images of whole-mount HCR showing the detection of OG04736 orthologs across species. Scale bars, 150 μm. Right, average signal intensity of OG04736 orthologs across defined cell type groups.

**Supplementary Fig. 21.**
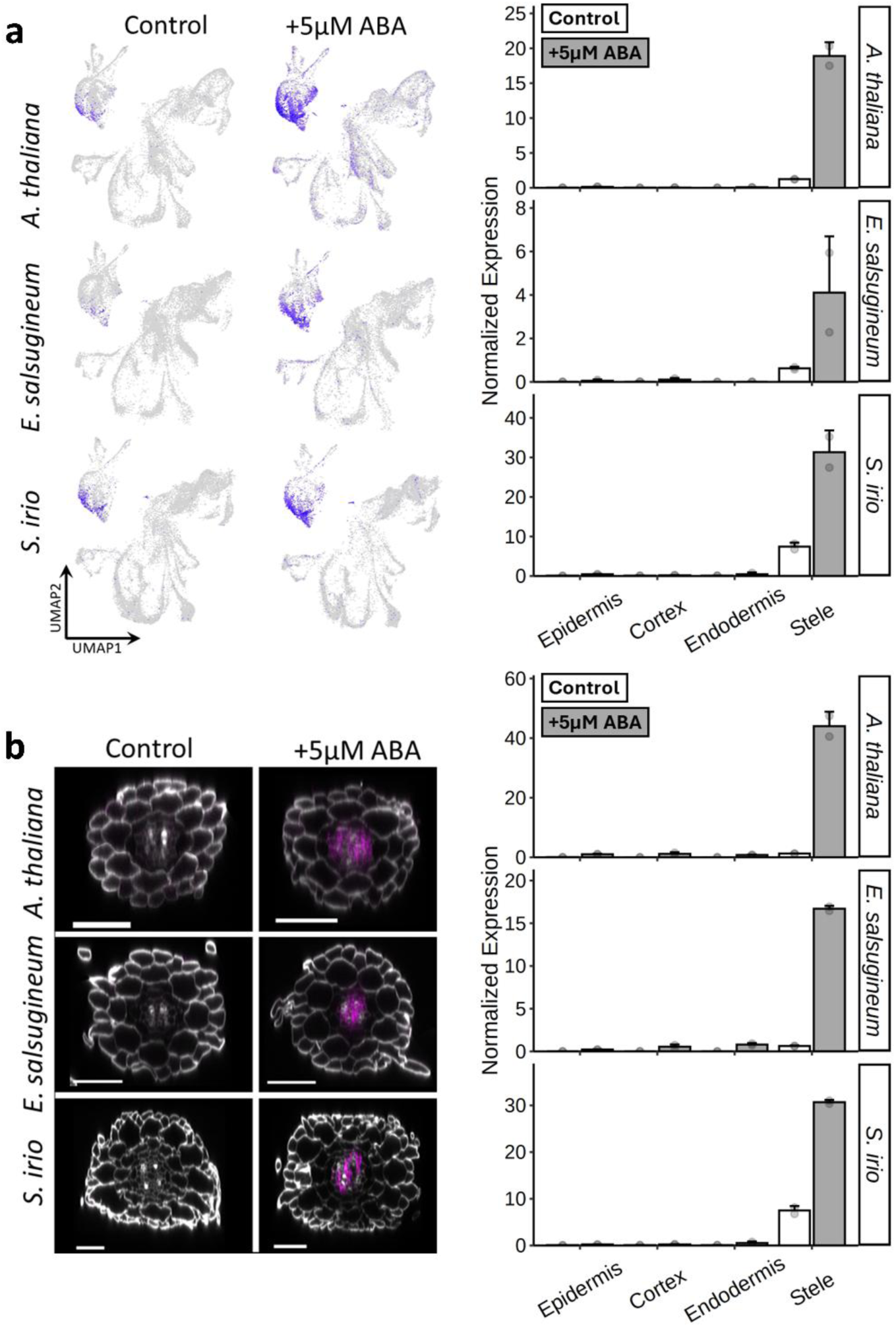
Expression changes of OG03053 orthologs (*AT2G37870*) across species in response to ABA. a Expression of OG03053 orthologs across species from scRNA-seq data under different conditions. Left, UMAP representation of the expression of OG03053 orthologs. Right, pseudo-bulk cell type expression of OG03053 orthologs. b Whole-mount *in situ* HCR detection of expression changes of OG03053 orthologs across species in response to ABA. Left, confocal images of whole-mount HCR showing the detection of OG03053 orthologs across species. Scale bars, 50 μm. Right, average signal intensity of OG03053 orthologs across defined cell type groups.

**Supplementary Fig. 22.**
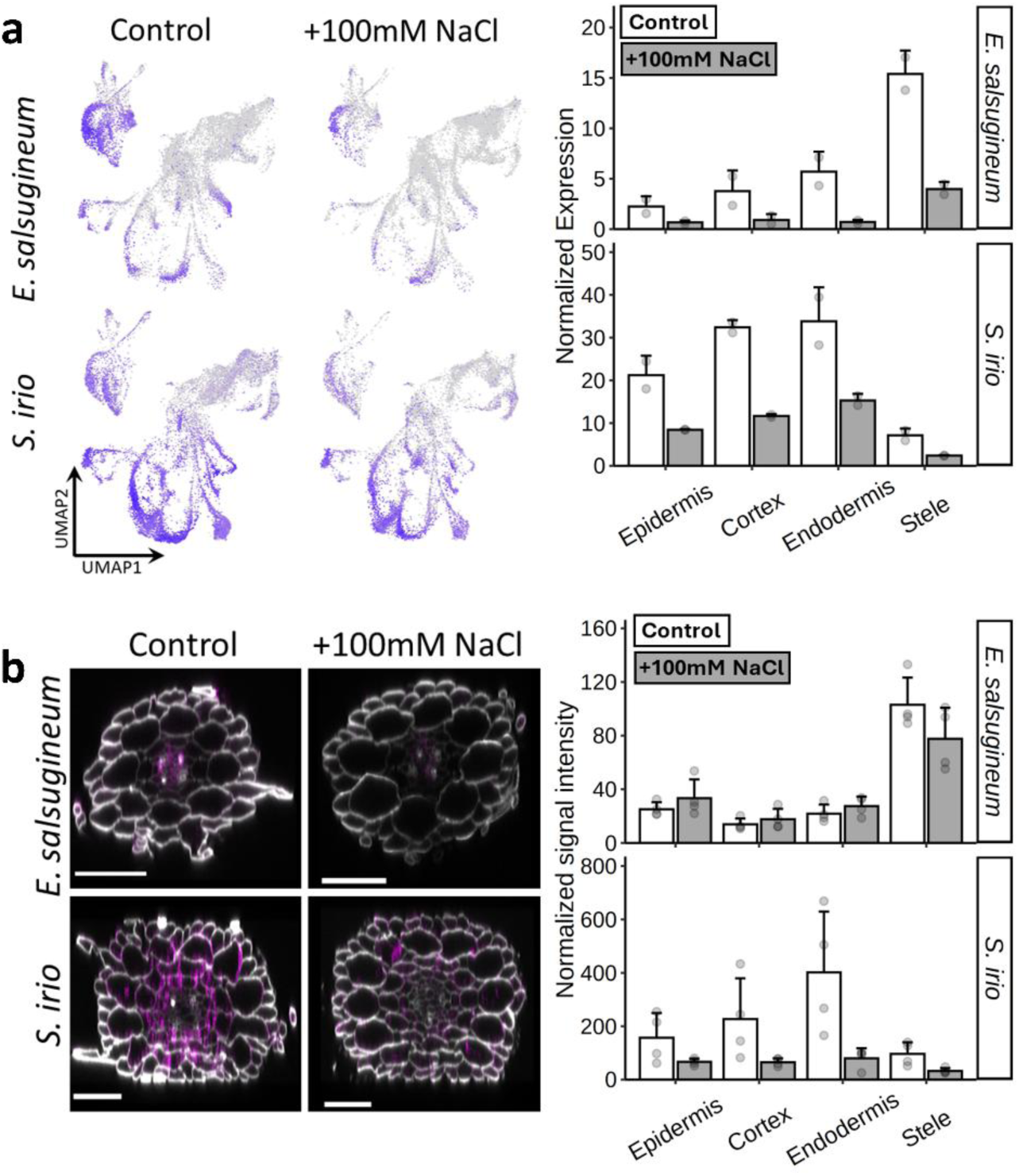
Expression changes of OG10873 orthologs (*AT5G47450*) across species in response to NaCl. a Expression of OG10873 orthologs across species from scRNA-seq data under different conditions. Left, UMAP representation of the expression of OG10873 orthologs. Right, pseudo-bulk cell type expression of OG10873 orthologs. b Whole-mount *in situ* HCR detection of expression changes of OG10873 orthologs across species in response to ABA. Left, confocal images of whole-mount HCR showing the detection of OG10873 orthologs across species. Scale bars, 50 μm. Right, average signal intensity of OG10873 orthologs across defined cell type groups.

**Supplementary Fig. 23.**
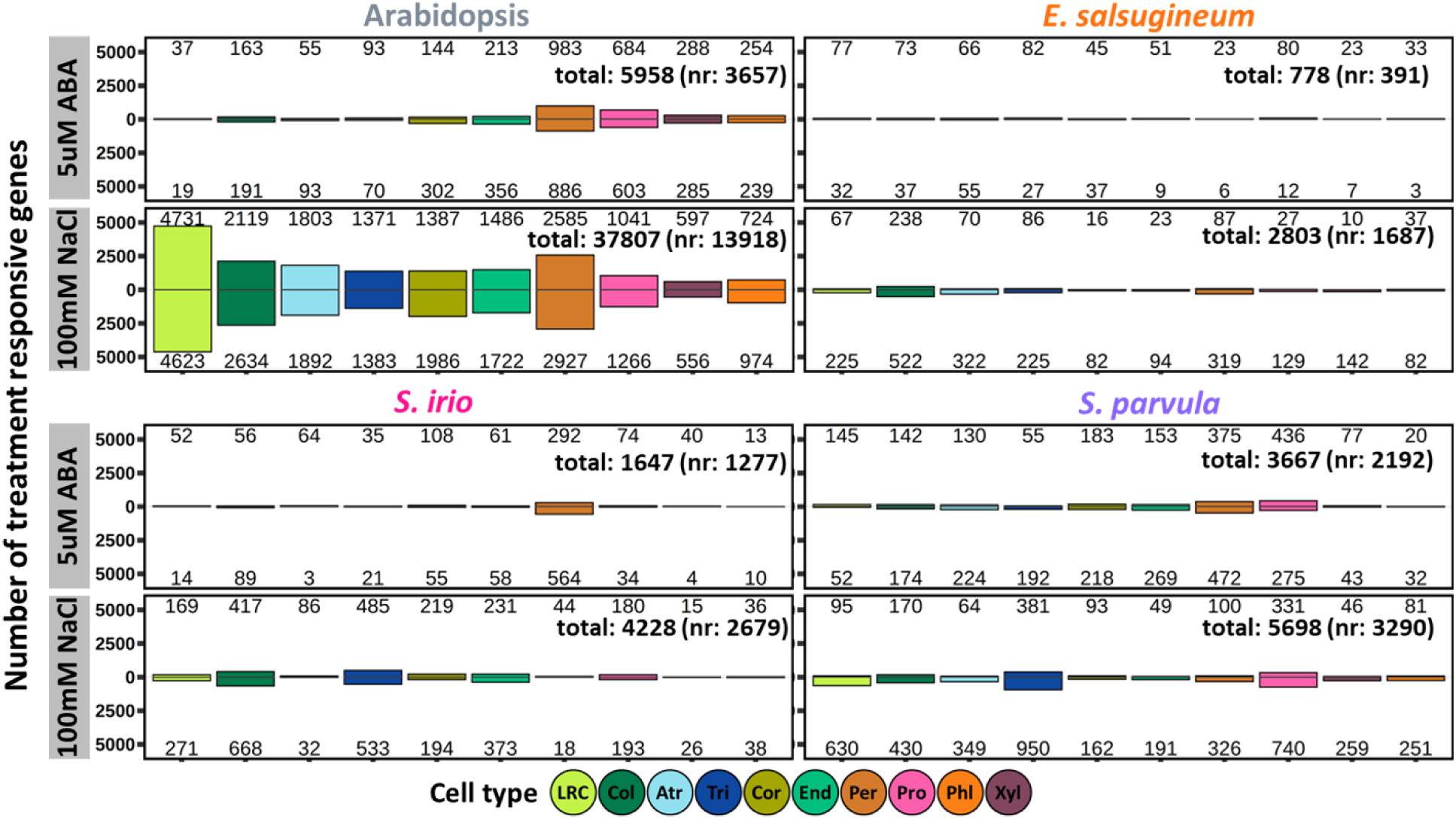
Number of stress responsive genes in each cell type from each species in response to 5 µM ABA and 100 mM NaCl. Total number of stress responsive genes (nr, non-redundant) from all cell types in each species was shown on the top left corner.

**Supplementary Fig. 24.**
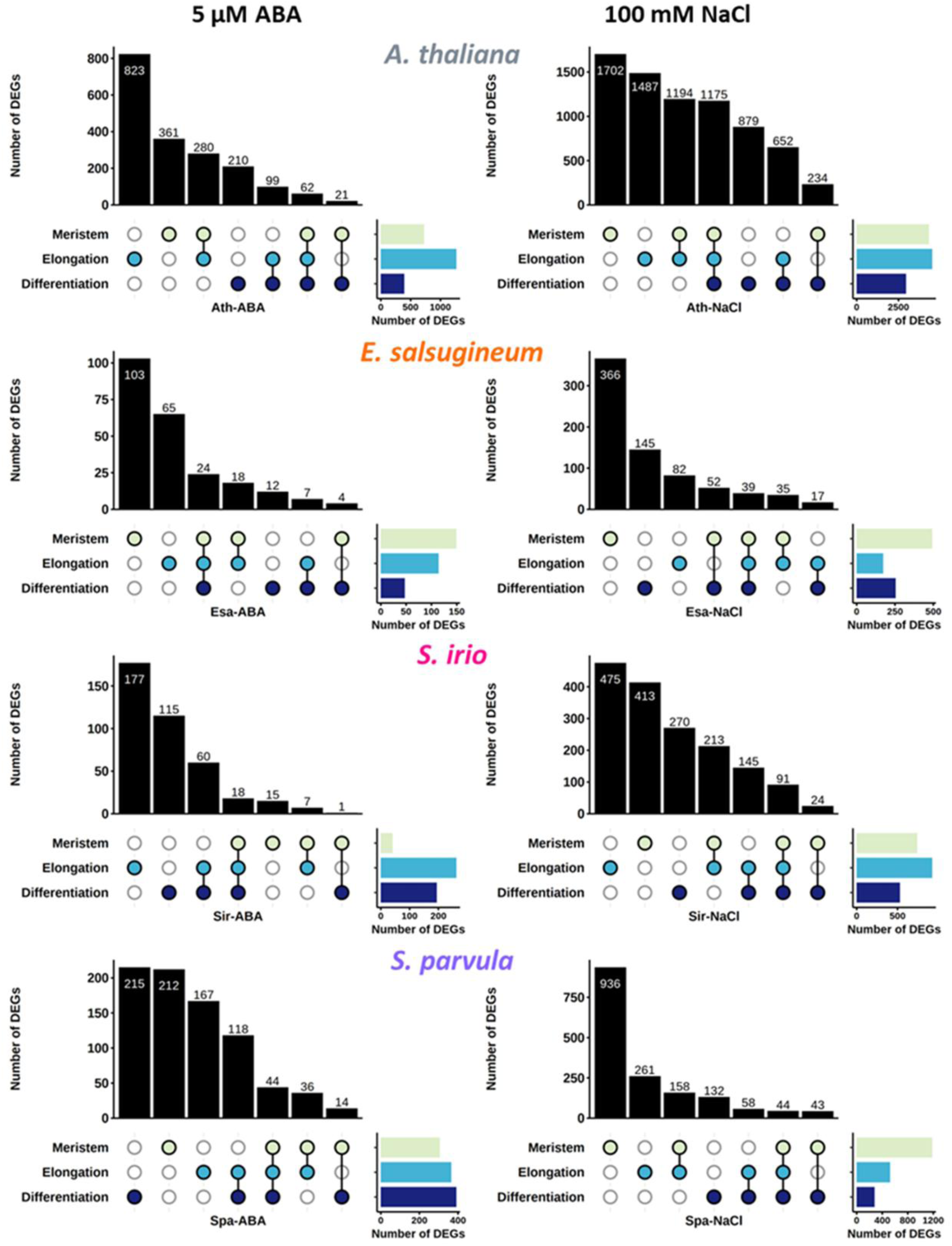
Overlap of stress responsive genes in each developmental zone from each species in response to 5 µM ABA (left) and 100 mM NaCl (right).

**Supplementary Fig. 25.**
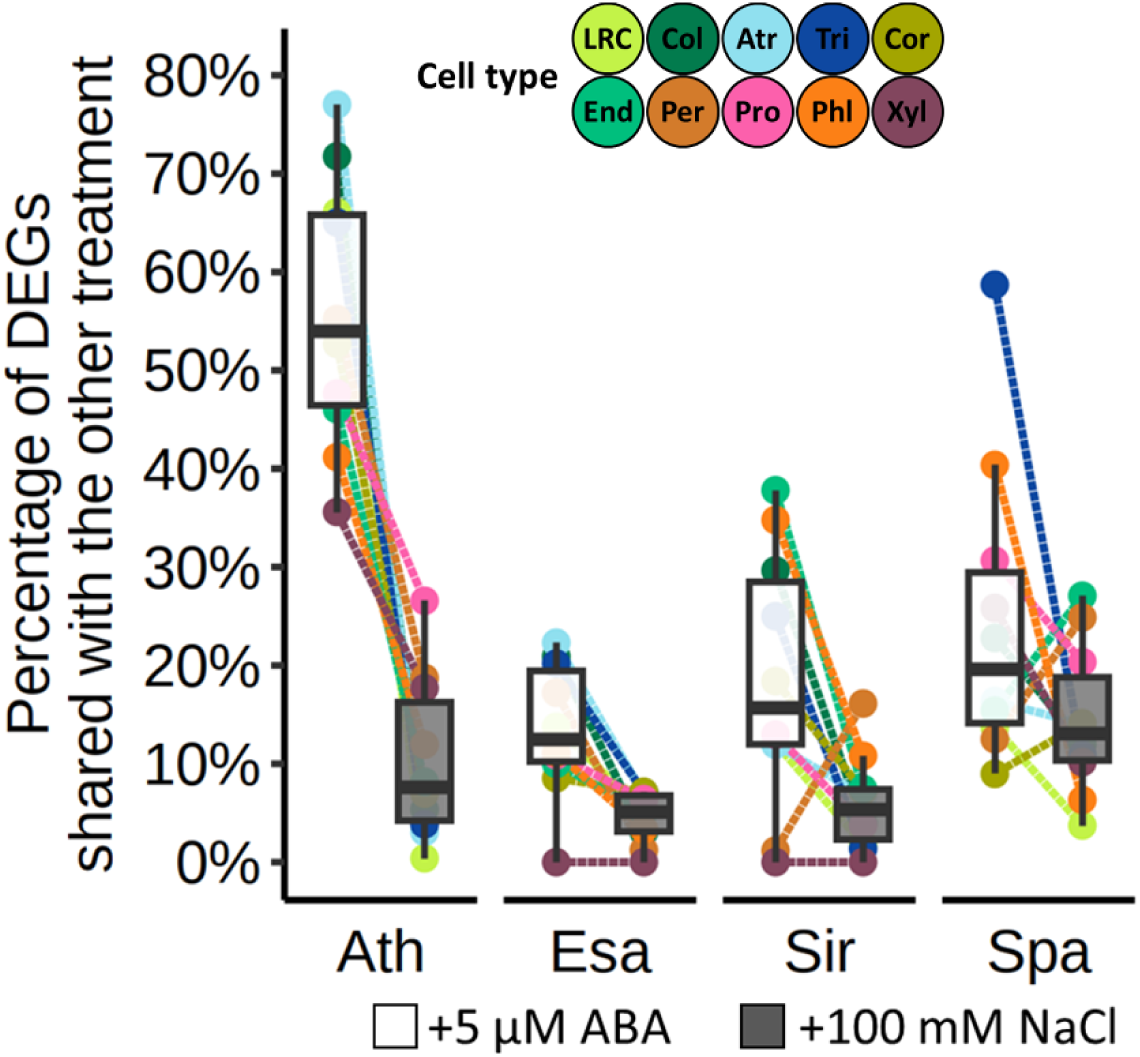
The percentage of DEGs shared between ABA and NaCl treatment. Ath, Arabidopsis; Esa, *Eutrema salsugineum*; Sir, *Sisymbrium irio*; Spa, *Schrenkiella parvula*.

**Supplementary Fig. 26.**
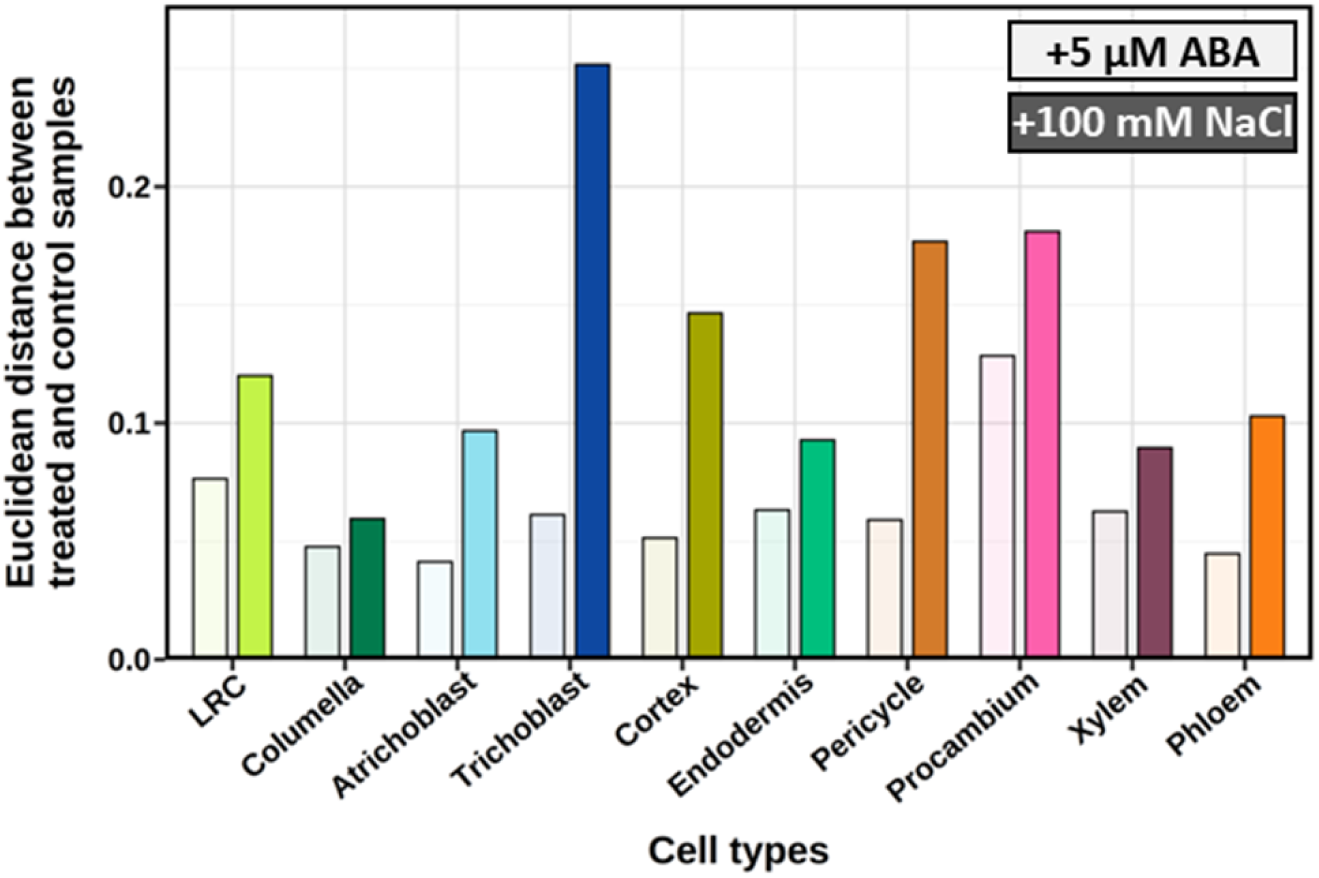
Euclidean distance between treated and control samples in Fig. 5e. The salt-treated samples were generally more distant (greater Euclidean distances) to the control samples than the ABA-treated samples.

**Supplementary Fig. 27.**
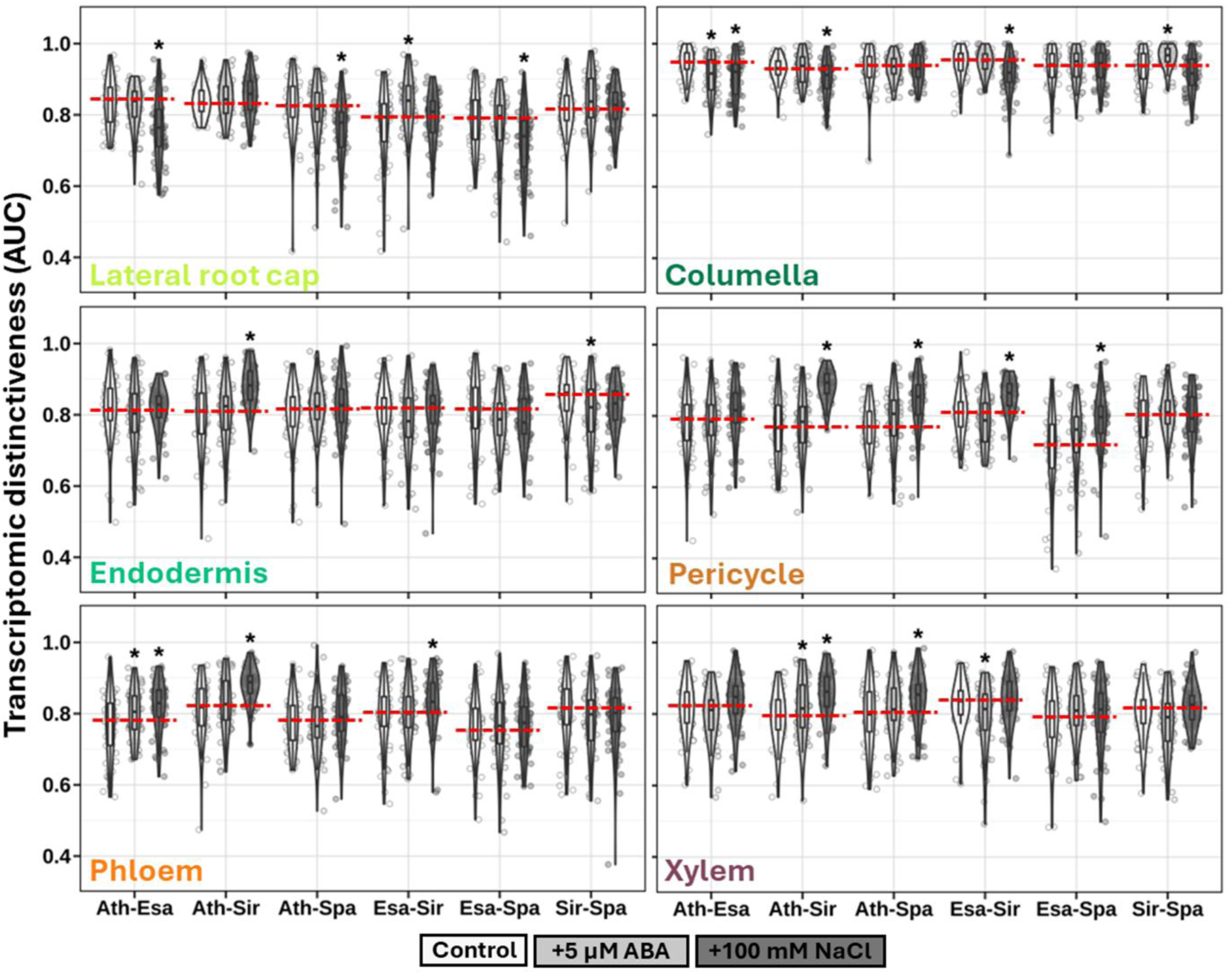
Transcriptomic distinctiveness between species pairs under different conditions. AUC here quantifies the separability between species for each cell type. AUC = 0.5 indicates random separability; AUC = 1.0 indicates perfect separability. Asterisks indicate significance after comparing to control (n = 50) with Student’s t-test at *p*-value cutoff of 0.05. The red dash lines mark the medians the transcriptomic distinctiveness under control condition. Ath, Arabidopsis; Esa, *Eutrema salsugineum*; Sir, *Sisymbrium irio;* Spa, *Schrenkiella parvula*.

**Supplementary Fig. 28.**
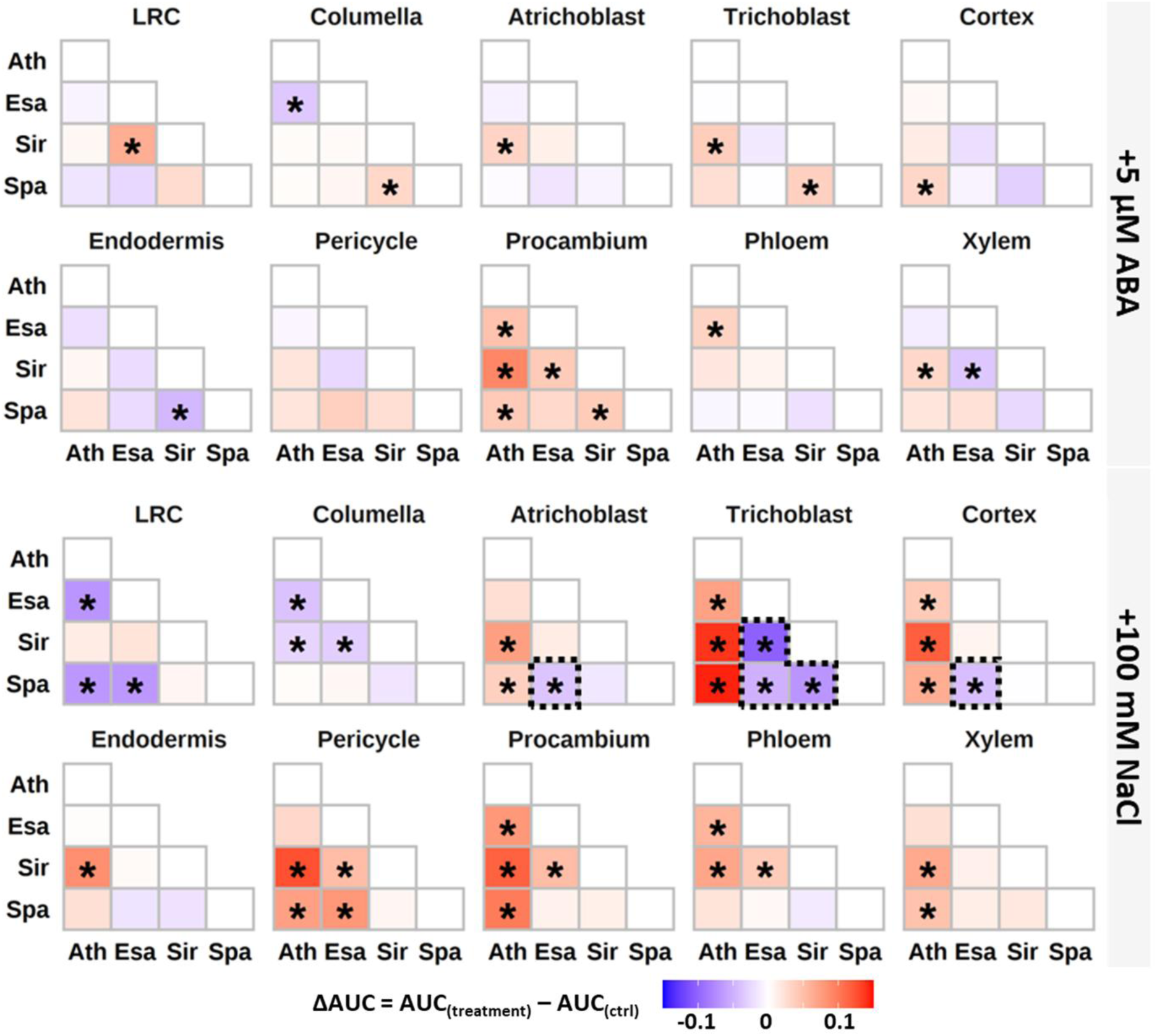
Changes in cell type transcriptome divergence between species in response to ABA (top) and NaCl (bottom). The darker colors indicate larger differences in transcriptome divergence induced by the treatment. Asterisks indicate significance after comparing to control (n = 50) with Student’s t-test at p-value cutoff of 0.05. The black boxes highlight the comparisons of interest. Ath, Arabidopsis; Esa, *Eutrema salsugineum*; Sir, *Sisymbrium irio*; Spa, *Schrenkiella parvula*.

**Supplementary Fig. 29.**
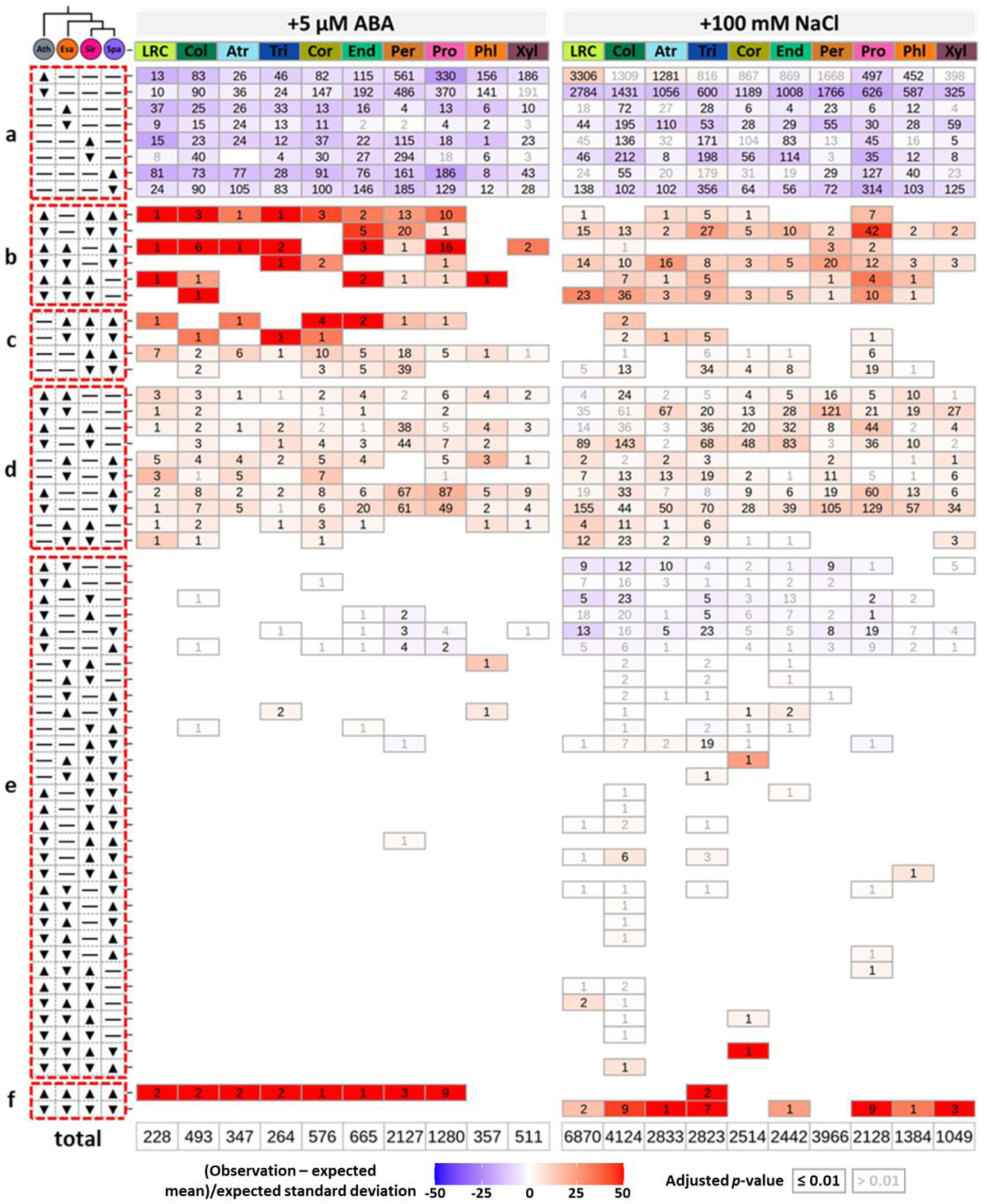
Differences between observed and bootstrapped (10,000 times) occurrences of different ortholog response patterns. The patterns were grouped according to orthologs showing a, lineage-specific gain of response; b, lineage-specific loss of response; c. synapomorphic response; d. convergent response; e, diverse response; f, conserved response. Number of orthologs for each pattern in each cell type is indicated in the corresponding tile. The differences were measured as the number of standard deviations between the overserved occurrences and the corresponding bootstrapped means and indicated with the shade of the color with darker colors corresponding to larger differences. Black and grey text represent significant and insignificant deviation of the observed occurrences from the expected distribution, respectively, at Benjamini-Hochberg-adjusted *p*-value ≤ 0.01.

**Supplementary Fig. 30.**
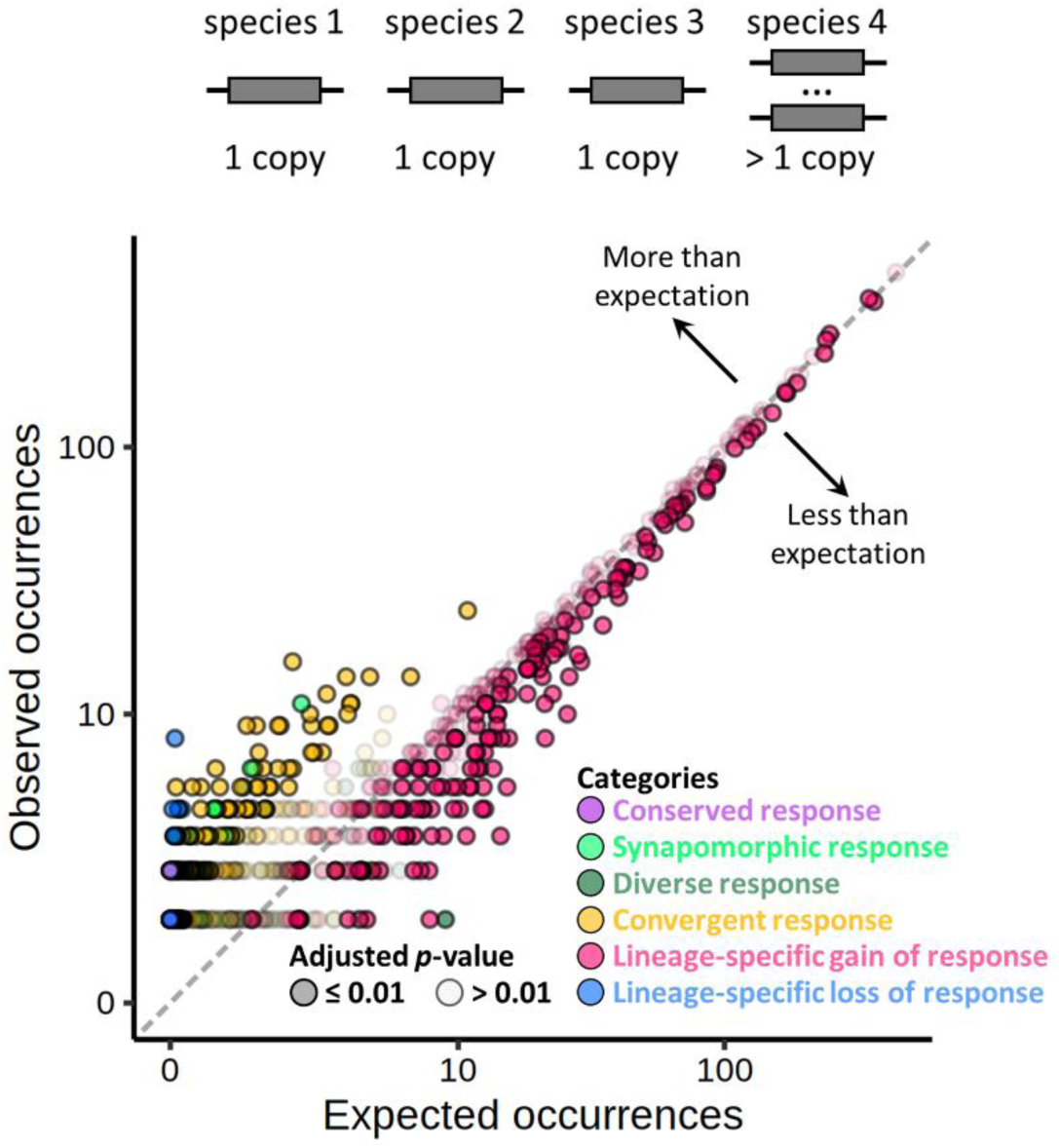
Differences between observed and bootstrapped (10,000 times) occurrences of response patterns among all ortholog groups that have been duplicated only in one species. The darker and lighter color intensities represent significant and insignificant deviation of the observed occurrences from the expected distribution, respectively, at Benjamini-Hochberg-adjusted p-value ≤ 0.01.

**Supplementary Fig. 31.**
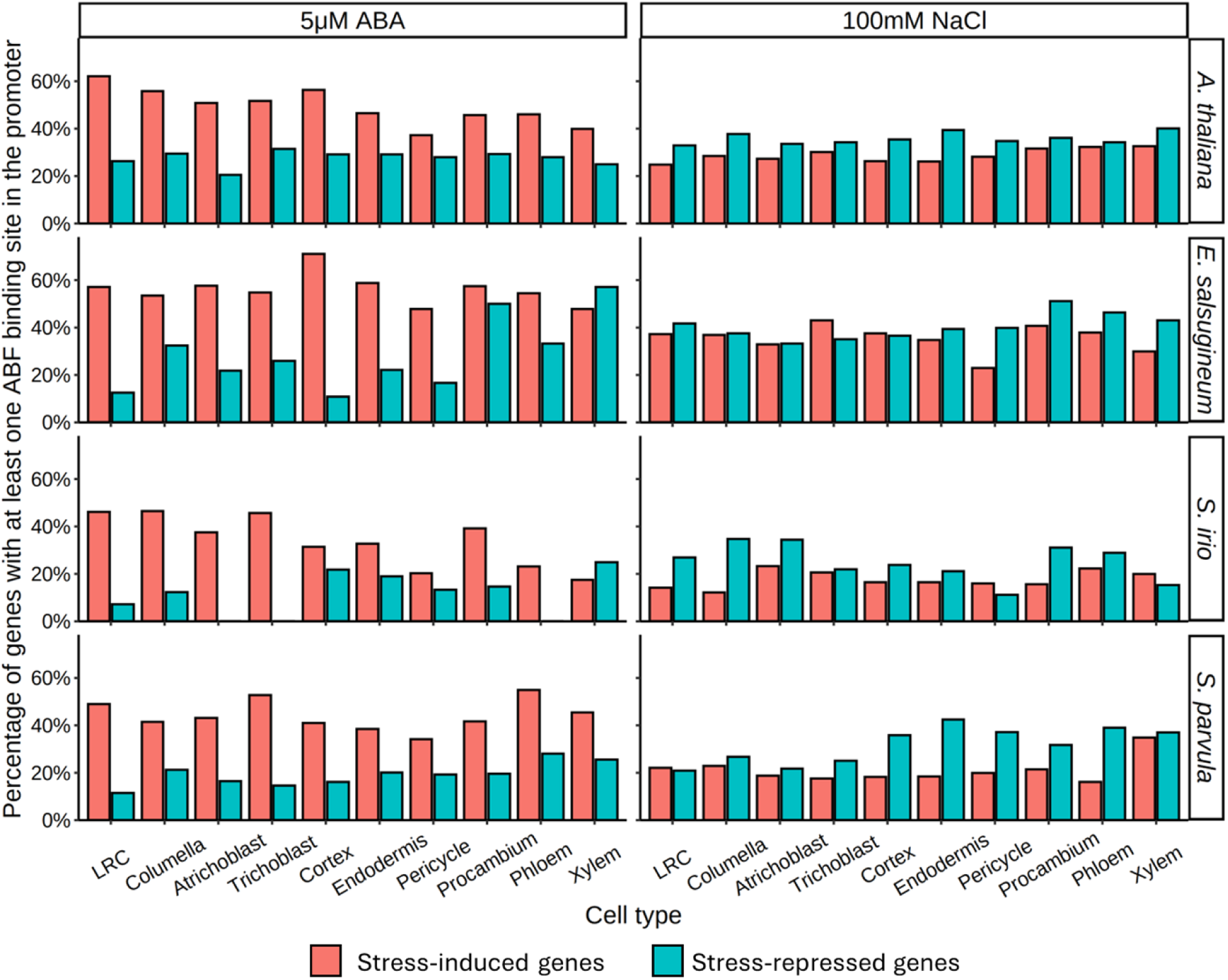
Percentage of genes with ABF binding among genes induced by 5μM ABA (left) or 100mM NaCl (right). ABF binding was defined as the presence of ABF binding events in the 2-kilobase upstream of the transcription start site using the DAP-seq data from Sun *et al.*, 2022^9^.

**Supplementary Fig. 32.**
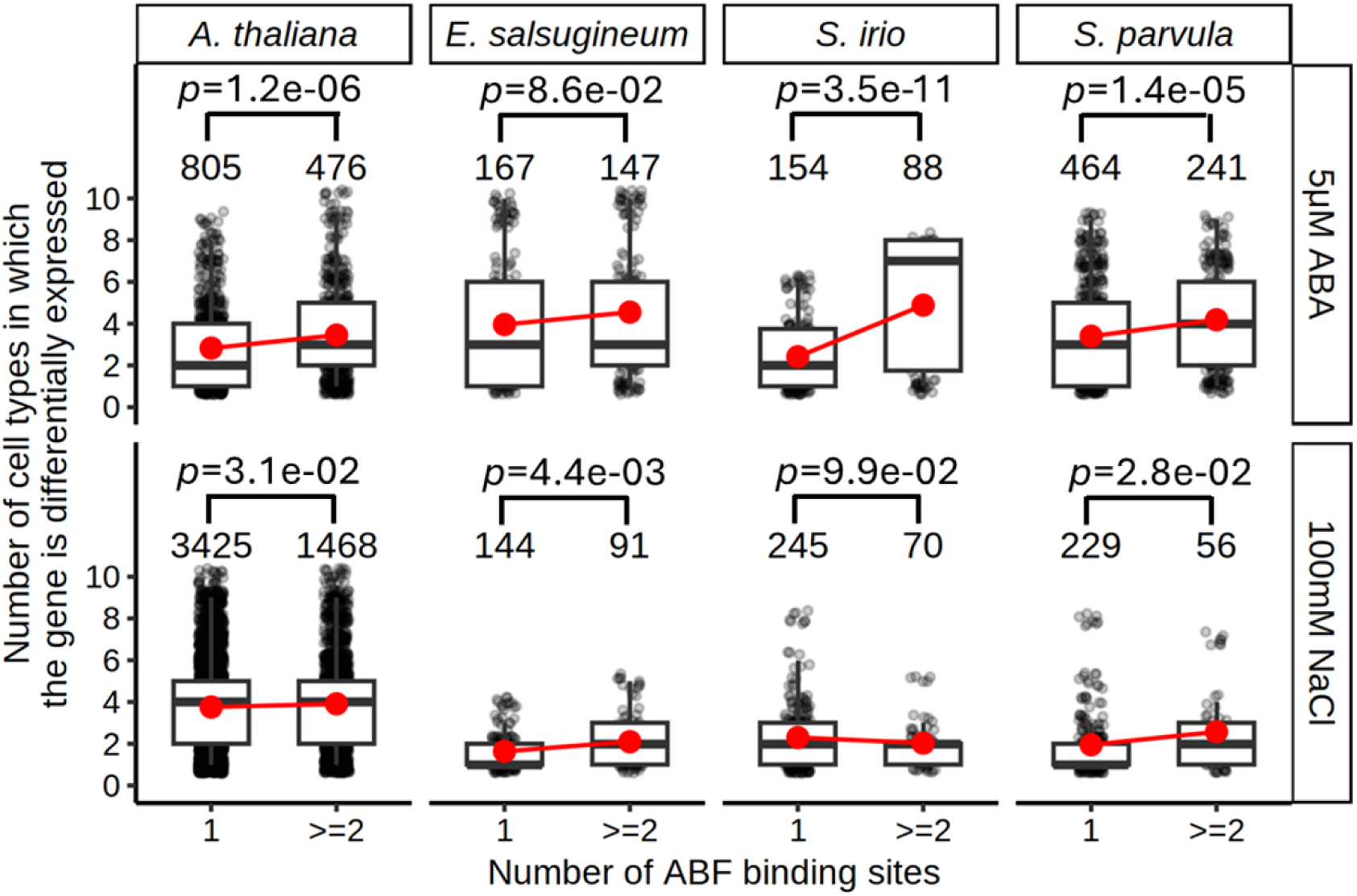
Number of responsive cell types for genes with different numbers of ABF binding sites. ABF binding was defined as the presence of ABF binding events in the 2-kilobase upstream of the transcription start site using the DAP-seq data from Sun *et al.*, 2022^9^. Significance was represented by *p*-values determined by Student’s t-test and shown on the top of each comparison. Numbers above each box indicate the number of genes included in each group.

**Supplementary Fig. 33.**
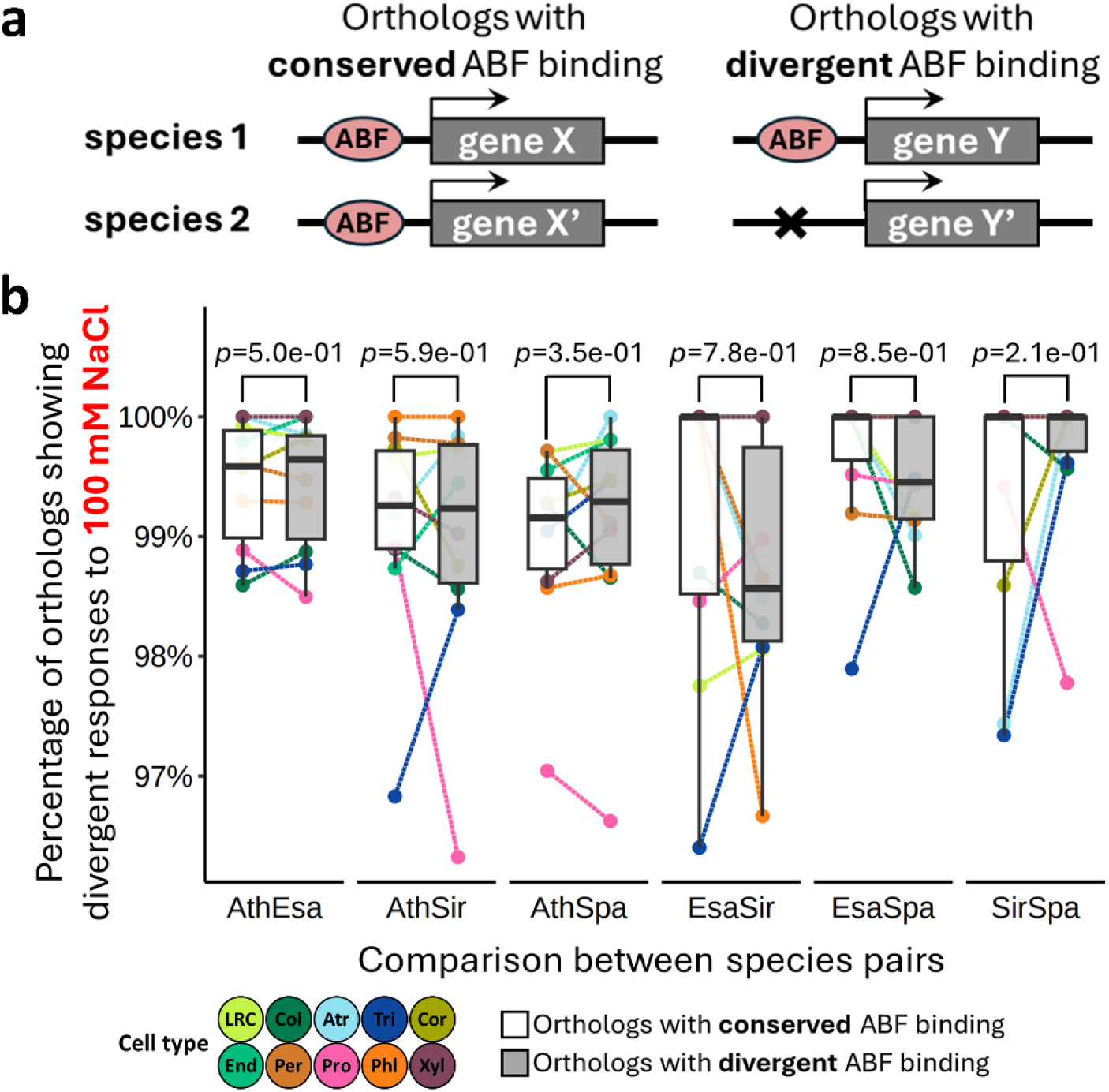
The relationship between ABF binding and stress responses of orthologs between species. a Categorization of orthologs with different ABF binding events. ABF binding was defined as the presence of ABF binding events in the 2-kilobase upstream of the transcription start site using the DAP-seq data from Sun *et al.*, 2022^9^. b Percentage of divergently regulated orthologs (in response to 100 mM NaCl) with conserved or divergent ABF binding. Significance was represented by *p*-values determined by Wilcoxon test and shown on the top of each comparison.

**Supplementary Fig. 34.**
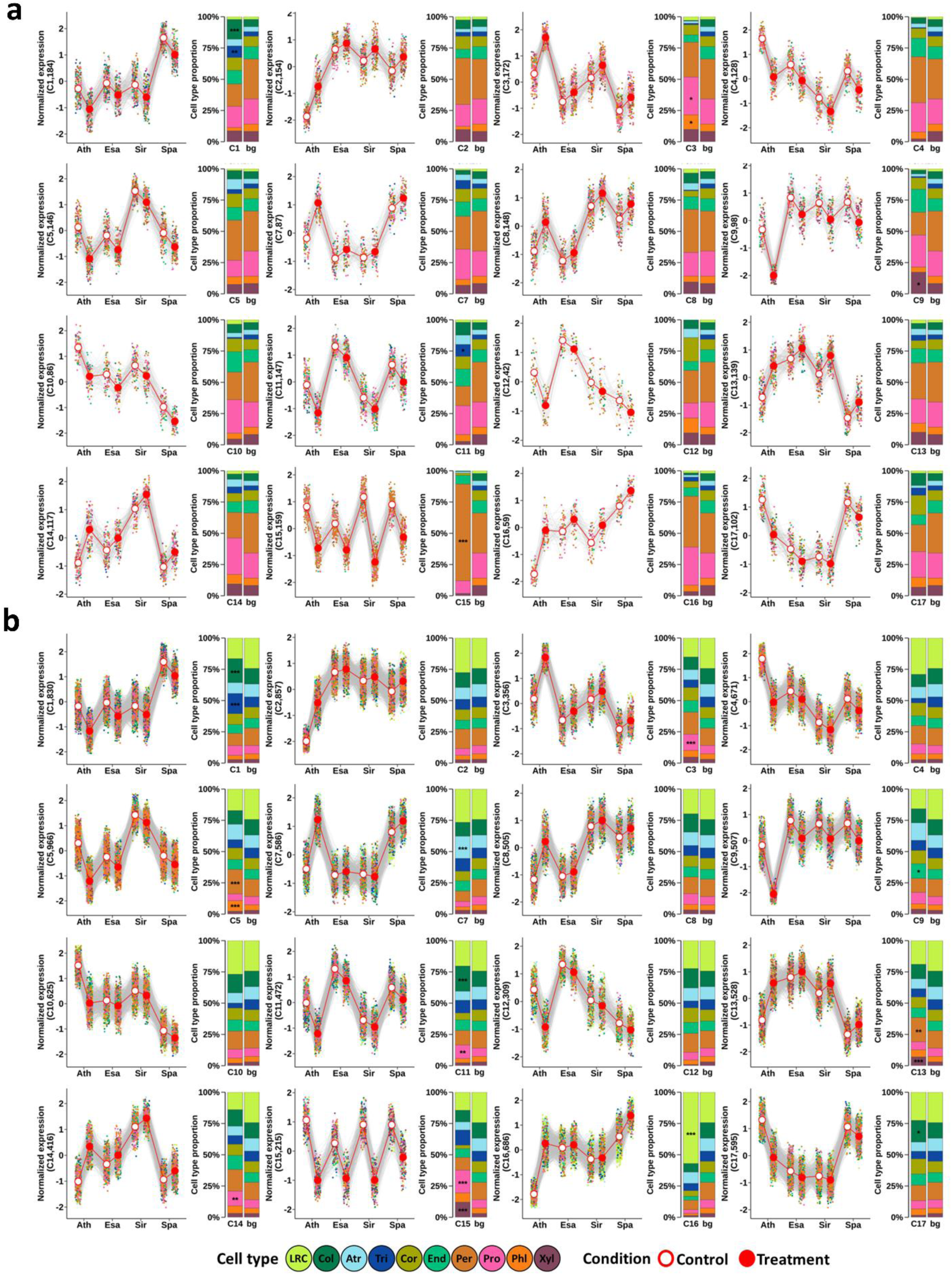
Contributions of each cell type to co-expression networks identified from stress-responsive orthologs under 5 µM ABA (a) and 100 mM NaCl (b). Cluster ID and the number of orthologs in the cluster are shown in parentheses in the Y-axis labels. Cell type proportions of the cluster were compared to background (bg) where orthologs from all clusters were averaged (see Methods for details). Asterisks represent significance determined by hypergeometric test, * *p*-value < 0.01, ** *p*-value < 0.001, *** *p*-value < 0.0001. LRC, Lateral Root Cap; Col, Columella; Atr, Atrichoblast; Tri, Trichoblast; Cor, Cortex; End, Endodermis; Per, Pericycle; Pro, Procambium; Phl, Phloem; Xyl, Xylem. Ath, Arabidopsis; Esa, *Eutrema salsugineum*; Sir, *Sisymbrium irio;* Spa, *Schrenkiella parvula*.

**Supplementary Fig. 35.**
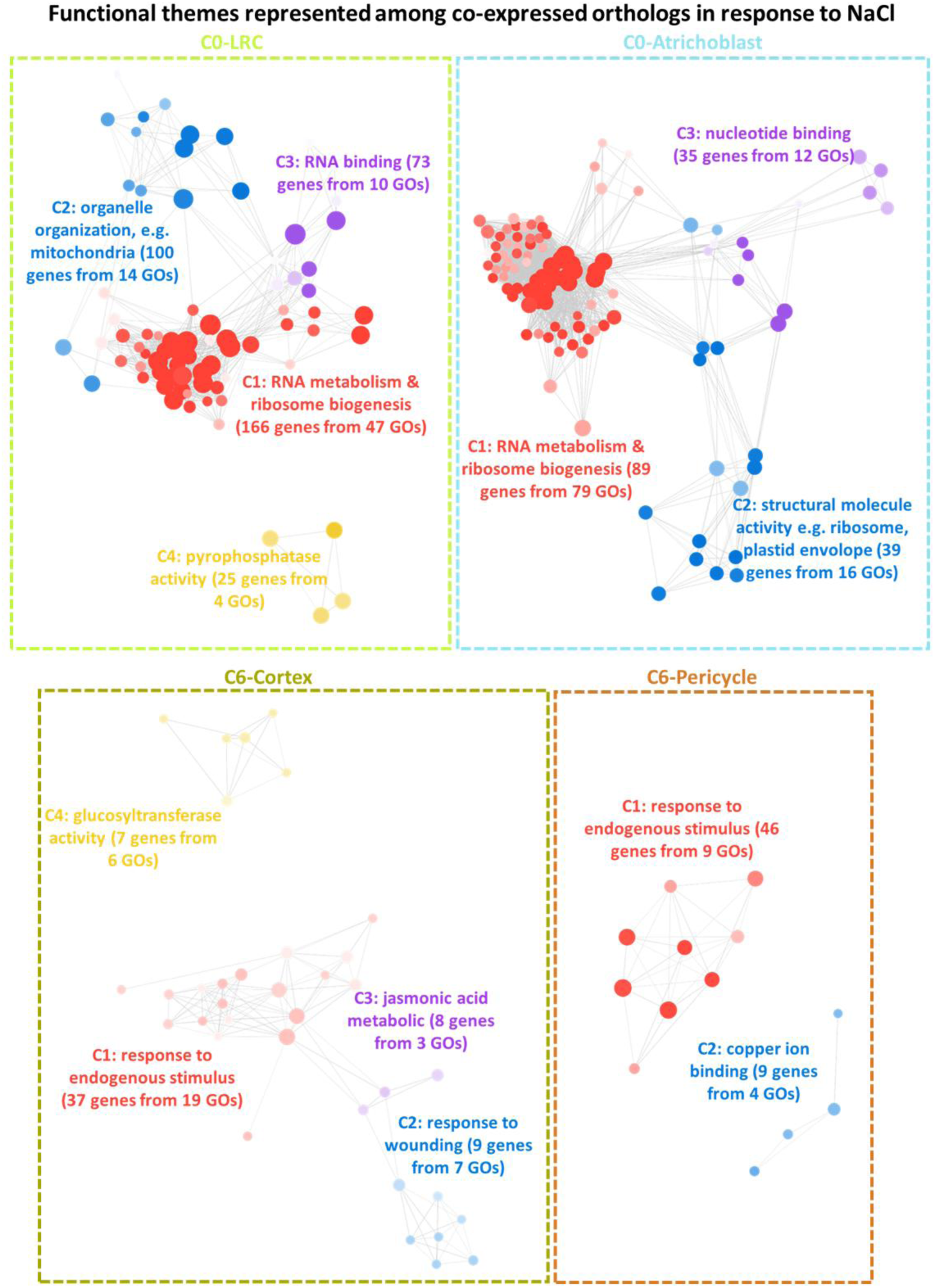
Functional representations of co-expressed orthologs in response to NaCl. GO clusters are differently colored and labelled with the representative functional term. Each node represents a gene ontology (GO) term; node size represents genes in the test set assigned to that functional term; GO terms sharing > 50% of genes are connected with edges; and the shade of each node represents the *p*-value assigned by the enrichment test at adjusted *p* values < 0.05 with darker shades indicating smaller *p*-values.

**Supplementary Fig. 36.**
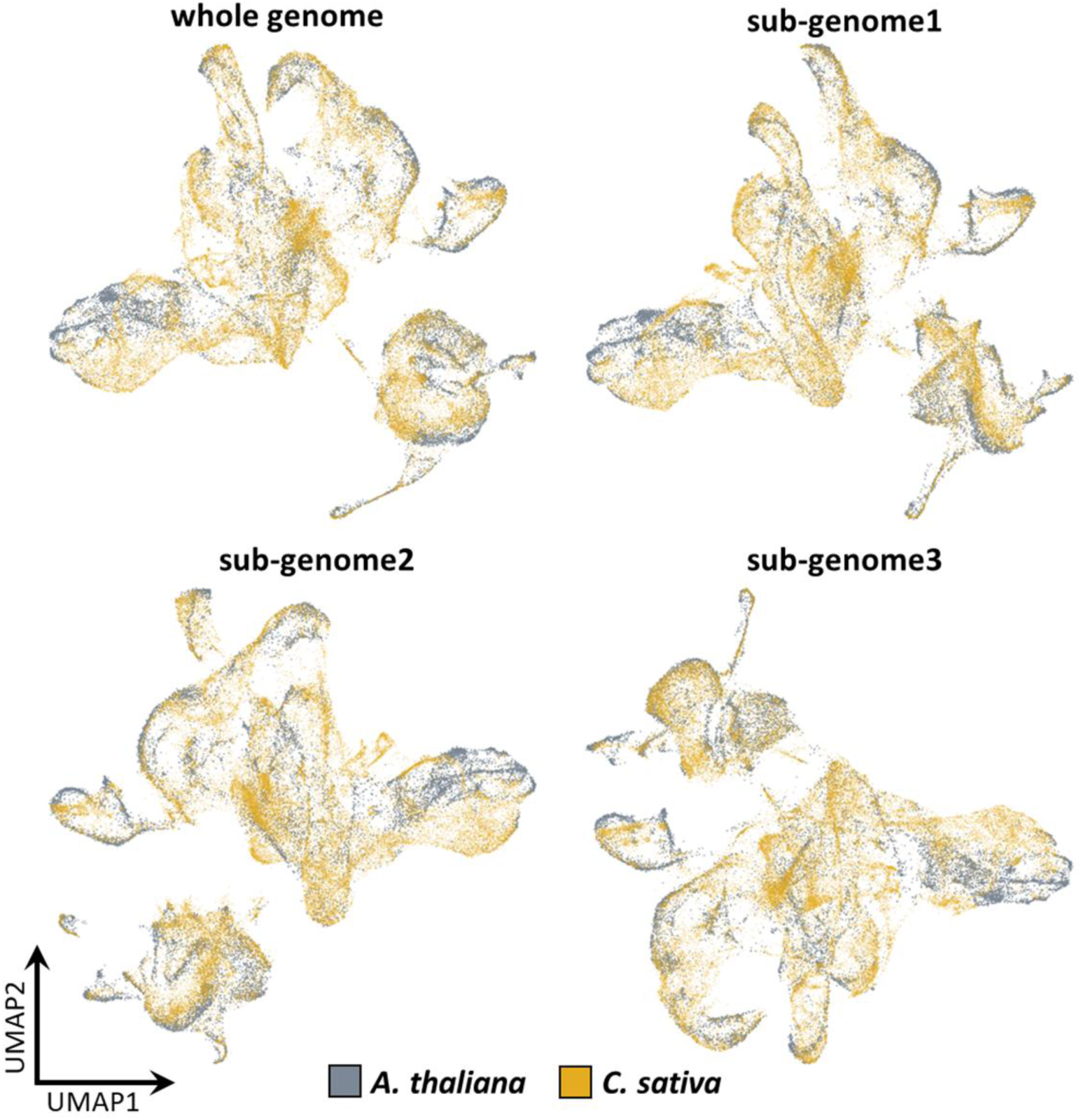
Integration between Arabidopsis and *C. sativa* using different sub-genomes in *C. sativa*.

**Supplementary Fig. 37.**
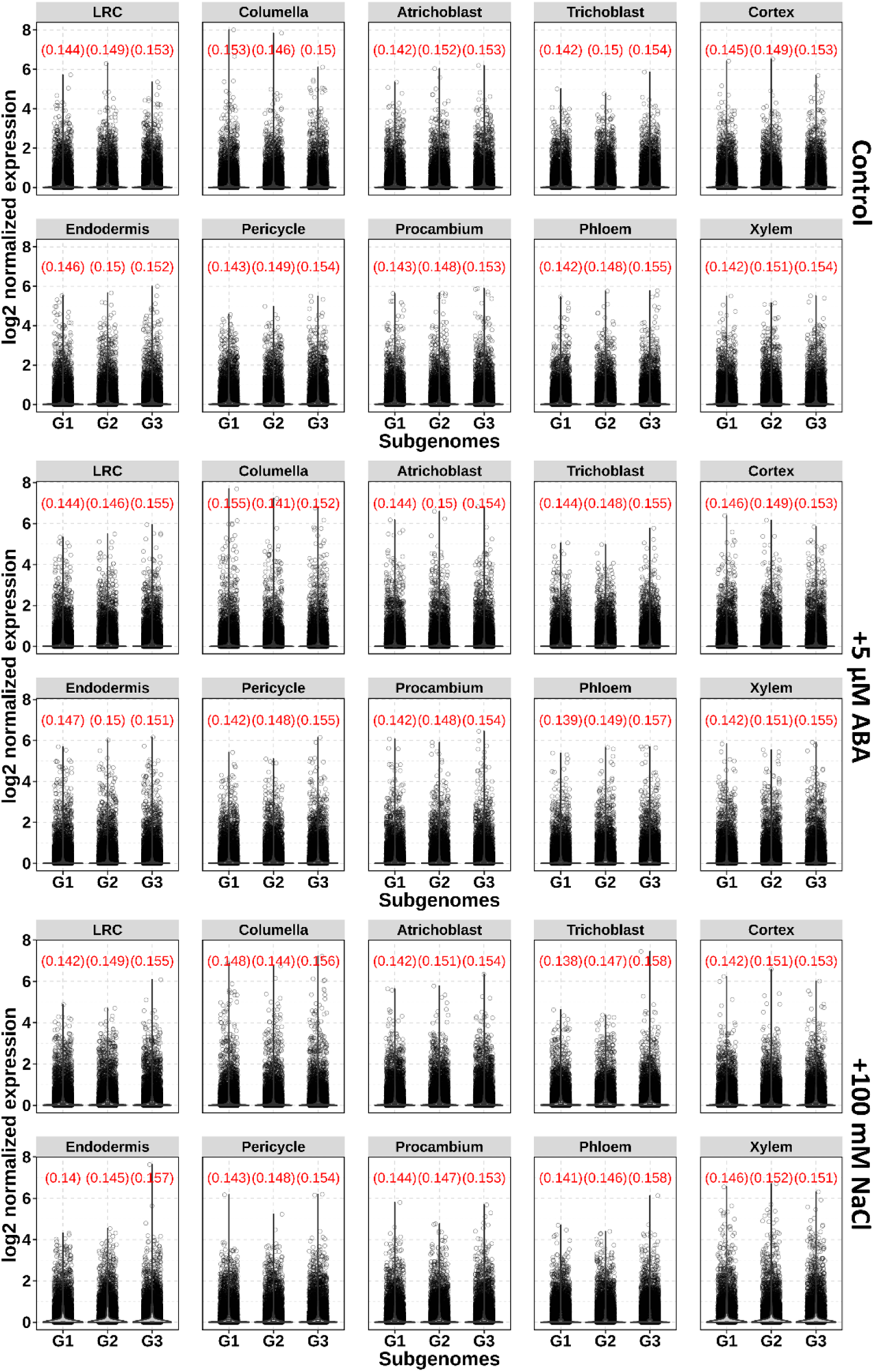
Overall transcriptome expression levels separated by the three sub-genomes for each cell type under different conditions. The number of genes considered for sub-genome G1, G2, and G3 is 21,848, 21,682, and 23,066, respectively. Numbers in red indicate the averages of transcript levels.

**Supplementary Fig. 38.**
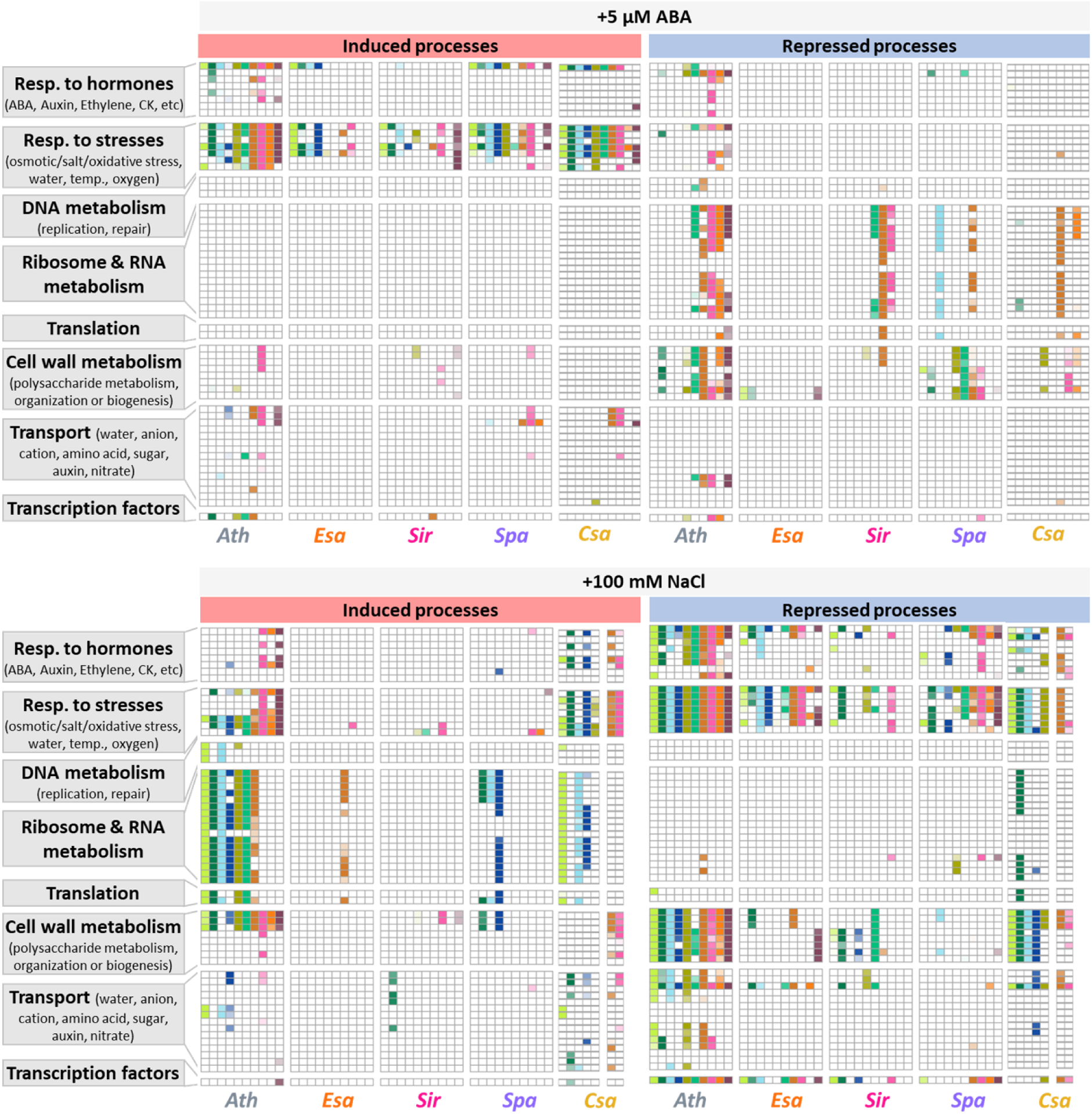
Comparison of GO functional enrichment among DEGs in response to ABA (top) and NaCl (bottom) for each cell type across species. The enrichment in *C. sativa* was performed on DEGs combined from all sub-genomes. Endodermis, phloem and xylem cells in *C. sativa* were removed from the comparison for NaCl treatment due to insufficient number of cells (< 50 cells). The detailed list of GO terms is present in Supplementary Data 6. The shade represents the *p*-value assigned by the enrichment test at adjusted *p*-value < 0.05 with darker shades indicating smaller *p*-values.

**Supplementary Fig. 39.**
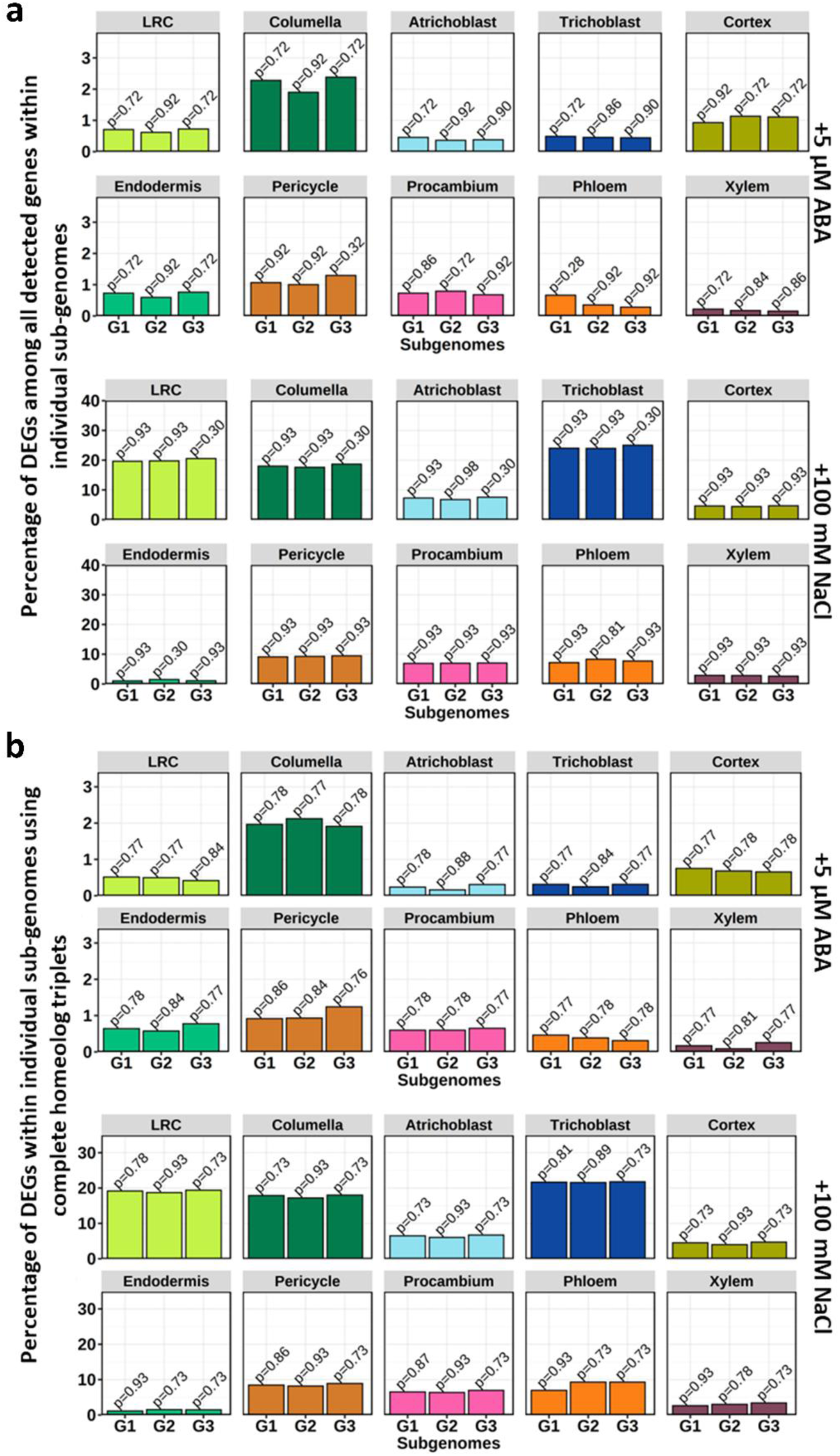
Percentage of DEGs in each sub-genome of *C. sativa*. The percentage for each genome was calculated considering all detected genes within individual sub-genomes (a) and only the complete homeolog triplets (b). Numbers on top of the bars represent adjusted *p*-values from hypergeometric test.

